# The cellular and genetic basis of inflorescence divergence between maize and teosinte

**DOI:** 10.64898/2026.08.10.743805

**Authors:** Yuebin Wang, Ruijie Mao, Yu Liu, Xing Guo, Nan Li, Qian Zhang, Manjun Cai, Pingli Xie, Yi Wang, Yun Luo, Qian Ding, Shenshen Wu, En Luo, Ling Ma, Zheng Luo, Tong Wei, Huan Liu, Mingqiu Dai, Fazhan Qiu, Yingjie Xiao, Xiaohong Yang, David Jackson, Zuxin Zhang, Jianbing Yan, Jeffrey Ross-Ibarra, Lei Liu, Ning Yang

## Abstract

The domestication of maize from teosinte involved dramatic remodeling of the ear, yet the cellular and genetic bases of this transformation remain unclear. Here, we generate a single-nucleus and spatial transcriptome atlas of developing maize and teosinte ears. Comparative analysis reveals divergence in cob-associated cell types, with enhanced cytokinin signaling and reduced growth-inhibitory signals collectively driving cob thickening and enlargement in maize. We further demonstrate that domestication expanded the spatial expression domain of key transcription factors in maize meristem cells, enhancing the potential for increasing kernel number. Additionally, we verified a major domestication gene, *ZmSPD1*, in which two nonsynonymous SNPs differentiate maize from teosinte and alter jasmonic acid (JA) levels in the ear, thereby suppressing spikelet abortion to effectively double kernel production. These findings provide a cell-resolved mechanistic framework for how cob architecture and kernel number were shaped during maize domestication, offering new insights into the formation of key agronomic traits.

## Introduction

The domestication of crop plants through sustained cycles of cultivation and selection by early farmers represents a pivotal transformation in human history, driving profound changes in the diversity, morphology, and physiology of key food species. Maize (*Zea mays* subsp*. mays*) is one of the most agriculturally important crops globally, and represents one of the most striking morphological transformations associated with domestication. Archaeological (*1*) and population genomic studies (*2, 3*) have established that early farmers domesticated maize in Mexico starting 9,000 years ago, and identified that at least two wild teosintes (*Zea mays* ssp. *parviglumis* and *Zea mays* ssp. *mexicana*) contributed to modern maize (*4*). Despite its relatively recent origin, the maize ear underwent a radical transformation during domestication (*5*). Teosinte ears have two rows of kernels, where each spikelet pair meristem initially produces two spikelets, but one of them aborts, leaving only a single functional spikelet at each node that produces fewer than a dozen kernels, covered by hardened glumes that disarticulate at maturity (*6*). The maize ear, in contrast, has evolved into a massive structure with at least four rows, both spikelets in a pair develop into fertile florets, effectively doubling the kernel production per node and yielding as many as 1,000 kernels on a large, soft cob (rachis) that prevents seed dispersal.

While population and quantitative genetic research has identified hundreds of genes selected during the domestication of maize (*7–9*), these genome-wide signals of selection contrast with classical genetic evidence indicating that a relatively small number of loci exert disproportionate effects on the major morphological differences between maize and teosinte, as originally demonstrated in early genetic studies (*5, 10*). Reconciling these two perspectives requires both the identification of major causal loci and a mechanistic understanding of how genes act to generate large-scale phenotypic change. Several loci underpinning variation in ear morphology have been identified, including *Kernel Row Number2 (KRN2)* (*9*), and *Kernel Row Number4 (KRN4)* (*11*) for increased kernel row number, and *TGA1 (Teosinte Glume Architecture1)* (*12*) for exposed kernels. Nonetheless, the genetic basis of another key trait – the lack of aborted spikelets in maize, which results in doubling the number of kernel rows compared to teosinte – remains poorly understood. Moreover, the cellular mechanisms through which these genetic loci orchestrate ear architecture divergence across specific cell types are also mostly unknown. As key agronomic traits in maize are largely governed by transcriptional variation (*13*), we therefore present here a cell-type-resolved spatiotemporal transcriptome atlas of developing maize and teosinte ears, illuminating how transcriptional programs were reconfigured during domestication. Our atlas links genetic variation and selection signals to cell-type-specific regulatory changes that ultimately shaped ear morphology and transformed a modest wild grass into one of humanity’s most productive crops.

## Results

### Construction of single-nucleus spatiotemporal transcriptome atlas of maize and teosinte ears

To understand the genetic and molecular mechanisms underpinning the morphological divergence between maize and teosinte ears, we sampled a series of maize ear primordia at stages where key developmental decisions are made (2 mm, 4 mm and 6 mm) (Fig. 1, A and B). For teosinte, we pooled samples across the 1–3 mm range to comprehensively capture the critical period of spikelet pair meristem patterning (Fig. 1C). We performed single-nucleus transcriptome sequencing (snRNA-seq) and retained 104,284 maize and 29,185 teosinte high-quality nuclei for downstream analyses (see Methods; detailed in table S1). Maize cell types were annotated using STRIDE (*14*) based on a published maize spatial transcriptome atlas (fig. S1, table S2) (*15*), successfully identifying most cell populations including the inflorescence meristem, meristem epidermis and vasculature. These annotations were also further validated using established marker genes (fig. S2 and S3, table S3).

**Fig. 1.**
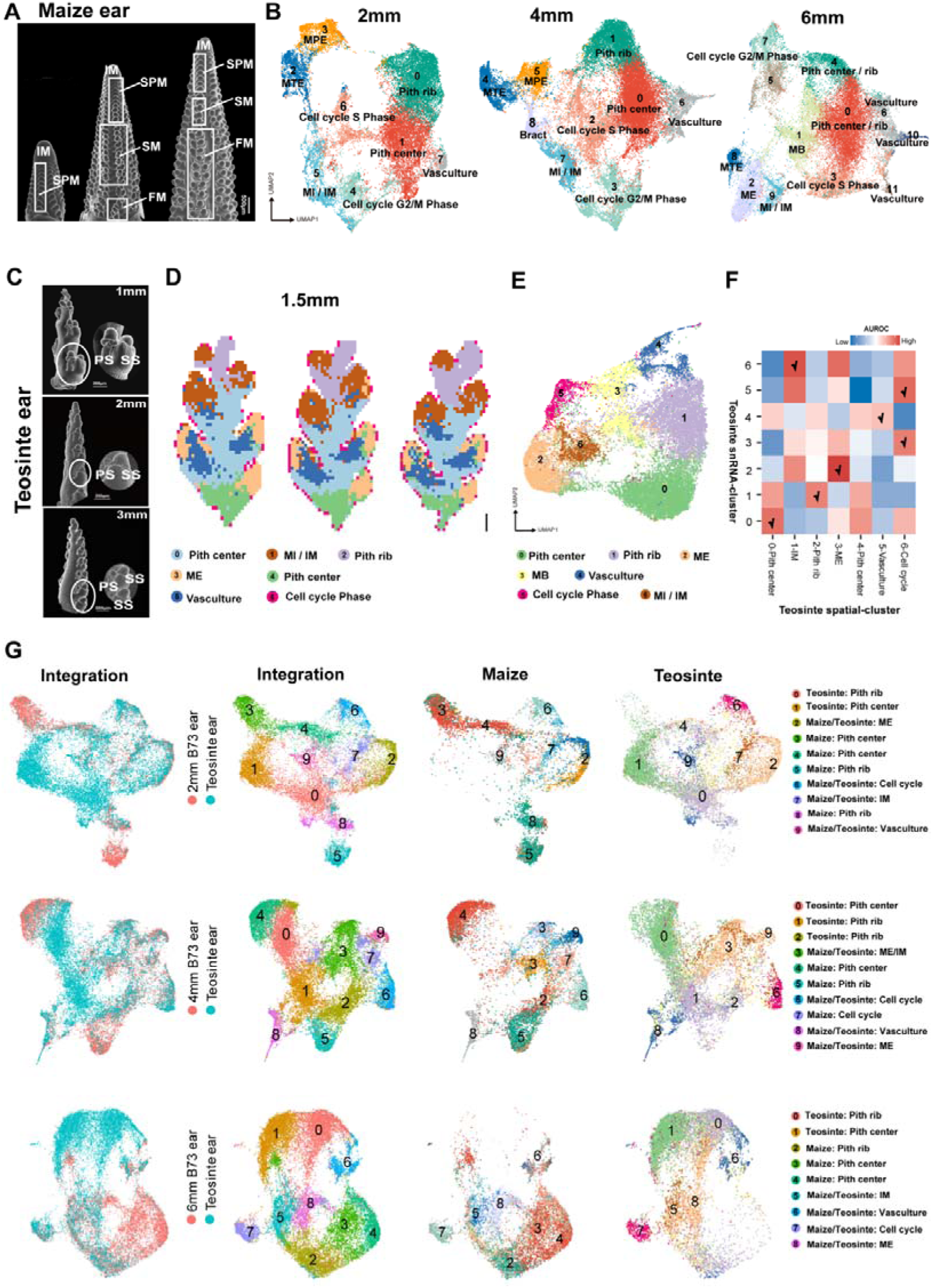
Single-nucleus and spatial transcriptomic atlases of maize and teosinte ear development. **(A)** Representative morphology of maize ears at three developmental stages: early (2 mm), mid (4 mm), and late (6 mm). **(B)** Cell clusters of maize ear visualized by an integrated UMAP plot from three developmental stages. Inflorescence meristem (IM); Meristem base (MB); Meristem internal (MI); Meristem tip epidermis (MTE); Meristem periphery epidermis (MPE); Meristem epidermis (ME). **(C)** Teosinte ear at 1–3 mm, highlighting the critical window for IM structure and pedicellate spikelet abortion. **(D)** Spatial distribution of cell types across three teosinte ear sections. Scale bar = 0.1mm **(E)** Seven cell clusters of teosinte ear visualized by an integrated UMAP plot in two replicates. **(F)** MetaNeighbor analysis identifying corresponding cell types between teosinte snRNA-seq and spatial transcriptomic datasets. Labels indicate cell types with highest similarity (AUROC > 0.7). **(G)** Cross-species integration of maize and teosinte transcriptomes. Integrated UMAP showing the combined distribution of nuclei from both species, colored by cluster identity. UMAP of maize nuclei (subsetted from the integrated data), with cell types annotated and colored according to the reference in panel (B). UMAP of teosinte nuclei (subsetted from the integrated data), with cell types annotated and colored according to the reference in panel (E).

To facilitate annotation of teosinte snRNA-seq data, we generated a high-resolution spatial transcriptomic atlas of 1.5 mm teosinte ears (within the sampling window for snRNA-seq), retaining more than 7,400 high-quality bin50s (spatial transcriptome data units, details see Methods) across 3 sections, with an average of 23,322 expressed genes per section (table S4). Next, bin50s from adjacent sections were integrated and subjected to unsupervised clustering following imputation (see Methods). Uniform Manifold Approximation and Projection (UMAP) visualization revealed seven distinct clusters with clear spatial patterns (Fig. 1D and fig. S4), corresponding to spikelet meristems, inflorescence meristem and pith, displaying a spatial organization closely resembling that of maize. Comparison of the spatially annotated maize and teosinte transcriptomes revealed that cell populations occupying analogous spatial domains had greater transcriptomic similarity between the two species (fig. S5 and table S5). We then projected the teosinte snRNA-seq data onto its spatial atlas, assigning seven corresponding cell types (Fig. 1E and F). To further validate the teosinte cell type annotations, we performed in situ hybridization for selected marker genes, confirming both the spatial localization and identities of the annotated teosinte cell types (fig. S6 and table S6). Together, these results demonstrate that both key cell type identity and spatial organization are largely conserved between maize and teosinte ears, indicating that their morphological divergence occurs within a shared cellular framework.

We then independently integrated snRNA-seq data from maize and teosinte across maize developmental stages. This strategy ensured that transcriptional similarities and differences were assessed separately for each maize developmental stage, thereby maximizing the resolution to capture stage-specific divergence between maize and teosinte. Our analysis revealed strong conservation for many spikelet-associated cell types, such as meristem epidermis, meristem internal and inflorescence meristem cells (Fig. 1G). In contrast, cob-associated cell types (pith rib, pith center and vasculature) showed only partial similarity between teosinte and maize (Fig. 1G and fig. S7), suggesting that divergence in cob-associated developmental programs underlies the dramatic differences between slender, reduced teosinte cobs and the enlarged, thickened maize cobs.

### Temporal and cellular regulatory divergence reshaped ear morphology between maize and teosinte

To delineate transcriptional changes underlying cob evolution, we compared cob-associated cell types (pith rib/center and vasculature) from maize and teosinte ears (fig. S8A). Differentially expressed genes (DEGs) in these cell types were then identified based on >2-fold changes in expression range (the proportion of cells expressing the gene in that cluster) or average expression levels per cell (fig. S8B and table S7). In each stage-specific comparison (2 mm, 4 mm, and 6 mm maize versus teosinte, respectively), DEGs showed distinct functional enrichments. DEGs with expanded expression range in maize pith cells were enriched in cytokinin signaling pathways at the 2 mm and 4 mm stages, whereas those at the 6 mm stage were mainly associated with growth- and development-related processes (Fig. 2A and fig. S8C). In contrast, teosinte pith cells show a limited cytokinin expression range (less than half the range in maize; fig. S9), suggesting reduced proliferative activity in teosinte cobs. Although most cytokinin-associated genes showed no signs of selection (*9*), several domestication-related transcription factors (TFs), such as *TSH4 (Tassel Sheath4)* (*16*), *UB2 (Unbranched2)* (*17*), and *UB3 (Unbranched3)* (*18*), showed increased expression range in maize pith cells (fig. S9A and B). These TFs might indirectly influence cob expansion processes, as evidenced by the role of *UB3* in regulating cytokinin levels (*19*). Our DEG comparison also identified *bb2* (*big brother 2*), a negative regulator of organ growth and size (*20*) located in a cob-diameter QTL (*21*) that shows evidence of selection (*9*) and expression changes. Our snRNA-seq data revealed that *bb2* was broadly expressed in teosinte pith cells (more than double the range in maize) (fig. S9D and table S7). To further explore regulatory interactions, we constructed a co-expression network centered on *bb2* in teosinte pith cells, which identified *CNR6* (*Cell Number Regulator 6*) as a strong co-expression partner (Fig. 2B). Similar to *bb2*, *CNR6* also showed broader expression in teosinte pith cells than in maize (fig. S9E). The *CNR* gene family negatively regulates cell number and organ size in maize (*22*), suggesting that *CNR6* may function together with *bb2* to constrain cob growth in teosinte. Taken together, these findings indicate that maize cob enlargement occurred through at least two complementary mechanisms: upregulating cytokinin-related genes to drive cell proliferation, and downregulating growth repressors such as *bb2* and *CNR6* to relieve constraints on cell proliferation and release organ growth potential.

**Fig. 2.**
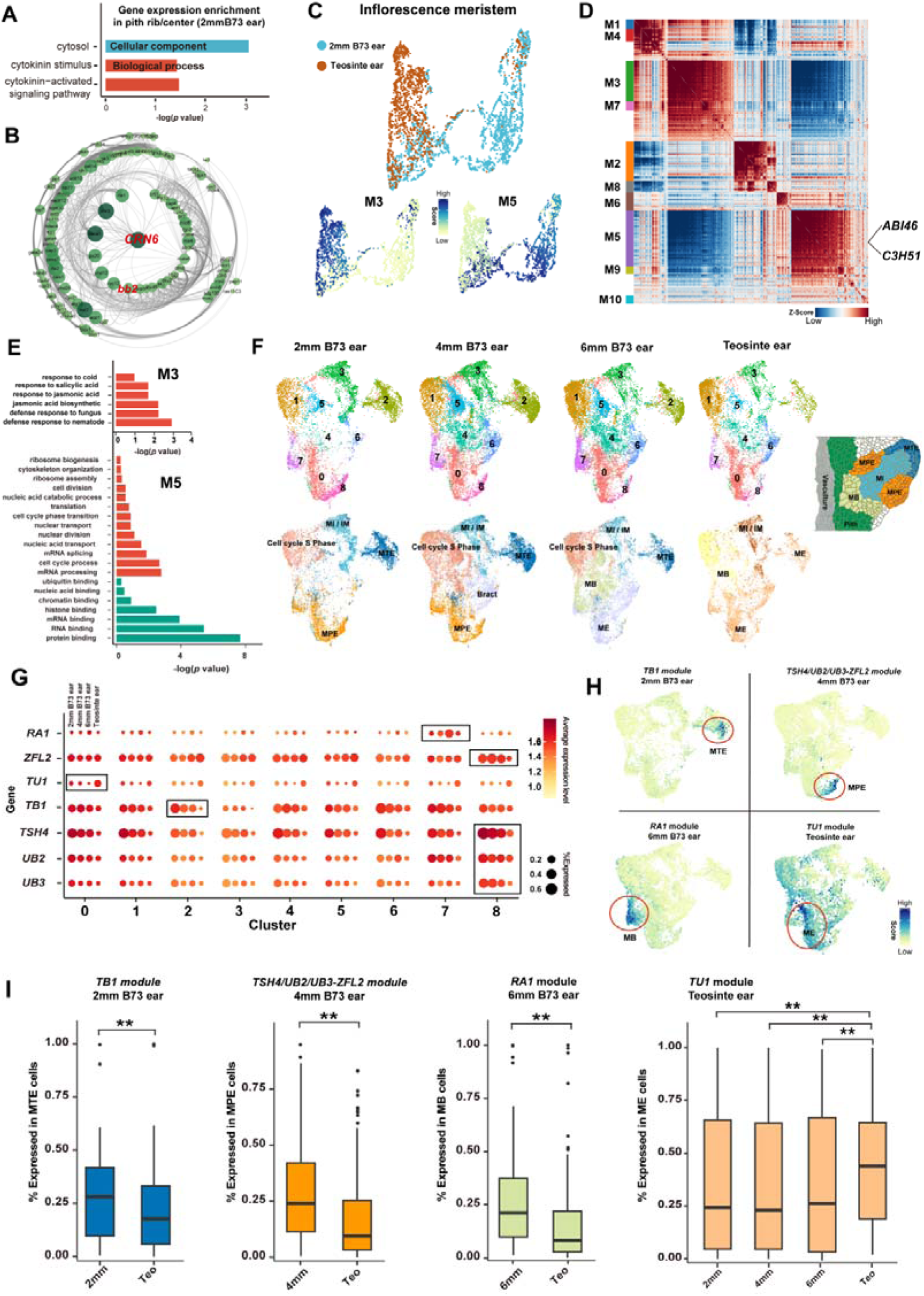
Comparative single-nucleus transcriptomic analysis of meristem cell populations in maize and teosinte ears. **(A)** Bar graph showing GO enrichment of highly expressed genes in 2mm maize pith cells. **(B)** Gene co-expression network constructed containing *bb2* and *CNR6*. Node size represents the degree of connectivity, The shade of the line color represents the weight of the edge between two nodes. **(C)** UMAP visualization and clustering of IM cells in maize and teosinte. (**D**) Hotspot analysis and identification of gene modules within isolated IM cells. Each module highlighted by a distinct color bar, on the basis of pairwise local correlation. (**E**) GO enrichment analysis of the maize-specific and teosinte-specific IM gene module. **(F)** UMAP visualization and clustering of spikelet meristem cell types in maize and teosinte. Sketches of longitudinal section of a spikelet meristem are based on (*33*). **(G)** Comparative %expressed (the proportion of cells expressing the gene in that cluster), and average levels of key domestication TFs across spikelet meristem cell subgroups show in Fig. 2F, and developmental stages. The black box indicates the cell cluster showing the most significant difference in gene expression between maize and teosinte. The four dots corresponding to each cluster represent the expression levels of the gene in the 2mm, 4mm, 6mm B73, and teosinte ear within that cluster; the color intensity of the dots indicates the gene average expression level, while the dot size indicates the gene %expressed in the cluster. **(H)** Spatiotemporal visualization of gene modules including key TFs in specific cell types and development stages. Gene module with higher hotspot scores indicate stronger co-expression patterns. **(I)** %Expressed (Expression range) are shown for all genes within the whole gene module across specific cell types and developmental stages: 2 mm B73 ear (2 mm), 4 mm B73 ear (4 mm), 6 mm B73 ear (6 mm), and teosinte ear (Teo).

Cob expansion and thickening underpin the increase in kernel number in maize, which is primarily determined by kernel row number and ear length. The developmental potential of ear length is strongly influenced by inflorescence meristem (IM) activity (*23*). We therefore independently extracted and compared IM cell clusters from our maize and teosinte snRNA-seq. Compared with the teosinte IMs, maize IMs had a greater number of highly expressed genes associated with chromatin modifications, G1/S cell-cycle regulation, and ribosome biogenesis (fig. S10), suggesting enhanced transcriptional activity and a more active proliferative state during maize IM development. Using Hotspot (*24*), we further identified maize and teosinte-specific gene modules, (Fig. 2C, D and fig. S11). The maize-specific module was significantly enriched for cell division-related GO terms, whereas the teosinte-specific module showed enrichment for stress-responsive pathways (Fig. 2E and fig. S11), suggesting an earlier transition of teosinte IMs away from an active proliferative state, resulting in reduced ear length. Although their roles in IM have not been experimentally validated, we identified other transcription factors such as *ABI3-VP1-transcription factor 46* (*ABI46)* and *C3H-transcription factor 351* (*C3H51)* in the maize module, which show both expanded expression range across developmental stages (Fig. 2D) and were selected during domestication (*9*).

Next, to investigate the regulatory basis underlying spikelet formation and kernel row number divergence, we compared teosinte and maize spikelet meristem cell types across three developmental stages. Although overall clustering revealed conserved spikelet meristem cell identities (Fig. 2F), the spatial and temporal expression patterns of key domestication regulators were extensively rewired during maize domestication. At 2 mm, *TB1* (*Teosinte Branched1*) (*25*) was more broadly expressed in the maize meristem tip epidermis (MTE) (Fig. 2G), consistent with the developmental suppression of lateral organs, such as lower florets and bracts by expanded *TB1* expression in maize ears suppresses lateral organs development (*26, 27*). At 4 mm, multiple domestication regulators involved in inflorescence architecture, including *ZFL2* (*28*), *TSH4*, *UB2*, and *UB3* (*29*), had expanded expression domains in the maize meristem periphery epidermis (MPE) compared with teosinte (Fig. 2G). This enhanced expression likely reinforces meristem boundary formation, contributing to the establishment of the multi-rowed spikelet arrangements characteristic of maize. At 6 mm, *RA1* (*Ramosa1*) (*30, 31*) was expressed more broadly in the maize meristem base (MB) relative to teosinte (Fig. 2G). This expression divergence is consistent with the role of *RA1* in promoting spikelet pair meristem determinacy, ensuring differentiation into paired spikelets that produce kernels rather than continued branching. Conversely, *TU1* (*Tunicate1*) (*32*) maintained broader expression in teosinte meristem epidermis (ME) than in maize (Fig. 2G), consistent with its ancestral role in promoting teosinte glume development. Furthermore, using Hotspot, we found that the gene modules containing these key domestication TFs had pronounced spatiotemporal specificity within critical meristem cell types. For example, the gene modules including *TB1*, *ZFL2, TSH4, UB2, UB3*, and *RA1* displayed highly spatiotemporal and cell type specific expression patterns (Fig. 2H and fig. S12A). More than 90% of genes in these spatiotemporally specific gene modules showed no evidence of selection (fig. S12B), but divergence in the expression range encompassed entire gene modules (Fig. 2I), suggesting that selection acting on a limited number of upstream regulators propagated through downstream gene networks to generate widespread transcriptomic reprogramming. Collectively, we propose that changes in the spatiotemporal expression domains of key domestication TFs drove the rewiring of their regulatory networks, which in turn altered meristem cell proliferation and fate, thereby shaping kernel row number differences between maize and teosinte.

### *ZmSPD1* controls halving kernel rows in teosinte

The teosinte ear inflorescence produces a spikelet pair meristem (SPM) that divides a sessile spikelet (SS) adaxially and a pedicellate spikelet (PS) abaxially at each node (*6*). PS development is halted early, so only the SS flowers, thereby halving the number of kernel rows (*6*). The phenotypic transition from semi-fertile spikelets of teosinte to fully fertile spikelets of maize constitutes a key step during domestication. Our snRNA-seq analyses have revealed that changes in spatiotemporal expression of multiple domestication genes across distinct meristem cell types drove ear morphological divergence. Nevertheless, the core domestication gene responsible for this process remains unclarified. To complement our snRNA-seq data and more closely investigate this trait, we previously employed a classical fine-mapping approach in a maize-teosinte BC_2_F_7_ near-isogenic line (NIL) population (Fig. 3A) (*34*), and mapped a major QTL to a ∼300 kb locus on chromosome1 named *Ter1* (*Teosinte ear rank1*) (*34*).

**Fig. 3.**
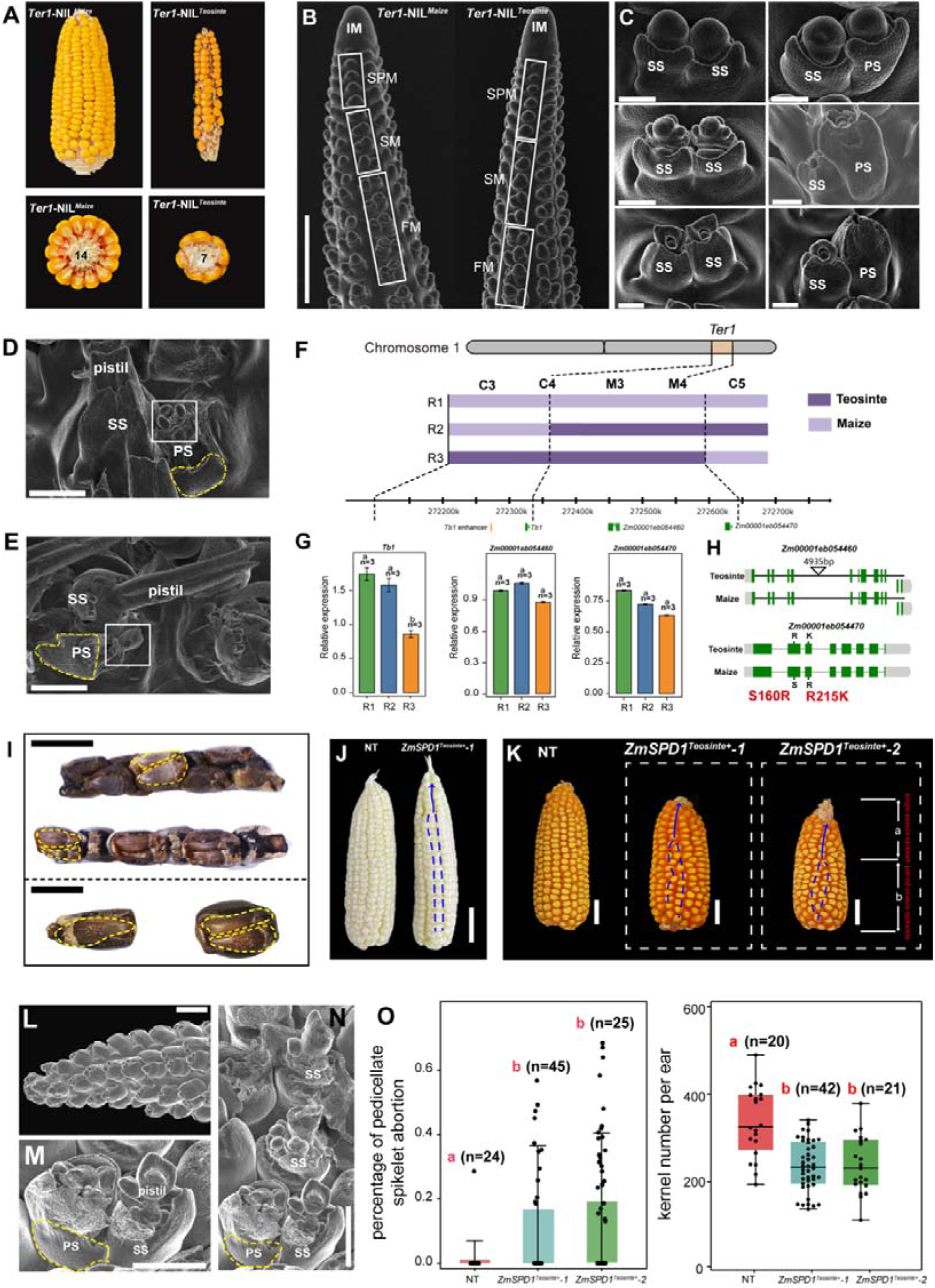
*ZmSPD1* controls paired spikelet abortion and underwent selection during maize domestication. **(A)** *Ter1*-NIL*^Teointe^* shows a kernel row number half that of *Ter1*-NIL*^Maize^*. **(B)** Scanning electron micrographs of ∼5 mm developing ears of *Ter1*-NIL*^Teointe^* and *Ter1*-NIL*^Maize^* sibling showed no difference at IM, SPM and the early stage of SM, scale bar = 1mm. **(C)** Close-up scanning electron micrographs of FM of *Ter1*-NIL*^Teointe^* and its normal sibling (left) with glume removed, scale bar = 100 μm. **(D-E)** Close-up scanning electron micrographs of FM with glume removed in *Ter1*-NIL*^Teosinte^*, the yellow dashed box indicates the stalk-like structure on the base of the meristem, scale bar = 100 μm. **(F)** These NILs was employed to exclude the regulatory effect of *TB1*. Dashed lines indicate the QTL intervals. **(G)** Relative expression levels of genes in the R1, R2, and R3 haplotypes from (F) in the 300 kb region. **(H)** Genomic sequence variation of the candidate genes (30). **(I)** Top, Teosinte with double rows of SS; the yellow dashed box indicates pairs of SS; scale bar = 1cm; Bottom, Detail of single (left) and paired SS of teosinte. Scale bar = 0.5cm. **(J-K)** Spikelet abortion phenotype in the immature (J) and mature ears (K) from NT (non-transgenic line) and *ZmSPD1^Teosinte+^* lines. Scale bar = 2cm. **(L-N)** Close-up scanning electron micrographs of FM with glume removed in *ZmSPD1^Teosinte+^*, the yellow dashed box indicates the stalk-like structure on the base of the meristem. Scale bar = 100 μm. **(O)** Comparison of percentage of pedicellate spikelet abortion and kernel number per ear of NT and *ZmSPD1^Teosinte+^* lines.

To determine if *Ter1* influences female spikelet development, we examined developing ears of *Ter1*-NIL*^Teosinte^*and *Ter1*-NIL*^Maize^* using scanning electron microscopy. In *Ter1*-NIL*^Maize^*, each SPM produces two unequally sized spikelet meristems but eventually forms morphologically similar SS (Fig. 3B). Conversely, in *Ter1*-NIL*^Teosinte^*, each SPM produces one SS, while the other develops into a teosinte-like PS with a stalk-like structure at its base (Fig. 3C). The SS of *Ter1*-NIL*^Teosinte^* develops a female floret with an ovule and silk, and stamens are arrested (Fig. 3, D and E). However, the floret meristem fails to produce a fertile female floret in the PS of *Ter1*-NIL*^Teosinte^*(Fig. 3, D and E). It either forms stamen primordia with halted pistil development or aborts early (Fig. 3, D and E). This leads to half as many fertile florets and fewer kernel rows on the developing ear of *Ter1*-NIL*^Teosinte^*.

The QTL region includes two annotated genes, a RING-type E3 ligase *(Zm00001eb054460),* and a putative membrane transporter *(Zm00001eb054470)*, and excludes the well-known domestication gene *TB1,* located approximately 2.5 kb away. Because *TB1* plays an important role as a transcriptional regulator of several other domestication genes (*27*), we evaluated the expression of *TB1* and both candidate genes across recombinant NILs. Our results indicate that *TB1* expression changes correlated with haplotype differences in maize and teosinte at its known enhancer locus (*35*), but not with haplotype differences at the *Ter1* locus. Neither of the two candidate genes had significant expression differences across NILs (Fig.3, F and G), consistent with our maize and teosinte snRNA-seq analysis (table S8 and S9). Combined, these results suggest that maize and teosinte differences at *Ter1* are not regulatory and are independent of *TB1*.

Subsequent sequence alignment revealed no amino acid changes in *Zm00001eb054460* between maize and teosinte, whereas *Zm00001eb054470* carried two nonsynonymous SNPs resulting in two amino acid changes (Fig. 3H) (30). Resequencing data further showed that both nonsynonymous variants occurred at significantly higher frequencies in maize than in teosinte (*34*), and XP-EHH analysis detected strong evidence of selection spanning the gene (fig. S13) [XP-EHH (chr1:272,634,050..272,638,050) = 21.35; *Zm00001eb054470* (chr1:272632427..272638723); (selection threshold=3.68)]. We also identified several rare teosinte accessions from wild populations that have double rows of SS, and sequencing analysis revealed that all of these plants carried the maize haplotype of the *Zm00001eb054470* gene (Fig. 3I and fig. S14). We therefore propose *Zm00001eb054470* as the key candidate gene controlling PS abortion, and refer to this gene hereafter as *ZmSPD1 (Spikelet Domestication 1)*.

To functionally validate *ZmSPD1* as the causal locus controlling PS abortion, we expressed the teosinte allele of *ZmSPD1* in the maize inbred KN5585 using the native teosinte promoter (*ZmSPD1^Teosinte+^*). Consistent with our interpretation that *ZmSPD1* is the causal gene, transgenic lines (fig. S15) had distinctive single-row phenotype at the tip of both immature and mature ears (Fig.3, J and K). Scanning electron microscopy also revealed that *ZmSPD1^Teosinte+^* plants had aborted PS and a stalk-like structure at the base of the meristem, similar to that observed in *Ter1*-NIL*^Teosinte^* (Fig.3, L to N). This developmental defect led to irregular kernel row organization and a higher percentage of pedicellate spikelet abortion, accompanied by ∼30% reduction in kernel number per ear relative to non-transgenic line (NT) controls (Fig. 3O), demonstrating that *ZmSPD1* is a key domestication gene controlling PS abortion.

### SnRNA-seq analysis of *Ter1*-NIL*^Teo^* and *Ter1*-NIL*^Maize^* ear

To investigate the molecular mechanism by which *ZmSPD1* causes spikelet abortion in teosinte, we performed snRNA-seq on ear tissue from *Ter1*-NIL*^Teosinte^* and *Ter1*-NIL*^Maize^*plants. UMAP clustering of more than 25,000 nuclei across three biological replicates of each genotype (table S10) was conducted, with cell types annotated as described above (Fig. 4A, fig. S16 and table S11). Spatial patterns of marker gene expression revealed that corresponding cell types exhibited similar distributions between the two genotypes (fig. S17), and comparative analysis of cellular composition revealed nearly identical proportions of major cell types (fig. S18A). *ZmSPD1* had comparable expression range and mean expression levels in both genotypes (fig. S18B), consistent with our quantitative RT-PCR results (Fig. 3G). These results further suggest that amino acid changes, rather than expression changes, in *ZmSPD1* are causative of kernel row number domestication.

**Fig. 4.**
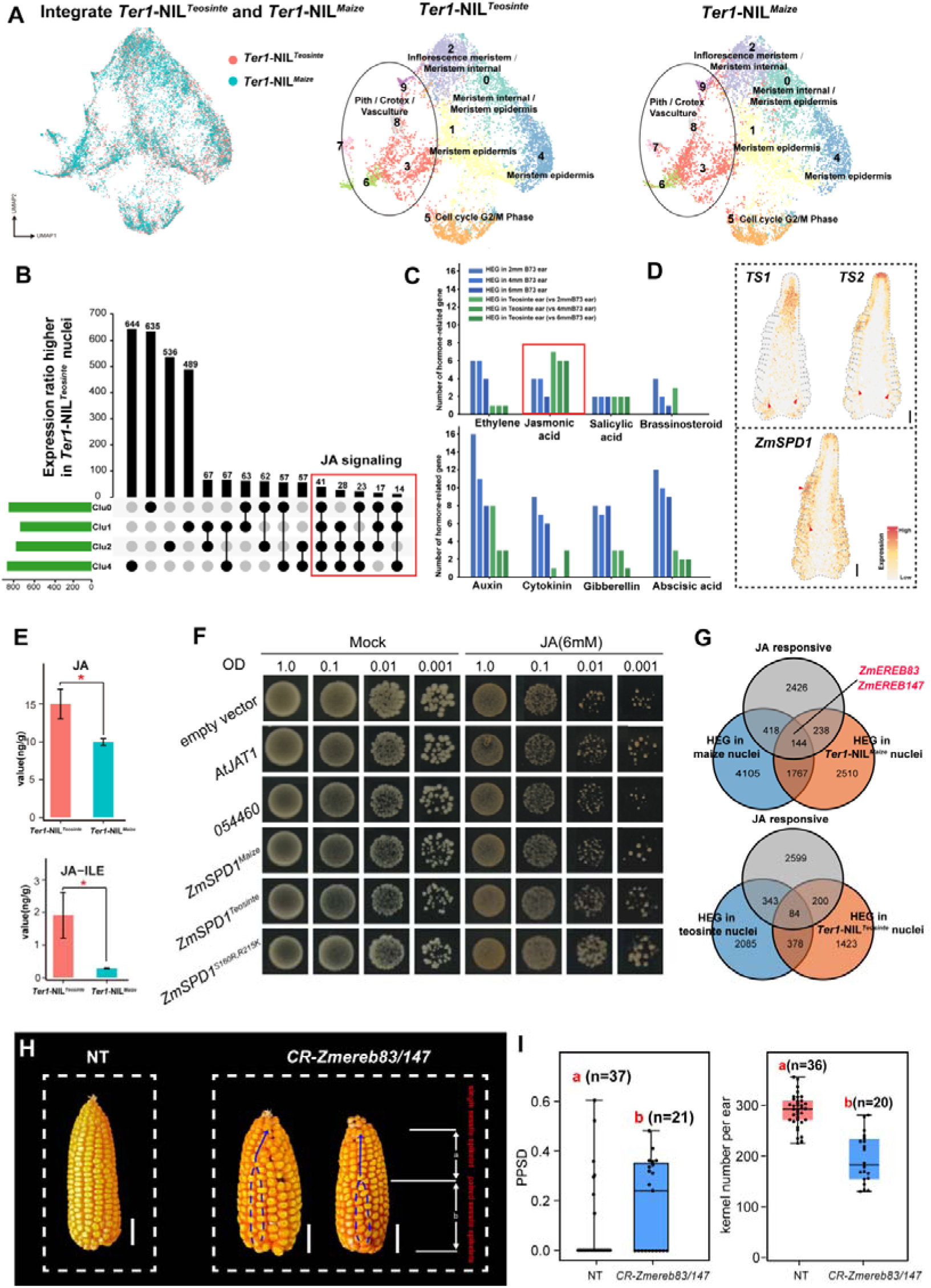
Molecular and biochemical characterization of *ZmSPD1* mediated regulation of ear development. **(A)** UMAP visualization and clustering of snRNA-seq from *Ter1*-NIL*^Teointe^* and *Ter1*-NIL*^Maize^* ear. **(B)** DEGs identified between *Ter1*-NIL*^Teointe^*and *Ter1*-NIL*^Maize^*. **(C)** Number of hormone-related genes showing significant differences in expression range or average expression level between maize and teosinte spikelet meristem cell types. Bars within each group represent number of highly expressed (higher average expression level or expression range) genes (HEG) in maize spikelet meristem cell types (HEG in 2/4/6 mm B73 ear) and HEG in teosinte spikelet meristem cell types (HEG in teosinte ear compared with 2/4/6 mm B73 ear). **(D)** Stereo-seq expression pattern of maize 6mm ear, and the arrow refers to the gene expressed region that matches the *in situ* result. Scale bar = 0.5mm. **(E)** Quantification of JA levels: jasmonic acid (JA) and jasmonic acid-isoleucine (JA-ILE) in ear tissues of *Ter1*-NIL*^Teointe^* and *Ter1*-NIL*^Maize^*, error bars indicate SD (n = 3). Asterisks indicate significant differences (Student’s t-test: *P < 0.05). **(F)** Jasmonic acid transport activity of maize and teosinte *ZmSPD1* alleles in a yeast heterologous system. Yeast cells expressing *AtJAT1*, *ZmSPD1^Teosinte^* and *ZmSPD1^Maize^* were diluted (OD = 1.0, 0.1, 0.01, and 0.001) and grown on SD-URA plates supplemented with 6 mM exogenous JA for 3 days. Yeast cells with an empty vector and expressing *Zm00001eb054460* (*054460*) were used as the control. **(G)** Top: HEG in both *Ter1*-NIL*^Maize^*and maize (B73) ear spikelet meristem cell types, and responsive to JA treatment. Bottom: HEG in both *Ter1*-NIL*^Teointe^* and teosinte ear spikelet meristem cell types, and responsive to JA treatment. **(H)** Spikelet abortion phenotype in the mature and immature ears from NT (non-transgenic line) and *CR-Zmereb83/147* lines. Scale bar = 2cm. **(I)** Comparison of the percentage of pedicellate spikelet abortion and kernel number per ear of NT and *CR-Zmereb83/147* lines.

We next aimed to assess the downstream effects of *Ter1* by identifying DEGs specifically in spikelet meristem cells (clusters 0, 1, 2, and 4) that express *ZmSPD1*. This analysis identified a group of jasmonic acid (JA) signaling genes (five genes) with more than 2-fold expanded expression range in *Ter1*-NIL*^Teointe^* spikelet meristem cells (Fig. 4B and table S12). To determine whether this JA signaling transcriptional signature reflected a broader hormonal divergence, we compared the expression of phytohormone genes between maize and teosinte spikelet meristem cells (Fig. 2F). Among all hormone genes showing differential expression, only JA signaling genes were upregulated in teosinte, with more JA signaling genes showing higher expression (higher average expression level or expression range) in teosinte than in maize (Fig. 4C).

We next confirmed the expression pattern of *ZmSPD1* by in situ hybridization and found expression in the meristem tip and vasculature (Fig. 4D and fig. S19). Key enzymes in JA synthesis, such as *Tassel Seed1 (TS1)* and *Tassel Seed2 (TS2)*, are mainly expressed in the meristem base and vasculature (Fig. 4D and fig. S19) (*36*), suggesting that JA may be synthesized in these domains rather than in the meristem tip. Given the expansion in expression of JA signaling genes in the spikelet meristem region (Fig. 4B), and the prediction that *ZmSPD1* is a membrane transporter, we hypothesized that *ZmSPD1* may transport JA from the meristem base and vascular tissues to the meristem tip, thereby influencing kernel row number through JA-mediated pathways.

To verify this, we first quantified phytohormone levels in both NILs and found significantly higher JA levels in *Ter1*-NIL*^Teosinte^*than in *Ter1*-NIL*^Maize^* ears (Fig. 4E and fig. S20). We next expressed the teosinte and maize haplotypes of *ZmSPD1* in yeast and tested whether *ZmSPD1* could transport JA based on previous findings that exogenous JA inhibited yeast cell growth (*37*). Among these yeast transformants, *ZmSPD1^Teosinte^* protein exhibited enhanced tolerance to exogenous JA, suggesting its stronger JA export capacity than the *ZmSPD1^Maize^* protein (Fig. 4F). Moreover, introducing the two teosinte-specific nonsynonymous substitutions into the *ZmSPD1^Maize^* protein (*ZmSPD1^S160R, R215K^)* also increased JA tolerance in yeast (Fig. 4F). AlphaFold3 (*38*) predictions further revealed striking conformational divergence among the *ZmSPD1^Maize^*, *ZmSPD1^Teosinte^*, and *ZmSPD1^S160R, R215K^* (fig. S21). Those results suggest that the nonsynonymous substitutions in the *ZmSPD1^Teosinte^*protein enhance its JA transport activity, contributing to elevated JA accumulation in *Ter1-NIL^Teosinte^* ears.

Next, to determine how elevated JA affects downstream gene expression, we treated normally developing KN5585 immature ears with exogenous JA and performed bulk RNA-seq. We identified 3226 JA-responsive genes (|log2FoldChange| > 2 and padj < 0.01; table S13). Separately, analysis of snRNA-seq data to identify DEGs between *Ter1*-NIL*^Teointe^* and *Ter1*-NIL*^Maize^*, as well as between maize and teosinte, yielded 1,911 genes more highly expressed in *Ter1*-NIL*^Maize^*/maize, and 462 genes more highly expressed in *Ter1*-NIL*^Teointe^*/teosinte spikelet meristem cells (Fig. 4G).

Among the DEG and JA-responsive genes, *ZmEREB83*, an ethylene-responsive element binding TF, had notable changes in expression. The crosstalk between ethylene and jasmonic acid signaling has been implicated in the regulation of spikelet development in maize (*39*). The expression level of *ZmEREB83* was altered following exogenous JA treatment and was consistently higher in both *Ter1*-NIL*^Maize^* and maize, than in *Ter1*-NIL*^Teointe^* and teosinte spikelet meristem cells (Fig. 4G). A close homolog, *ZmEREB147*, was expressed similarly, although with a smaller magnitude of differential expression (Expression range in *Ter1*-NIL*^Maize^*: *Ter1*-NIL*^Teointe^* = 1.7). We therefore generated *Zmereb83/147* double mutant lines by CRISPR-Cas9 (*CR-Zmereb83/147*, fig. S22), and observed a PS degeneration phenotype and ∼33% reduction in kernel number per ear (Fig.4, H and I), indicating that *ZmEREB83/147* participates in regulating spikelet abortion processes.

Taken together, our findings establish a regulatory pathway underlying *ZmSPD1*. Structural variation in *ZmSPD1* enhances JA transport activity, resulting in elevated JA levels in the teosinte spikelet meristem. Through a mechanism likely mediated by JA–ethylene crosstalk, the increased JA subsequently alters the expression of downstream target genes, including *ZmEREB83* and *ZmEREB147* (downregulation). This ultimately leads to abortion of one spikelet in each pair in teosinte, and an effective doubling of kernel number in maize.

## Discussion

The transition from the small, two-rowed ear of teosinte to the modern maize ear represents a classic example of rapid adaptive morphological evolution. Crosses between maize and teosinte have uncovered a number of genes with variants that contribute to this transition (*40, 41*), but we have lacked a holistic comparison of the genetic differences between maize and teosinte ears. Here, we use spatial and single-nucleus transcriptomics to characterize cell composition and gene expression changes underlying this transition. Perhaps surprisingly, we show that the radical shift in gross ear morphology largely made use of existing cell types, accompanied by extensive rewiring of gene expression and regulatory networks. This observation potentially links cob morphology evolution to the well-characterized *TSH4-UB2-UB3* regulatory network. The *TSH4-UB2-UB3* module regulates maize inflorescence architecture by controlling meristem fate specification (*29*), with *UB3* additionally modulating cytokinin homeostasis and signaling (*19*). Our findings raise the possibility that this regulatory module may have broader roles beyond spikelet patterning, potentially contributing to cob thickening during maize ear evolution through cytokinin-mediated cell proliferation. We also observed downregulation of two negative regulators of organ growth, *bb2* and *CRN6*, in maize pith cells. Although previous studies have implicated their roles in controlling organ size through regulation of cell number (*20*) (*22*), their relationship with cytokinin signaling remains unclear. Our data suggest a potential indirect antagonistic relationship between *bb2/CRN6*-mediated growth restriction and cytokinin-driven proliferative activity, which requires further investigation. Collectively, these observations suggest that maize enlarged its cob through multiple, complementary transcriptional strategies, both promoting cell proliferation and releasing latent growth potential.

Despite these cob-specific changes, spikelet meristem cell types were largely conserved between maize and teosinte. Nonetheless, the expression patterns of certain key genes have shifted. Early in development (2 mm ear primordia), numerous genes associated with cell division are upregulated in maize IM cells, accompanied by enhanced proliferative capacity that could generate a larger pool of developmentally competent cells. At later stages (4 mm and 6mm ears), changes in regulatory networks centered on key domestication genes such as *ZFL2*, *TSH4, UB2, UB3, and RA1* are observed, which are deployed across a broader spatial and temporal range in maize, recruiting meristem epidermis periphery and meristem base cells into specific developmental programs. Our observations suggest that selection acted primarily by expanding the expression domains of a limited number of key regulatory genes within a small subset of meristematic cell types, reshaping cellular composition (fig. S23) and thereby coordinately influencing both cob and spikelet development. Crucially, this mechanism depends on changes in expression domain rather than substantial changes in per-nucleus average expression levels. snRNA-seq, unlike standard bulk RNA-seq, is able to distinguish altered spatial deployment of gene expression across cell types and developmental stages from uniform changes in transcriptional output within individual cells.

Additional insight into maize evolution came from fine-mapping of a novel domestication gene *ZmSPD1*. We identified two amino acid substitutions as the key suppressor of spikelet abortion and a doubling of kernel row number during domestication. Treatment with JA (Fig. 4G) and knockout experiments (Fig. 4H and I) confirmed that these substitutions are responsible for changes in downstream regulation of JA-responsive genes, such as *ZmEREB83* and *ZmEREB147.* Given the complexity of the JA signaling network, additional *ZmSPD1*-mediated regulation of downstream components may await identification, thereby deepening understanding of how biotic/abiotic stress-related phytohormones are crucial for inflorescence development, domestication, and plant adaptation.

Together, our results help resolve a deeper model of how selection acted during maize domestication. First, our results highlight how domestication largely made use of existing variation. Our spatial and single-nucleus transcriptomic data reveal that the maize ear, especially the spikelet, has conserved cell types already present in teosinte, with differences confined to the amount and timing of cell proliferation. Like other domestication genes (*42*), the maize haplotype of *ZmSPD1* already existed as standing variation in teosinte. These results reinforce the utility and importance of wild populations as valuable reservoirs of genetic diversity (*2, 43–45*). Our data also help resolve apparent conflicts between early results from QTL mapping that showed a small number of loci responsible for domestication (*46, 47*) and population genetics work suggesting that domestication was highly polygenic (*7, 48*). Our transcriptomic data found a key role in early development for a relatively small number of large-effect regulatory loci with clear evidence of selection, each of them driving widespread transcriptional programs. In addition, our genetic analysis of spikelet abortion is consistent with other domestication traits (*5, 49*) in showing contributions from a number of loci beyond the largest effect QTL (*40, 41, 50*) and highlighting the effect of genetic background (*49*) (table S14). Combined, these data suggest that maize domestication targeted polygenic traits for which teosinte harbored substantial genetic variation (*51*). Consistent with simulations of large optimum shifts for polygenic traits (*52*), selection focused on large-effect loci, here genes that allowed large scale rewiring of downstream regulatory networks.

## Materials and Methods

### Plant Growth Conditions

Maize B73 inbred plants and teosinte (*Zea mays ssp. parviglumis*) were grown either in the field (April–July) at Huazhong Agricultural University (Wuhan and Sanya) or in a greenhouse under controlled conditions (12 h light/12 h dark, 26–28°C day, 22–24°C night). Developing ears (∼2 mm, 4 mm, and 6 mm) were collected at the 6, 8, and 11 leaf stages. The maize mutant transgenic lines were generated in the KN5585 background using Agrobacterium-mediated transformation. Mutations and transgene insertions were confirmed by PCR with primers listed in table S15.

### Tissue fixation

Developing teosinte ears (1-3 mm) were collected and immediately fixed in a 4% formalin-acetic acid-alcohol (PFA) solution for 2 minutes. The tissues were then treated with 2% sucrose solution (1×PBS buffer, pH=7.4, 2% sucrose, v/m) and vacuumed on ice for 15 minutes each, then the solution was changed and vacuum applied for an additional 15 minutes. The ears were then embedded in pre-chilled OCT (Sakura), snap-frozen in liquid nitrogen pre-chilled isopentane for 10 seconds, and stored at −80°C for later use.

### Scanning Electron Microscopy (SEM)

Immature ears from B73 (∼2-6 mm) Teosinte ear (∼1-3 mm), *Ter1*-NIL*^Teosinte^* and *Ter1*-NIL*^Maize^* plants were collected. Husk leaf primordia were carefully removed, and the ears were observed using a JEOL JSM-7900F SEM. Samples were processed as previously described (*16*): briefly, ears were fixed in 2.5% glutaraldehyde, dehydrated in a graded ethanol series, critical-point dried, and sputter-coated with gold. SEM observations were performed at 5–10 kV accelerating voltage.

### mRNA *in situ* Hybridization

Immature teosinte ears (1–3 mm) were fixed in 4% paraformaldehyde (PFA) in 1× PBS (pH 6.5–7.0) at 4°C for 16–24 h. Samples were dehydrated in ethanol series, cleared in histoclear, and embedded in Paraplast Plus (Sigma, P3683). Sections were cut at 10 μm thickness. Sense and antisense RNA probes were generated by PCR amplification of target fragments (primers listed in table S15), incorporating a T7 promoter sequence (CATTAATACGACTCACTATAGGG) in the 5′ or 3′ end as appropriate. Probes were transcribed in vitro using T7 RNA polymerase (*53*) and labeled with digoxigenin-UTP. Hybridization, immunological detection, and signal capture were performed following established protocols.

### Single nucleus isolation

The frozen materials were placed in 3 ml of Nuclear Isolation Buffer A (NIBA:0.8 M sucrose, 10 mM MgCl2, 25 mM Tris-HCl (pH 8.0), 0.1 mM DTT, 0.4 U/μl RNase inhibitor and 0.1 mM PMSF) at 4 °C. The nuclei were harvested by chopping in NIBA at 4 °C. The mixture was subsequently filtered through 40 μm, 30 μm strainer and rinsed with 7 ml of NIBA, and then incubated for 10 minutes at 4 °C. Afterward, nuclei were centrifuged at 50 g for 5 min at 4 °C, the pellet was then removed and the supernatant was centrifuged at 2000 g for 10 min at 4 °C. The pellet was resuspended in 1ml Buffer B (0.4 M sucrose,10 mM MgCl2, 25 mM Tris-HCl (pH8.0), 0.2 U/μl RNase inhibitor, 0.1 mM PMSF, 1% Triton X-100). Percoll gradients were prepared on ice, i.e., 1 volume 75% percoll was added to tube, followed by 1ml 25% percoll. The nuclei suspension was then centrifuged at 3000g for 15min at 4 °C. The middle layer containing most of nuclei was transferred to a new tube and then washed with 5 volumes of Buffer B, and recovered by centrifuging at 1800g for 10 min at 4 °C. The nuclei pellet was resuspended in cell suspension buffer (CRB). The nuclei activity and concentration were measured by DAPI staining and hemocytometer, respectively.

### Single-nucleus transcriptome sequencing (snRNA-seq) and raw data processing

Single-nucleus suspensions were used for library preparation according to the DNBelab C Series Single-Cell Library Prep Set (MGI, 1000021082) as previously described (*54*). The concentration of DNA library was measured by Qubit (Invitrogen). Libraries were sequenced by DNBSEQ-T7RS. The raw sequencing reads were filtered and demultiplexed by PISA (Version 1.1.0) (https://github.com/shiquan/PISA), and aligned to the B73_v5 (*55*) reference genome using STAR (version 2.7.4a) (*56*) with default parameters. Then we obtained the raw matrix of Cell-gene UMI counts by using PISA.

### snRNA-seq Analysis

Sequencing reads from multiple biological replicates of B73, teosinte, *Ter1*-NIL*^Teosinte^* and *Ter1*-NIL*^Maize^* ears were aligned to the B73 v5 reference genome. The replications of maize and teosinte shared similar gene and UMI numbers in each cell, supporting the reproducibility of the snRNA-seq data (fig. S24 and fig. S25). Downstream analyses were performed using Seurat (v4.1.1) (*57*), which was implemented in R (4.3.1). Data were log-normalized (NormalizeData, scaling factor = 10,000), variable genes were identified with FindVariableFeatures (vst method, 2,000 features), and PCA was performed (RunPCA). Batch effects were corrected using Harmony (RunHarmony, fig. S26). A shared nearest neighbor (SNN) graph was constructed, and clustering was performed with the Louvain algorithm (FindNeighbors and FindClusters). Dimensionality reduction and visualization were conducted using UMAP (RunUMAP).

### Cluster Marker Identification

Cluster-enriched genes were identified with FindMarkers (Seurat) using a two-sided Wilcoxon rank-sum test. Thresholds: log2 fold change >0.25, adjusted p-value <0.01, and >30% of cells in the cluster expressing the marker gene.

### MetaNeighbor Analysis

MetaNeighbor (R v1.20.0) (*58*) was used to assess cell type similarity between clusters. AUROC scores indicated the correlation degree between cell groups.

### Stereo-seq Library Preparation and Sequencing

Developing ear tissues were longitudinally sectioned at 10 μm using a Leica CM1950 microtome. Sections were adhered to the Stereo-seq chip surface (*59*), incubated at 37°C for 2 min, fixed in methanol, and incubated at −20°C for 40 min. Nuclei were stained with a nucleic acid dye (Thermo Fisher, Q10212) for 5 min, and tissue integrity was checked under a microscope. Bright-field and fluorescent images were captured with a Motic PA53 Scanner. Sections were de-crosslinked in TE buffer (10 μM Tris, 1 μM EDTA, pH 8.0) at 55°C for 1 h, permeabilized at 37°C for 12 min, and subjected to reverse transcription overnight at 42°C. Tissue was digested at 37°C for 30 min and treated with Exonuclease I (NEB, M0293L) for 1 h at 37°C. cDNA was purified with Ampure XP beads (0.6× and 0.15×, Vazyme N411-03), used for DNA nanoball (DNB) generation, and sequenced on an MGI DNBSEQ-Tx platform (paired-end 50 bp or 100 bp).

### Raw Stereo-seq data processing and quality control

Stereo-seq raw reads were generated from a MGI DNBSEQ-T5 sequencer. Read 1contained CID and UMI sequences (CID: 1-25bp, UMI: 26-35bp), while read 2 contained the cDNA sequence. Retained reads were then aligned to the reference genome B73 via STAR. and mapped reads with MAPQ 10 were counted and annotated to their corresponding genes using an in-house script (available at https://github.com/BGIResearch/handleBam) and generated a gene-location expression matrix containing location information.

### Binning data of spatial Stereo-seq

After obtaining raw spatial data, transcripts captured by 50 × 50 DNA nanoballs were merged into one bin50 (∼25μm × 25μm). We treated the bin50 as the fundamental analysis unit, and bin IDs were composed of X and Y coordinates on the capture chip. The threshold for filtering low-quality bins was set to < 150 or >5000 gene counts. After filtering, the remaining bin50s were included in the downstream analysis; details are in Table S4.

### Recovering missing values and unsupervised clustering of Stereo-seq data

The raw gene-bin50s matrices were loaded into the Seurat package (4.1.1), which was implemented in R (4.3.1). We then performed normalization in the original dataset (LogNormalize, scaling factor 10,000), using ‘FindVariableFeatures’ function (vst method, 2000 features) to identify highly variable genes, scaled the data with the ‘ScaleData’ function, performed PCA analysis with the RunPCA’ function (100 principal components), and determined statistical significance of PCA scores with the ‘JackStraw function. Clusters were identified using the Seurat function ‘FindClusters’ with ‘resolution = 1’. The table Structures were separately visualized and explored by UMAP (run the ‘RunUMAP’ function with ‘dims = 15, metric = correlation, min.dist = 0.01’). However, many genes were only expressed in only a subset of bin50s. We therefore used the Seurat Wrapper function RunALRA (*60*) (*61*) to impute missing expression values, increasing the non-zero percentage of expressed genes from 46.1% to 81.9% in three teosinte sections (fig. S27). After obtaining the imputed matrices, we performed the downstream analyses as previously described. Briefly, We then detected variable genes with the ‘FindVariableGenes’ function (vst method, 2000 features), scaled data with ‘ScaleData’ function, performed PCA analysis with the ‘RunPCA’ function, clustered bin50s with the Louvain method (‘FindNeighbors’ and ‘FindClusters’), and visualized data with non-linear dimensional reduction algorithms (‘RunUMAP’). The physical distribution of cell types before and after imputation was consistent (fig. S28). More importantly, the clustering of imputed datasets showed clearer anatomical characteristics in some regions, especially in the meristems and vasculature (Fig. 1, D).

### Integration of snRNA-seq and Stereo-seq Data

The proportion of cells from each snRNA-seq cluster of maize ear in spatial spots was estimated using STRIDE (*14*). The proportion of cells from different Sn-clusters (snRNA-seq clusters) in the bin50s are listed in table S2.

### Integration of maize and teosinte snRNA-seq Data

Integration of maize (B73) and teosinte snRNA-seq datasets was performed using the Seurat standard pipeline. Briefly, highly variable genes were first identified in each dataset using FindVariableFeatures (vst method, 2000 features). Integration anchors between the two datasets were then determined using FindIntegrationAnchors (dims = 1:30). The datasets were integrated with IntegrateData (dims = 1:30) to generate a batch-corrected expression matrix. The integrated data were scaled, subjected to PCA, and clustered using a shared nearest neighbor graph (FindNeighbors and FindClusters, resolution). UMAP (RunUMAP) was used for visualization, and cluster marker genes were identified to assign biological cell-type identities.

### Gene module identification

Gene module analysis was performed using Hotspot (*24*). Briefly, normalized single-cell expression matrices were used as input, and highly variable genes were selected based on expression variability across cells. The top 2,000 variable genes were retained for downstream analysis. A cell neighborhood graph was constructed using the low-dimensional embedding generated from the integrated single-cell dataset, and pairwise local correlations between genes were calculated using the Hotspot framework. For each identified module, module scores were calculated across all cells to evaluate activity patterns among cell populations and developmental stages. Modules showing cell type-specific or stage-specific enrichment were further examined. All module scores across cell populations and developmental stages, as well as gene-to-module assignments, are reported in table S16. Default parameters were used unless otherwise specified.

### Co-expression Network Construction and GO/KEGG Enrichment

Gene co-expression networks were constructed using WGCNA (v1.71) (*62*). Average gene expression per cluster was used as input. Modules and gene connectivity were obtained, and gene set enrichment analysis was performed using gProfiler (https://biit.cs.ut.ee/gprofiler/gost). Networks were visualized with Cytoscape (v3.7.1) (*63*).

### Differential gene expression analysis across cell clusters

Differentially expressed genes across cell clusters were identified using a custom, cluster-level summary approach. For each gene, we calculated its average expression level in a single nucleus within each cluster, as well as the proportion of nuclei expressing the gene in that cluster (defined as the expression range, %Expressed). Genes were considered to be expressed in a given cluster if their expression was detected in at least one nucleus. Within each cell cluster, genes showing at least a two-fold difference in average expression level or expression range (%Expressed) between samples were defined as differentially expressed genes. The expression range (%Expressed) was used as an additional metric to characterize the distribution and prevalence of gene expression within each cluster. Key domestication genes identified as differentially expressed genes between maize and teosinte were further validated by in situ hybridization (fig. S29).

### Selective sweeps analysis

The genetic distances between SNPs and SVs were interpolated based on the physical distances from an ultra-high-density genetic map of the maize-wild maize population (*64*). The physical distances were then converted to the B73v5 reference using CrossMap (version v0.7.0) (*65*). Selective sweeps were identified using *Zea mays* subsp. *parviglumis* as the reference population and *Zea mays* subsp. *mays* as the test population. Following previous studies (*9*), 110 accessions of *Zea mays* subsp. *mays* and 70 accessions of *Zea mays* subsp. *parviglumis* were selected, with their high-quality SNPs and pSVs (polymorphic SVs) used for analysis. SNPs were downloaded from the ZEAMAP database (*66*) and subjected to quality control (MAF ≥ 0.05, missing rate < 0.2, and biallelic sites), followed by genotype imputation using Beagle (version r1399, with parameters ‘window=50000 overlap=5000’) (*67*). Beagle is a widely used tool for phasing, imputation, and haplotype inference in genomic data. It employs hidden Markov models (HMMs) and haplotype clustering to estimate missing genotypes and phase variants, enhancing genotype accuracy and enabling robust downstream analyses in large-scale genomic studies. Only SVs without missing data were retained for subsequent analyses. The XP-EHH (Cross-Population Extended Haplotype Homozygosity) values for each locus were calculated using selscan (version v2.0.3, with parameters ‘selscan-2.0.3 --xpehh --max-gap 400000 --unphased’) (*68*). XP-EHH is a statistical method for detecting selective sweeps by comparing haplotype extension differences between two populations, thereby identifying population-specific selection signals. The resulting XP-EHH values were smoothed using GenWin (version 1.0, with parameters ‘smoothness = 100, method = 4’) (*69*) to compute the *W* statistic and spline windows. Genomic regions with W statistic values in the top 5% of the genome-wide distribution were considered putative selective sweep regions. Genes overlapping with these regions were identified as candidate genes under selection.

### Protein structure prediction

The structures of *Zm00001eb054470* were predicted by AlphaFold3 (*38*) under the assumption that each protein forms a dimer. The predicted conformations of the three proteins differ significantly, suggesting that the mutation may have altered their conformations, thereby affecting their functions.

### Bulk RNA-seq analysis

All libraries were sequenced as paired-end 150-bp reads on an Illumina HiSeq 3000 platform. Low-quality reads and adapter sequences were removed using Trimmomatic (*70*). Clean reads were aligned to the B73 RefGen_v5 reference genome using RSEM (*71*) with default parameters, and gene expression levels were quantified simultaneously. Differentially expressed genes were identified using DESeq2 (*72*) with thresholds of |log2 fold change| > 1.5 and adjusted P value < 0.01.

### JA Treatment

Immature ears of *Ter1*-NIL*^Teosinte^* and *Ter1*-NIL*^Maize^* with consistent growth and a length of 1–1.5 cm were selected, and the innermost glumes were retained. Two treatment groups were set up: a control group treated with 0.005% ethanol and a JA treatment group treated with 1 mM jasmonic acid (JA). The respective solutions were applied directly to the inflorescence meristem (IM) region of each ear. Samples were incubated in a 25°C incubator in the dark for 24 h, after which the outer spikelet tissues of the immature ears were dissected and collected for total RNA extraction (*36*).

### Functional Analysis of JA Transport by *AtJAT1* and *ZmSPD1* Variants in Yeast

The CDS sequences of *AtJAT1*, maize haplotype *ZmSPD1^Maize^*, teosinte haplotype *ZmSPD1^Teosinte,^* and the point-mutated variant *ZmSPD1^S160R,R215K^* were individually cloned into the pYES2 yeast expression vector. Empty vector and all recombinant constructs were transformed into Saccharomyces cerevisiae strain Y2H Gold, and positive single colonies were selected on SD-URA medium. Yeast suspensions were adjusted to equivalent optical densities and spotted onto SD-Ura plates with or without 6 mM exogenous jasmonic acid (JA). Plates were incubated at 30°C for 2–3 days, and the JA transport activity of each protein was assessed based on differential yeast growth under JA treatment (*37*).

### Metabolite extraction for maize ear

Lyophilized samples (>1–2 g tissue) were prepared by freezing in liquid nitrogen or at –80°C followed by vacuum freeze-drying. Dried tissues were ground to a fine powder, and 0.05 g aliquots were extracted with 500 μL of solvent containing Acyclovir and Roxithromycin as internal standards. Extracts were vortexed, ultrasonicated for 30 min, centrifuged at 12,000 rpm for 10 min at 4°C, and filtered through a 0.22 μm membrane into HPLC vials. Samples were stored at –20°C short-term or –80°C long-term prior to analysis.

### Metabolite Analysis by LC-ESI-MS/MS

Metabolites were separated on a Waters HSS T3 column (100 × 2.1 mm, 1.8 μm) using a flow rate of 0.35 mL/min and a column temperature of 40°C. The mobile phases were 0.04% acetic acid in HPLC-grade water (solvent A) and 0.04% acetic acid in acetonitrile (solvent B). A 2 μL aliquot of each sample was injected, and the gradient elution program was as follows: 0–1 min, 98:2 A/B; 1–11 min, linear to 50:50 A/B; 11–14 min, 50:50 A/B; 14.1–17 min, 2:98 A/B. Samples were kept at 15°C in the autosampler and analyzed in a randomized order, with quality control (QC) samples evenly inserted to monitor system stability and data reliability. Mass spectrometry was performed on a SCIEX QTRAP 5500 system equipped with an electrospray ionization (*56*) source in positive and negative ion modes. Source parameters were: spray gas 60 psi, auxiliary gas 60 psi, ion spray voltage +5500 V/-4500 V, and source temperature 550°C. Data were acquired in multiple reaction monitoring (MRM) mode. Raw data were processed using OS software, including baseline filtering, peak detection, peak alignment, retention time correction, and peak matching, generating a data matrix containing retention time, m/z, and peak intensity for each metabolite (*73*).

## Statistical analysis

An unpaired two-tailed Student’s t test was used to compare the differences in tested traits between two samples. A oneway analysis of variance (ANOVA) followed by Tukey’s multiple comparison test was used to compare the differences in gene expression levels among three or more samples. Both the two-tailed Student’s t test and ANOVA were carried out in Microsoft Excel.

## Data availability

scRNA-seq and Stereo-seq data from this study can be found in CNGBdb (https://db.cngb.org/), and under project accession code CNP0009173. Additional data, including processed H5ad data, the original gene expression matrix, expression patterns of marker genes across all sections can be accessed from STOmicsDB database (*74*), https://db.cngb.org/stomics/mdesta/.

## Acknowledgements

The work was supported by the National Natural Science Foundation of China (W2511027 to N.Y., 32341029 to L.L, 32401868 to Y.W., 32101778 to M.C., and 32321005 to J.Y.), the Scientific Research Innovation Capability Support Project for Young Faculty (SRICSPYF-ZY2025135 to N.Y.) and the Biological Breeding-National Science and Technology Major Project (2023ZD04073 to N.Y.), the China Postdoctoral Science Foundation (2024T170311) to Y. W. Computation resources were provided by the high-throughput computing platform of the National Key Laboratory of Crop Genetic Improvement at Huazhong Agricultural University and supported by H.L.

## Author contributions

N.Y. and L.L. conceived and supervised this study. Y.W., R.M., Y.L., P.X., Y.W., Y.L., Q.D. prepared the samples for sequencing. Y.W., R.M., X.G., M.C., P.X., Y.W., S.W., E.L., Z.L., L.M. performed the bioinformatics analysis. R.M., Y.L, N.L., Q.Z. finished the transgenic and *in situ* experiment. Y.W., R.M., Y.L., X.G., T.W., H.L., D.M., F.Q., Y.X., X.Y., Z.Z., L.L. and N.Y. discussed the data and prepared the manuscript, J.R.-I, D.J., and J.Y. revised the manuscript. All authors read and approved the manuscript.

