## Supplementary for "The cellular and genetic basis of inflorescence divergence between maize and teosinte"

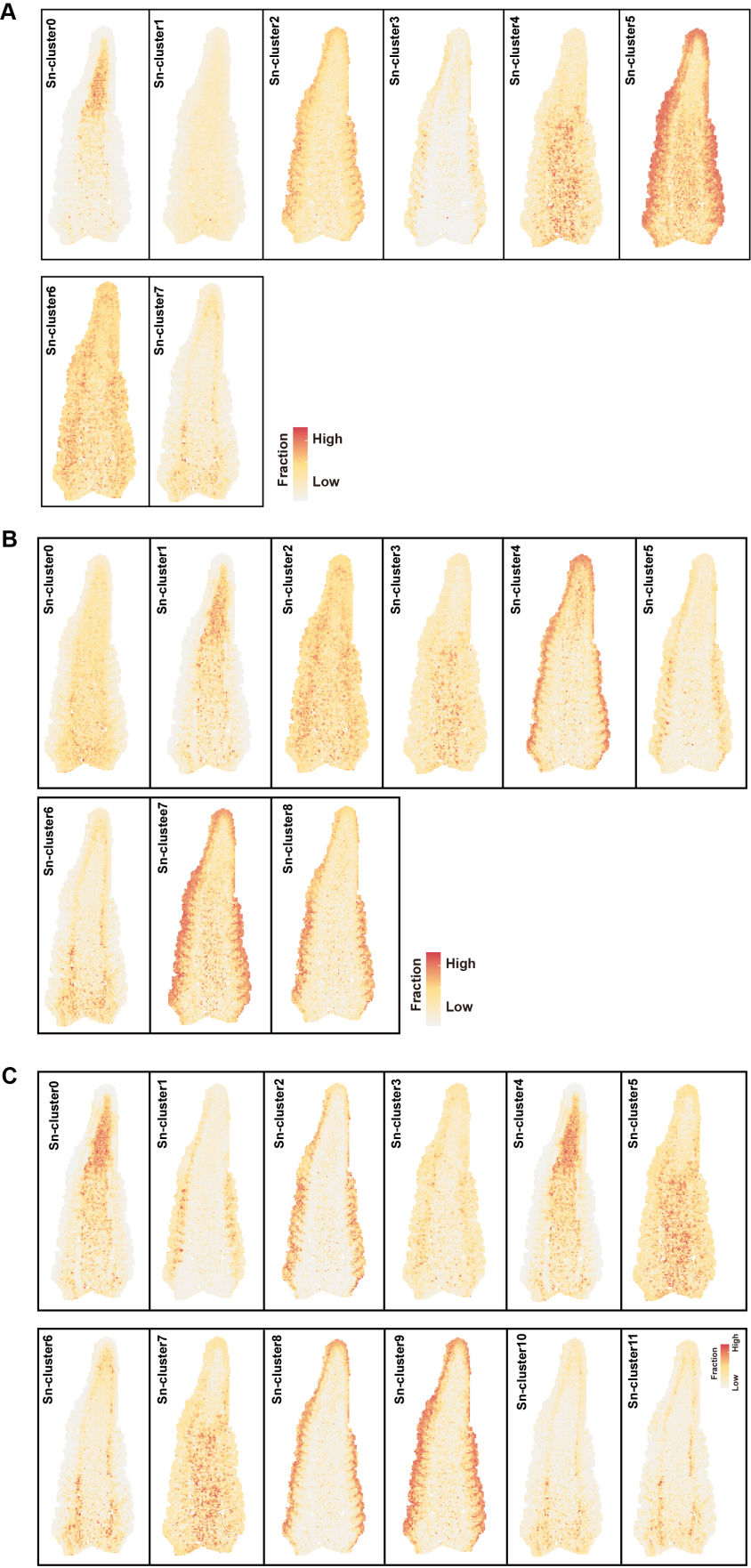

**Fig. S1 STRIDE was applied to project Sn-Cluster onto the maize spatial transcriptome map (A-C).** STRIDE was applied to infer Sn-Cluster identity, with color scale indicating the proportion of cells from different Sn-clusters in the spots. Sn-cluster from 2mm B73 ear (A), 4mm B73 ear (B) and 6mm B73 ear (C).

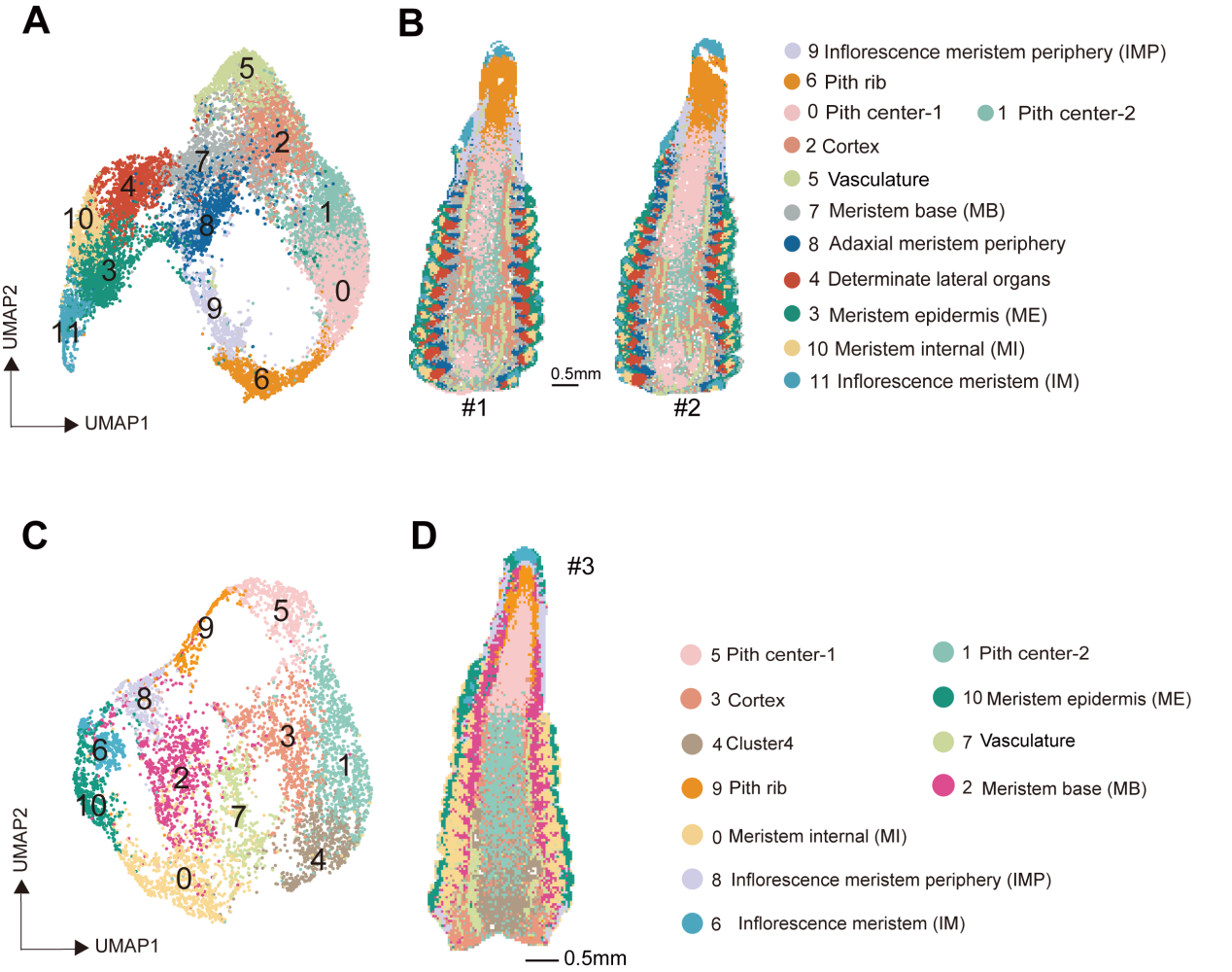

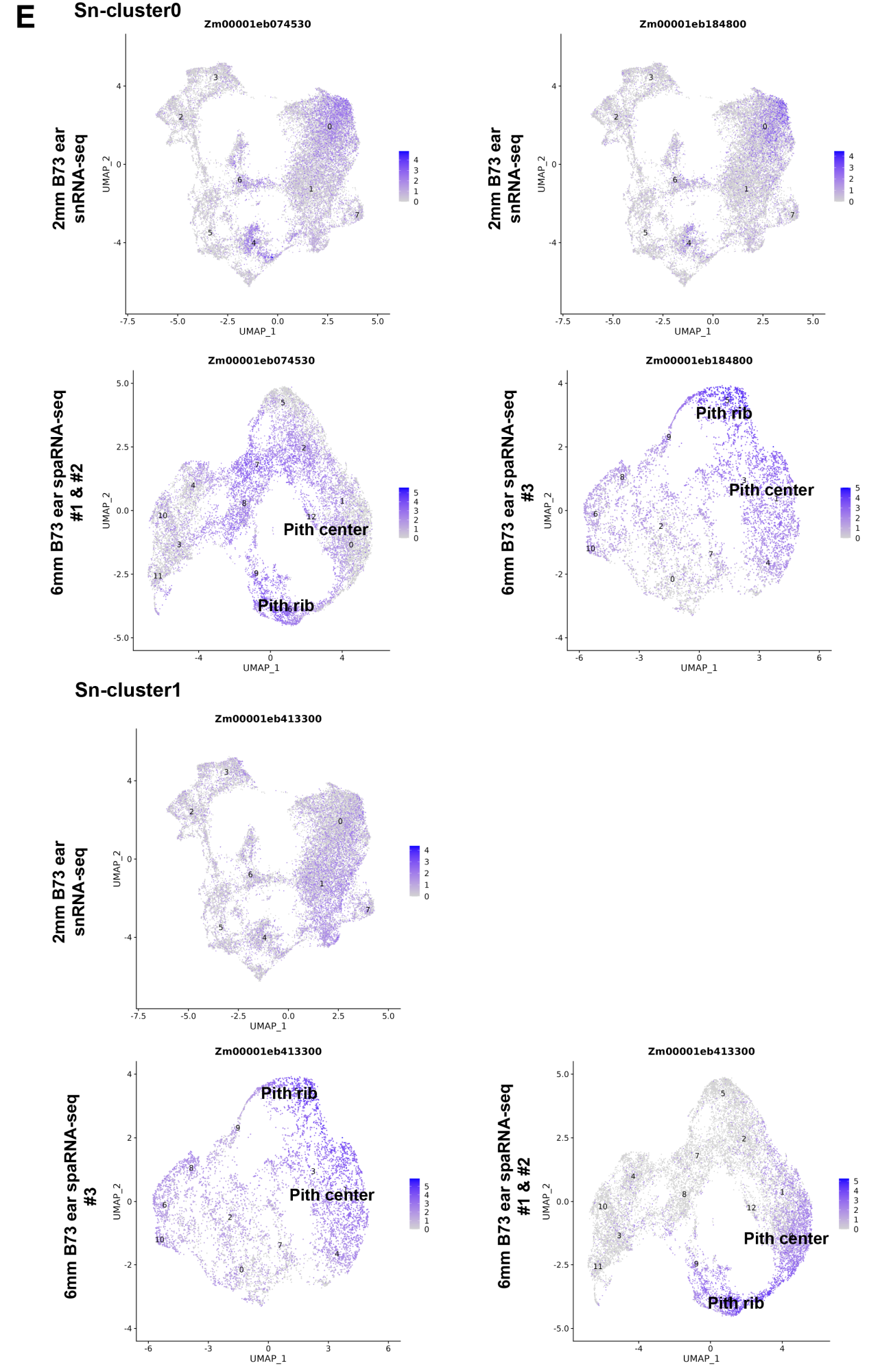

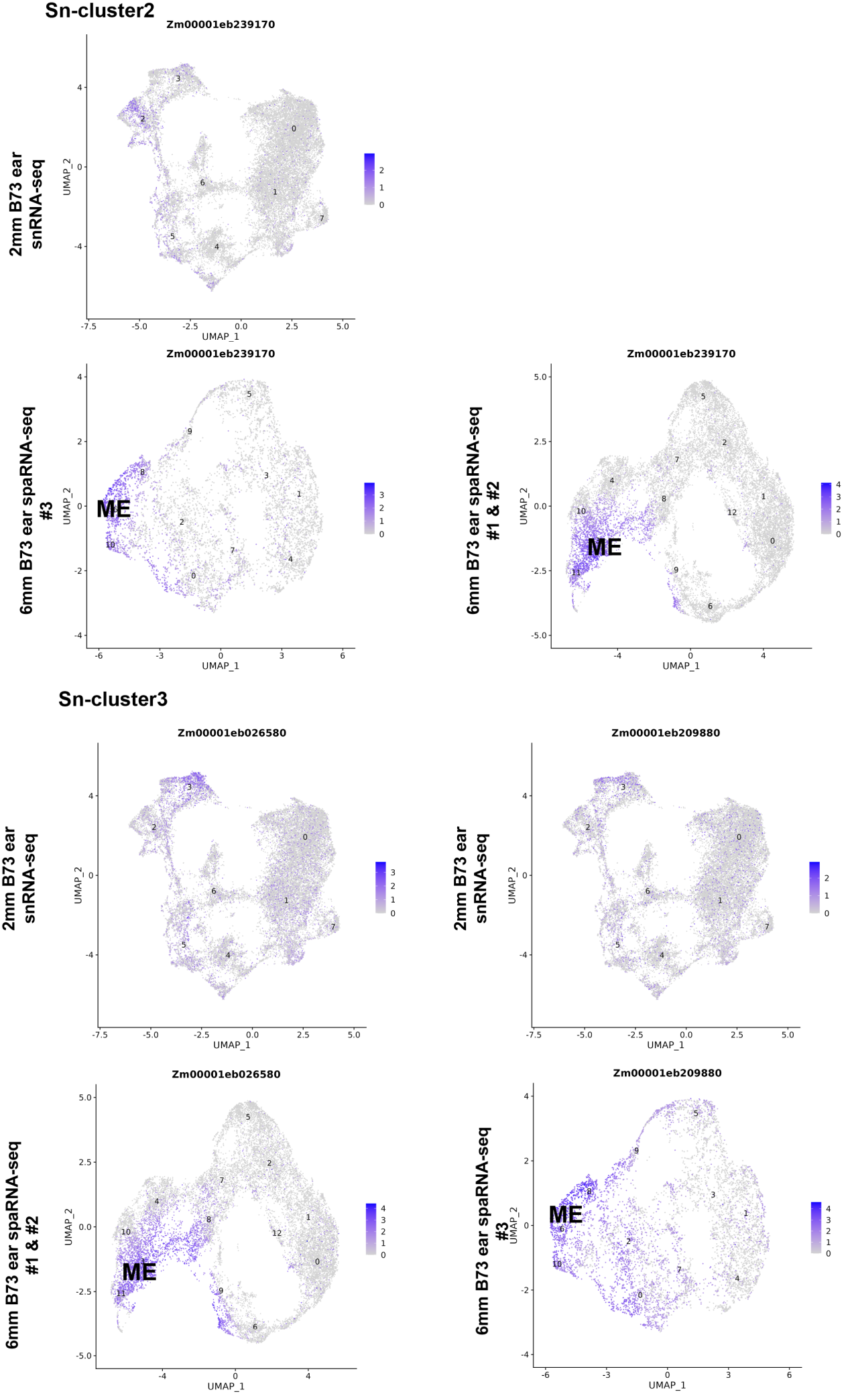

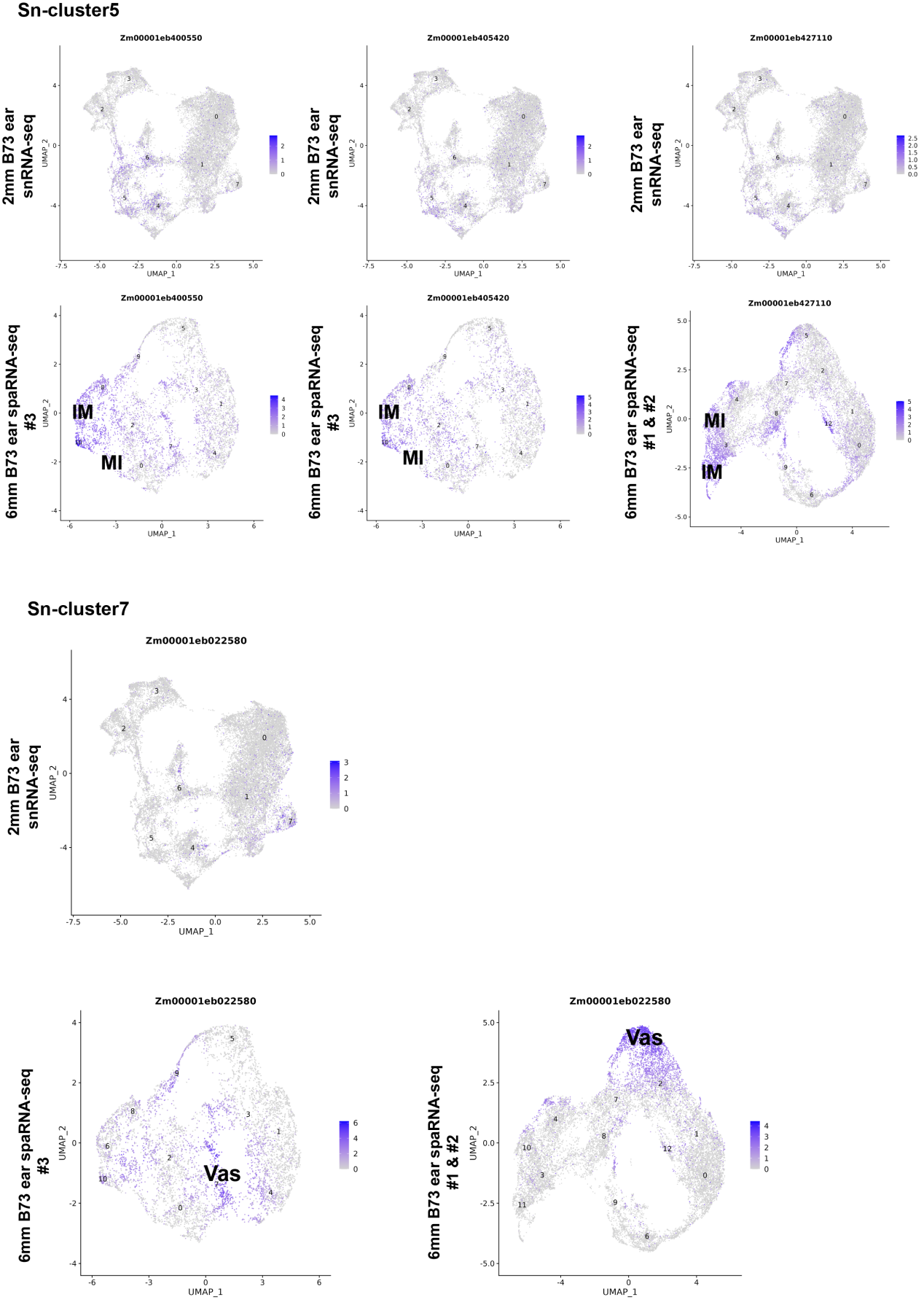

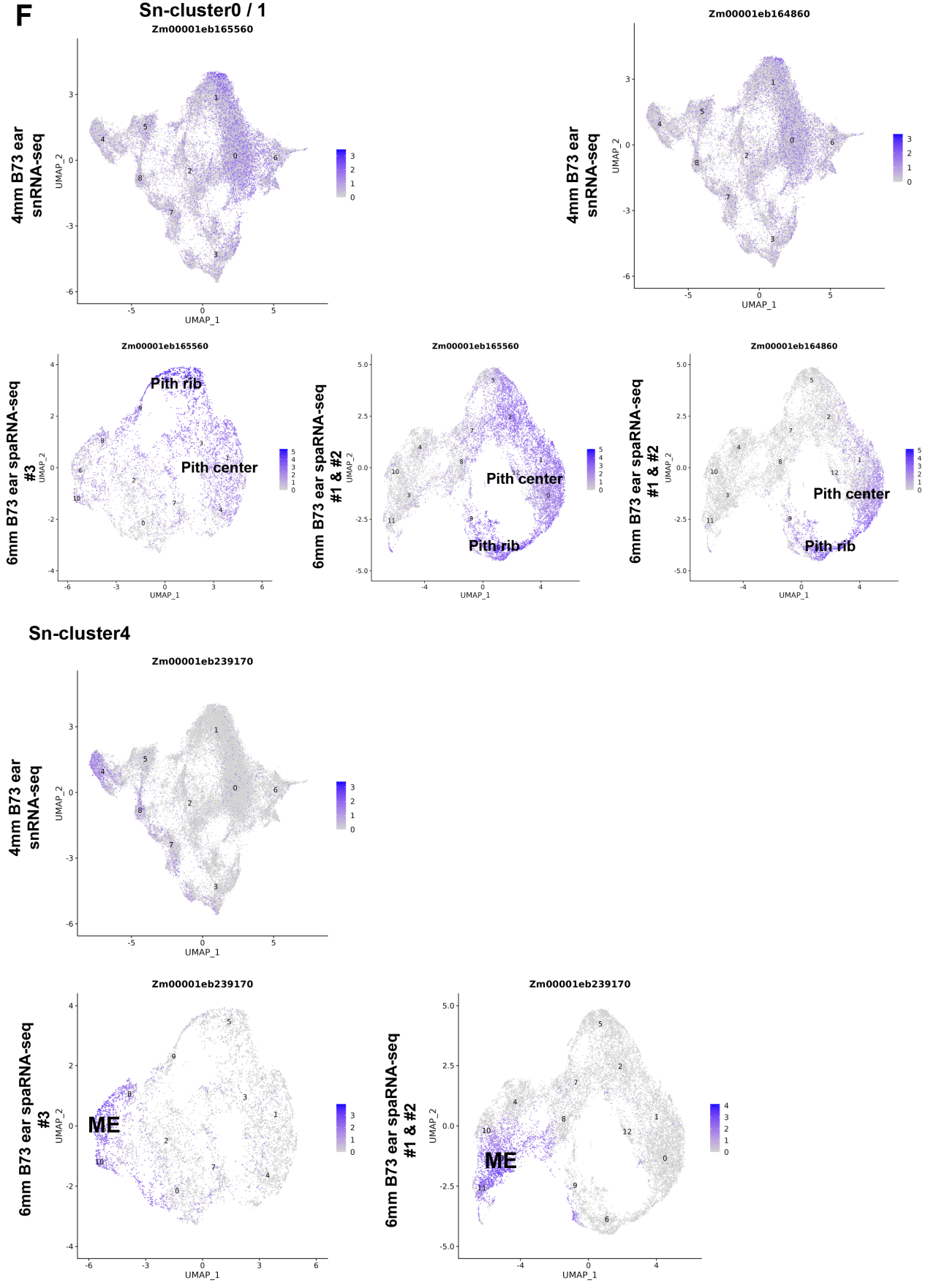

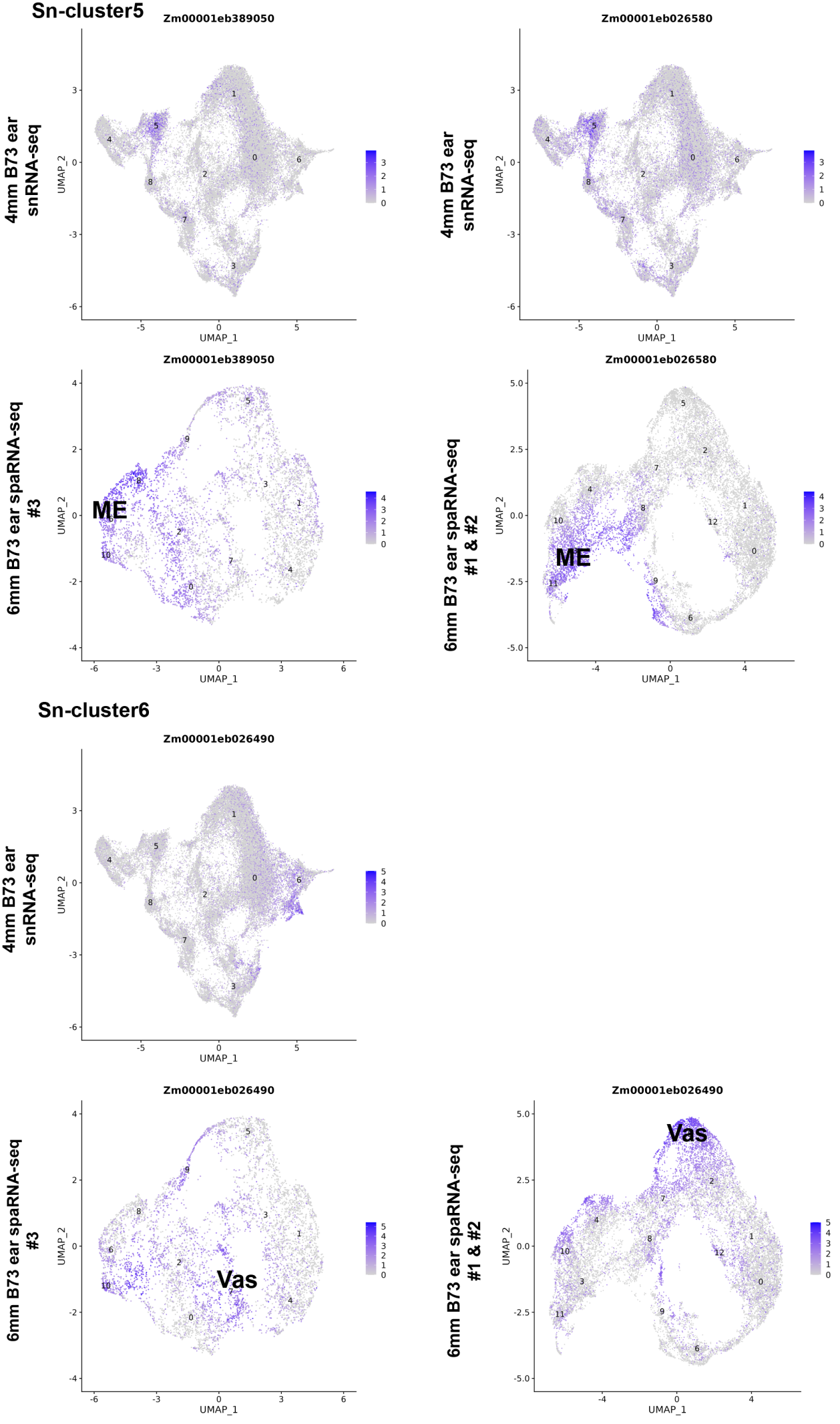

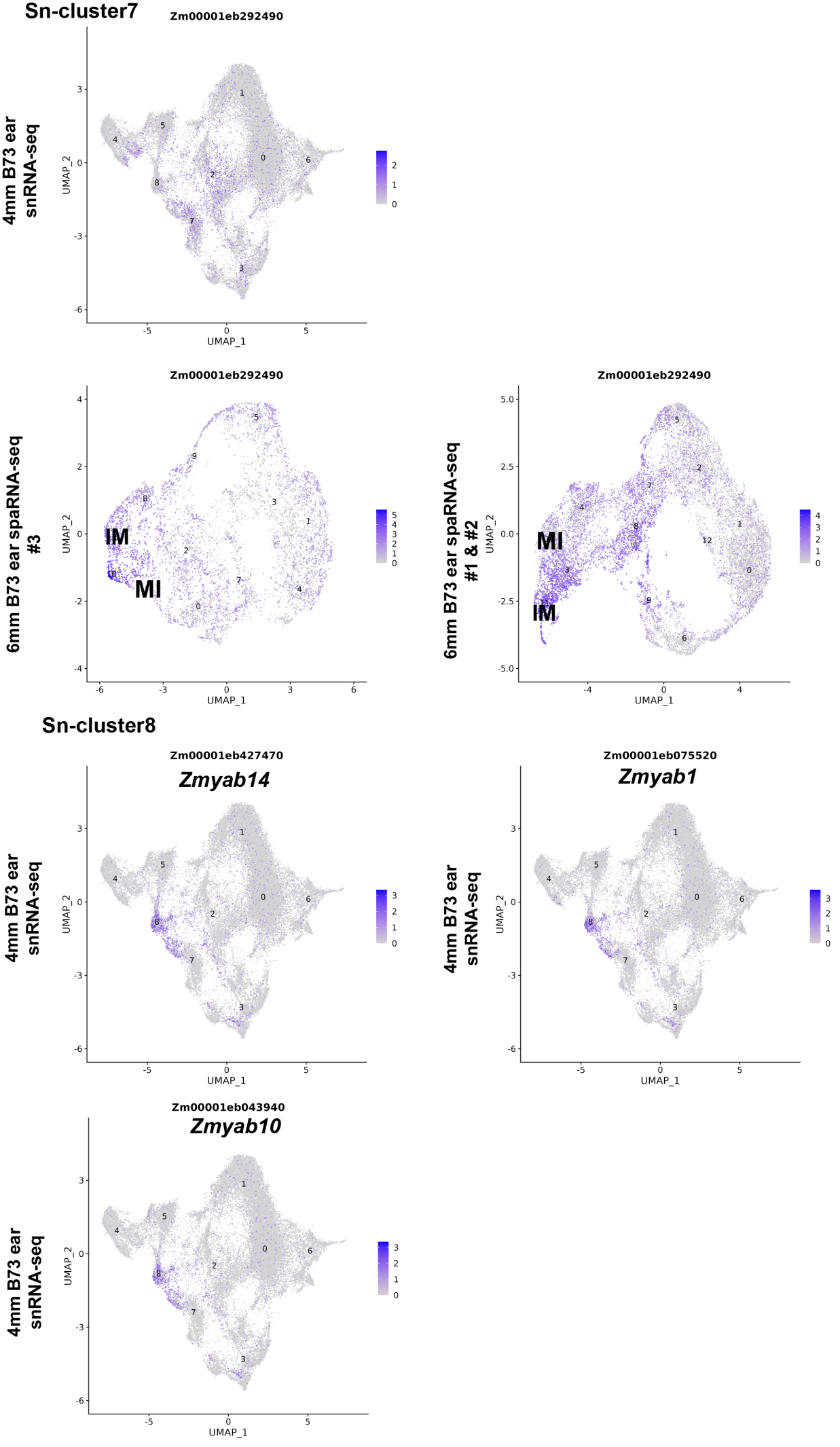

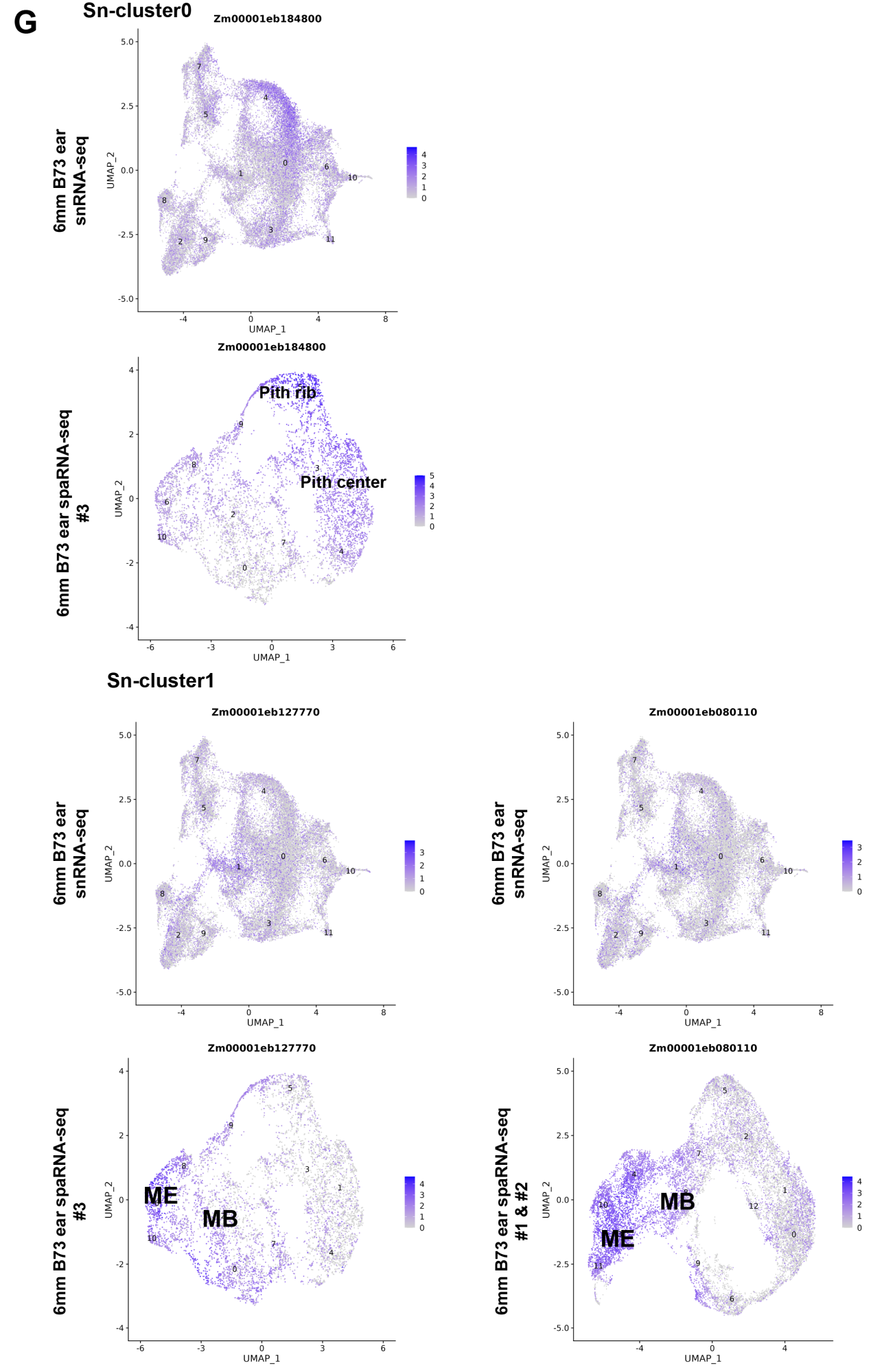

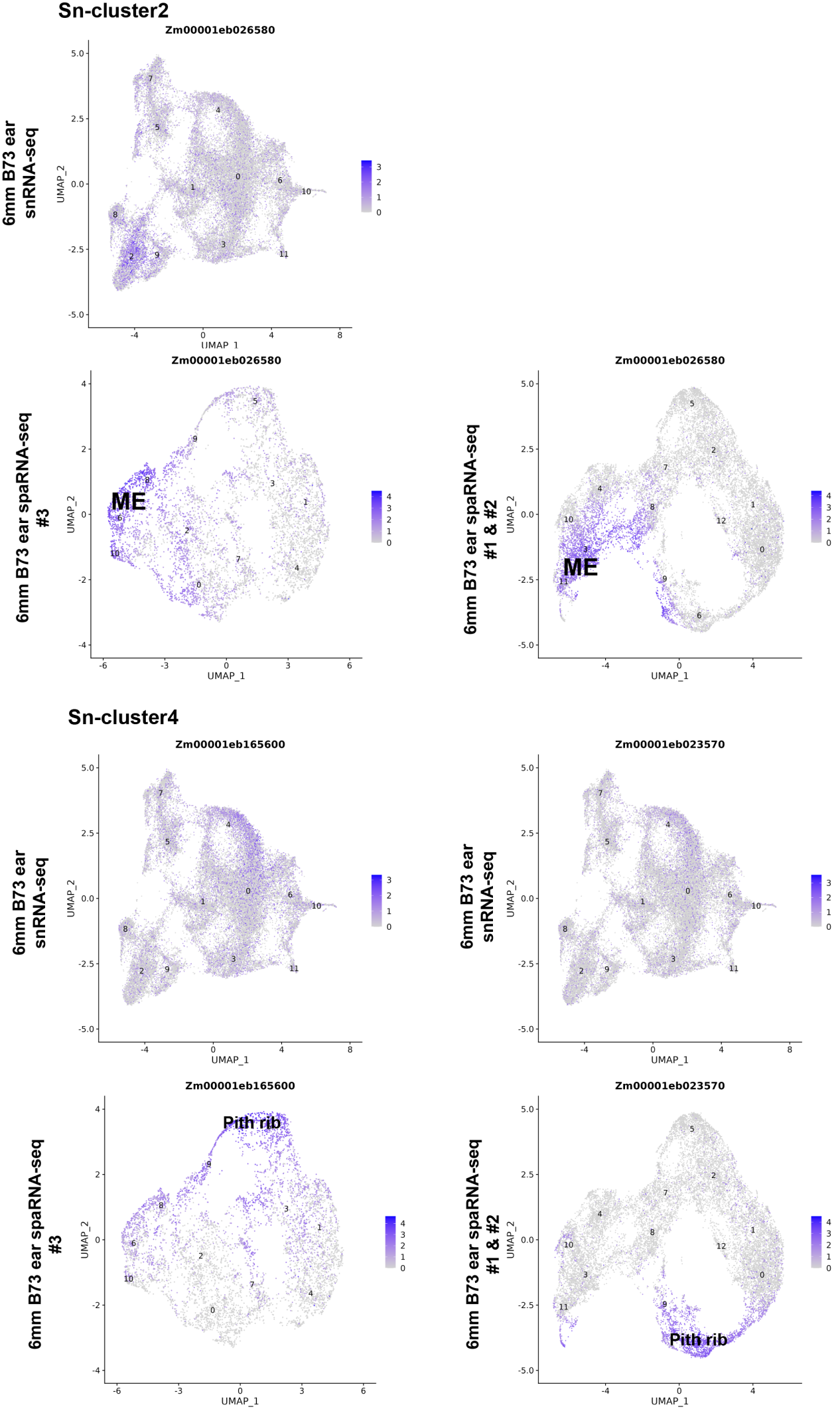

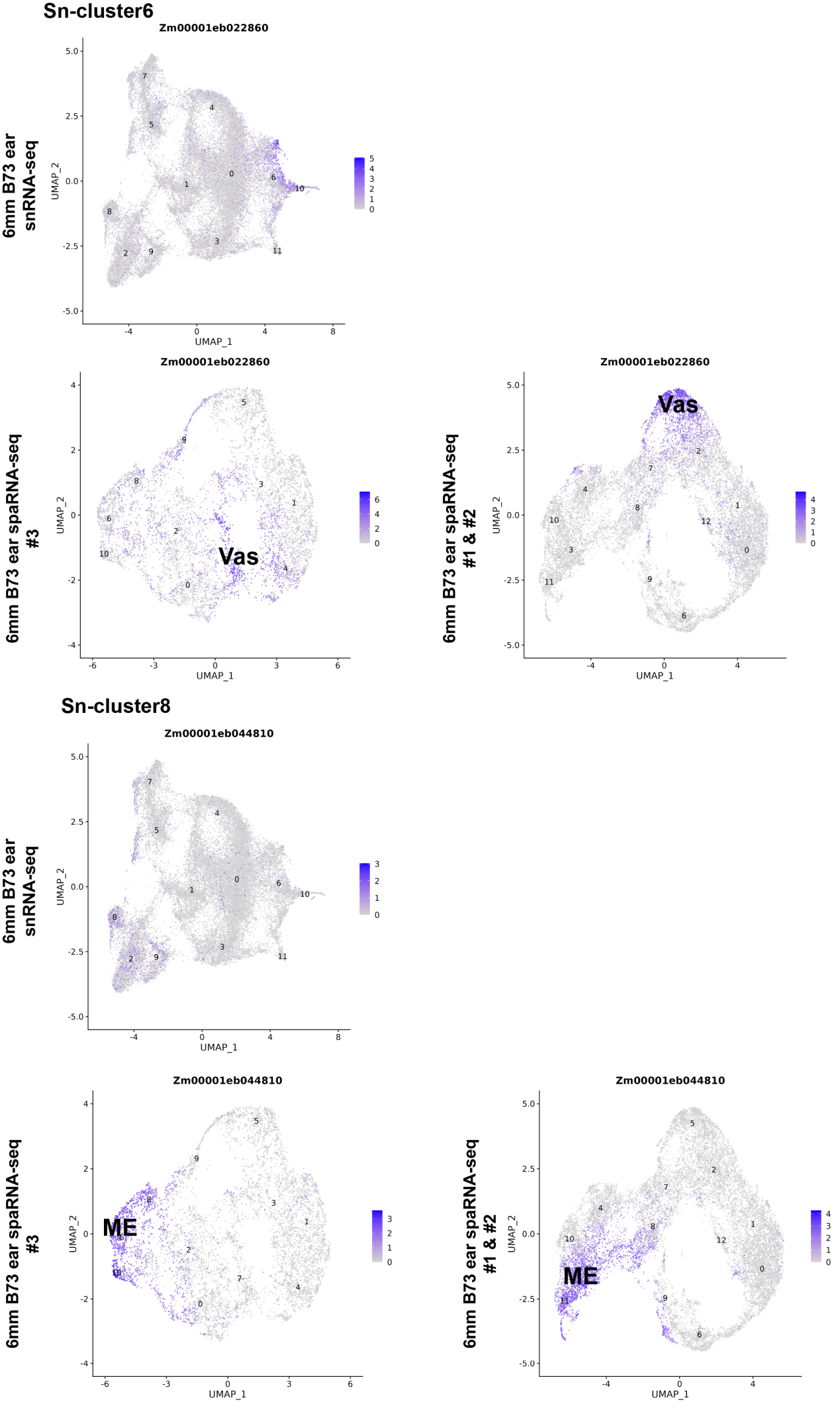

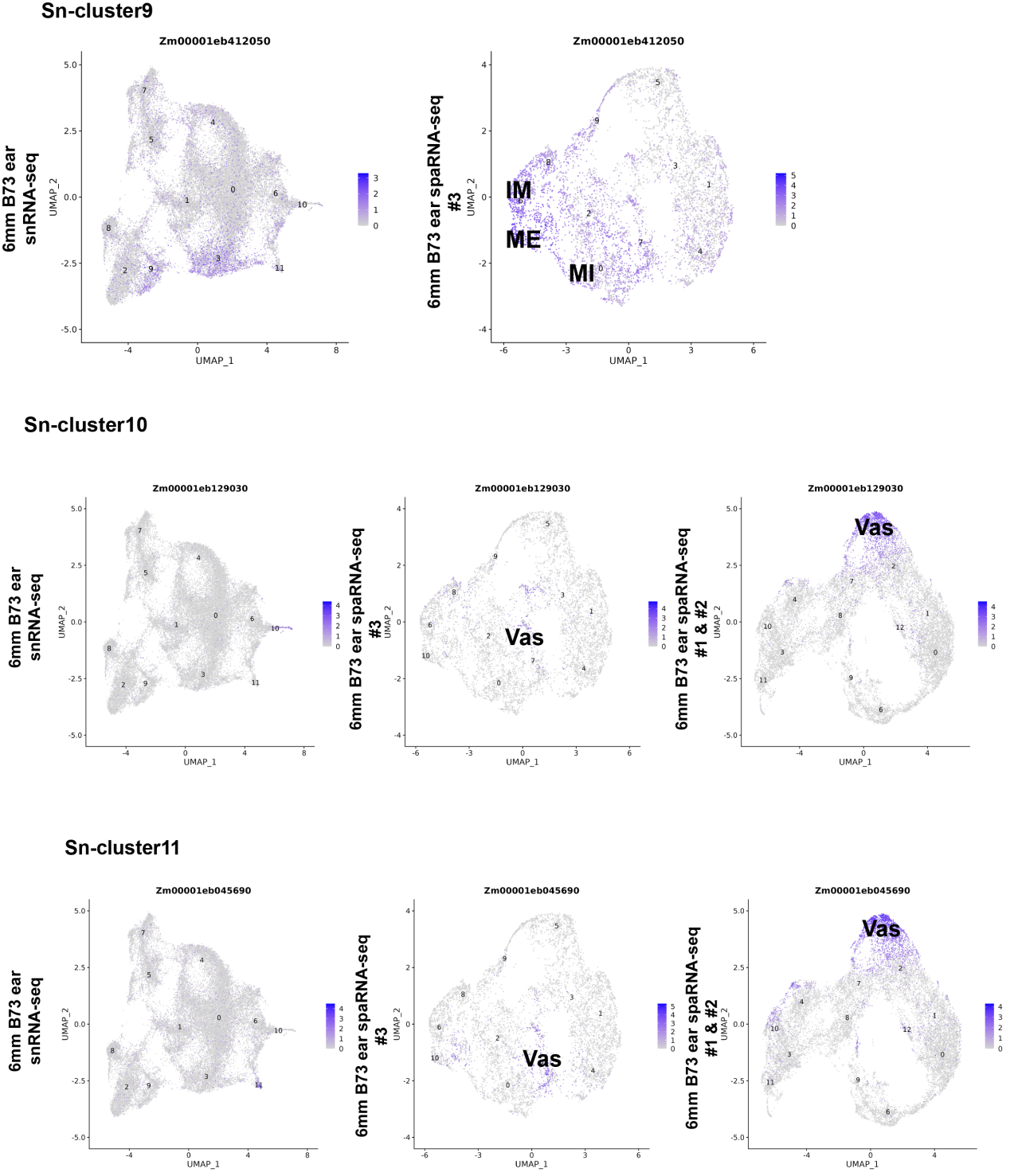

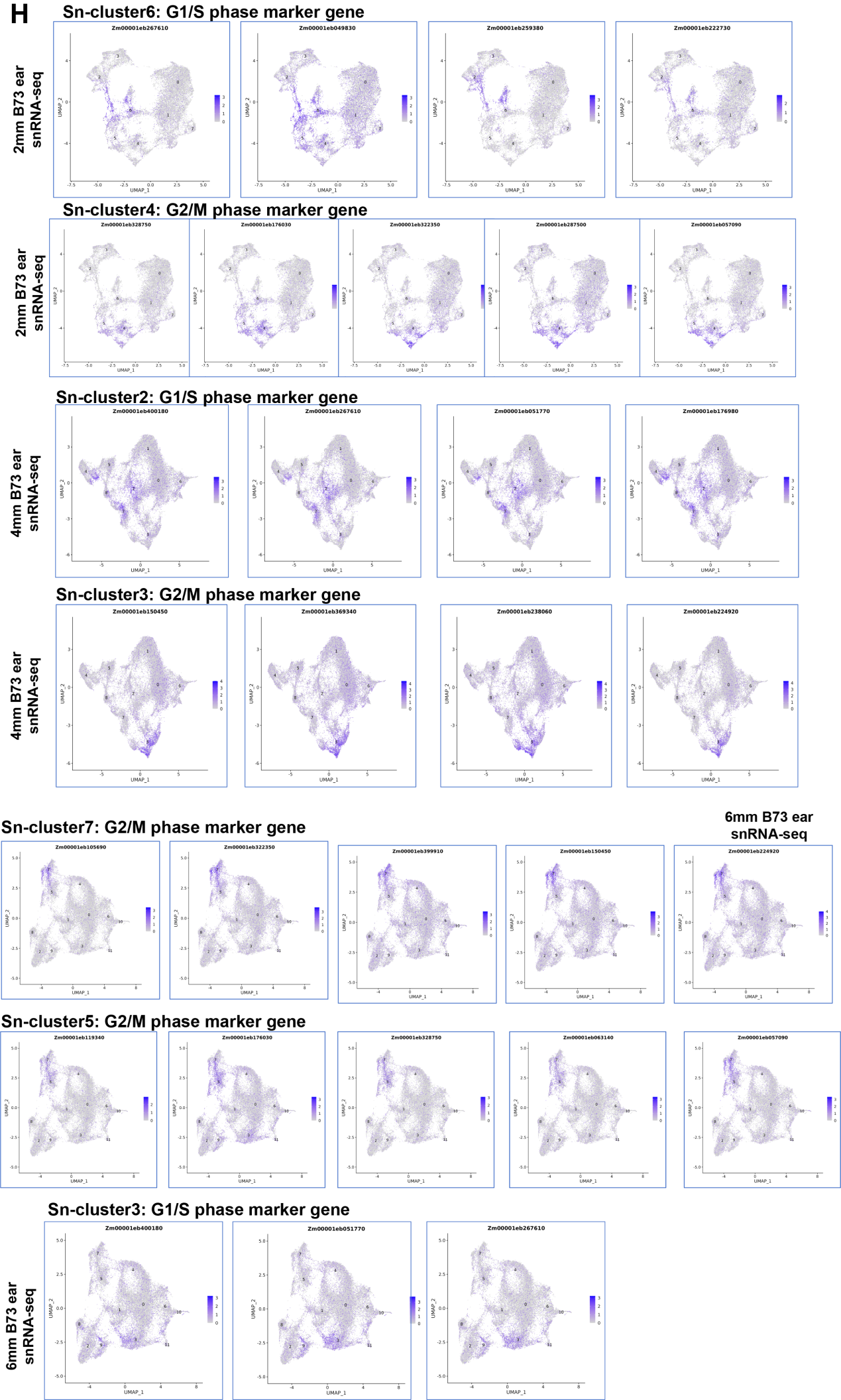

**Fig. S2 UMAP plots of marker genes predicting the identities of Sn-Clusters. (A)** Twelve clusters are displayed in an integrated UMAP plot in two dimensions. Each dot represents a bin50 from published maize spatial transcriptome data section #1 and #2 (*1*). **(B)** Spatial distribution of cell types that were identified in sections #1 and #2. **(C)** Eleven clusters are displayed in an integrated UMAP plot in two dimensions. Each dot represents a bin50 from published maize spatial transcriptome data section #3 (*1*). **(D)** Spatial distribution of cell types that were identified in section #3. **(E-G)** 2mm (E), 4mm (F), and 6mm (G) B73 snRNA-seq represent UMAP plots of their marker genes; the numbers in the figures indicate that the gene is the marker gene of the corresponding Sn-Cluster. 6mm B73 ear spaRNA-seq #1& #2 represent UMAP plots of the same genes in the published maize spatial transcriptome map shown in **(A-B),** 6mm B73 ear spaRNA-seq #3 represents UMAP plots of the same genes in the published maize spatial transcriptome map shown in **(C-D),** with color scale indicating normalized expression level. **(H)** Cell cycle marker genes from (*2*) used to annotate G2/M or S phase cell types.

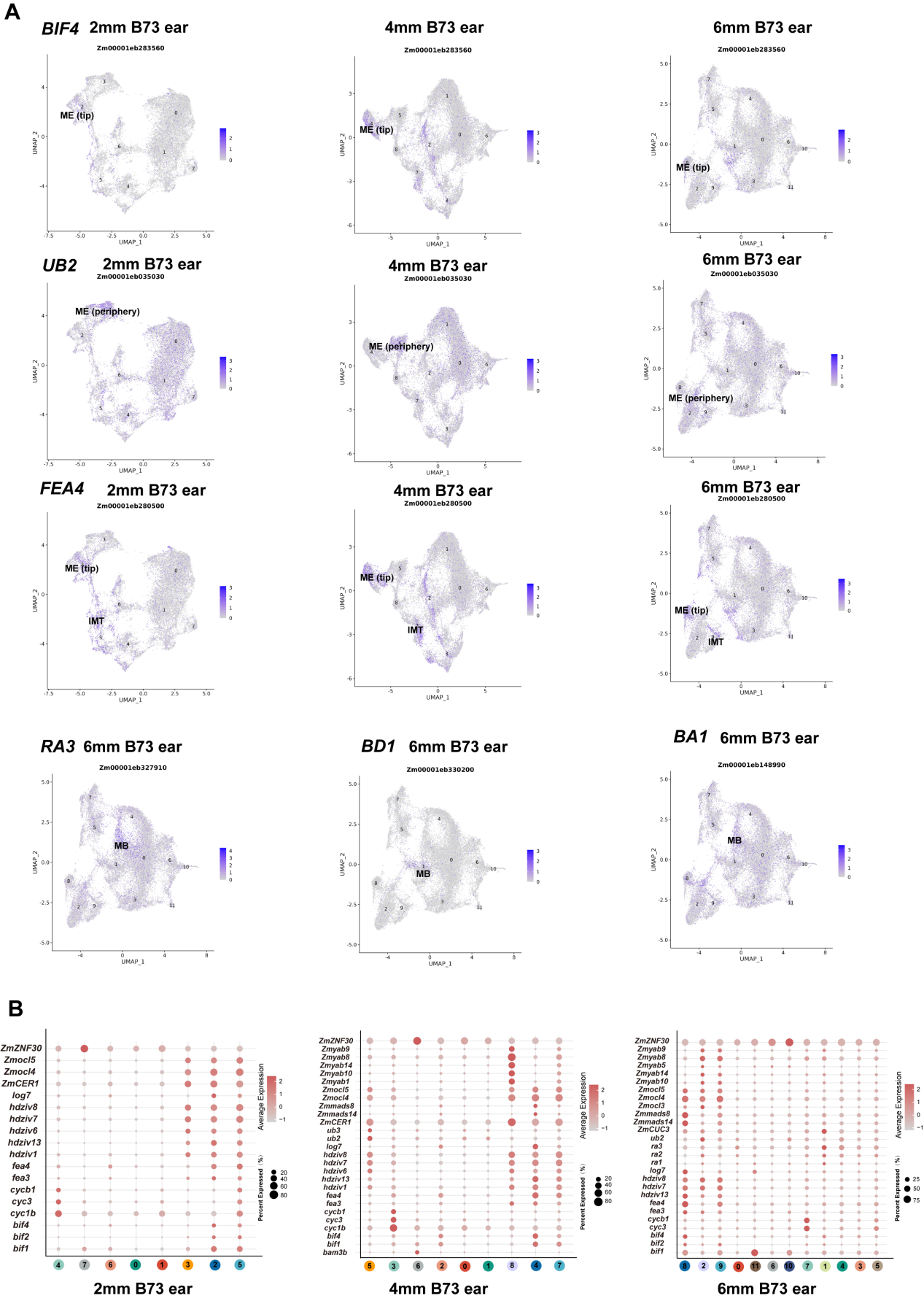

**Fig. S3 UMAP plots of marker genes predicting the identities of Sn-Clusters. (A)**. UMAP plots of known marker genes; the text in the figures indicates that the gene is the marker gene of the corresponding cell types with published mRNA in situ hybridization results, with the color scale indicating normalized expression level in snRNA-seq. **(B)**. Expression pattern of representative cluster-specific known marker genes. Dot diameter, proportion of cluster cells expressing a given gene. All known gene annotations were retrieved from (*3*)

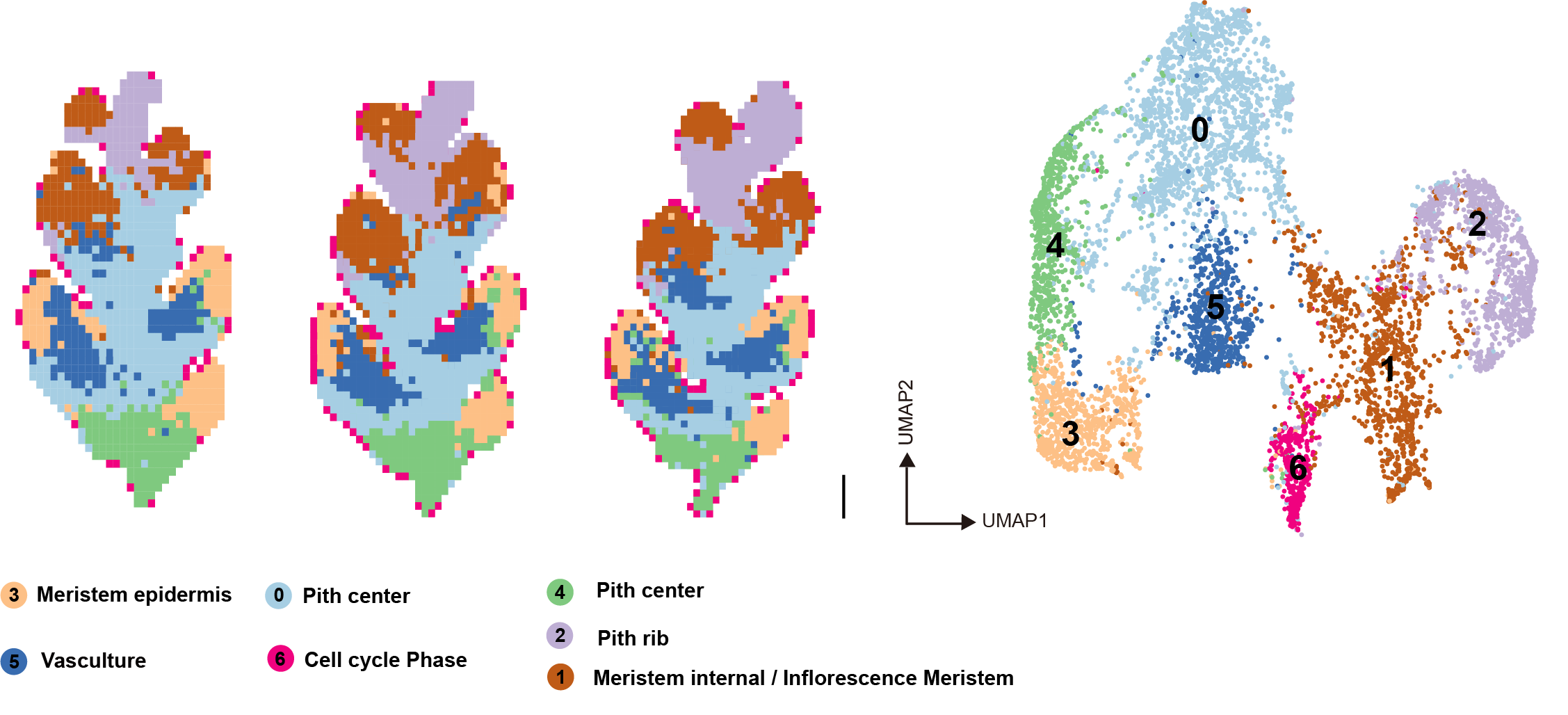

**Fig. S4 Unsupervised clustering of three teosinte ear sections based on imputed Stereo-seq data.** Left, Physical distribution of bin50s corresponding to different clusters. Scale bar = 0.1mm. Right, 7 meta-clusters displayed by an integrated UMAP plot in two dimensions, with each dot representing a bin50.

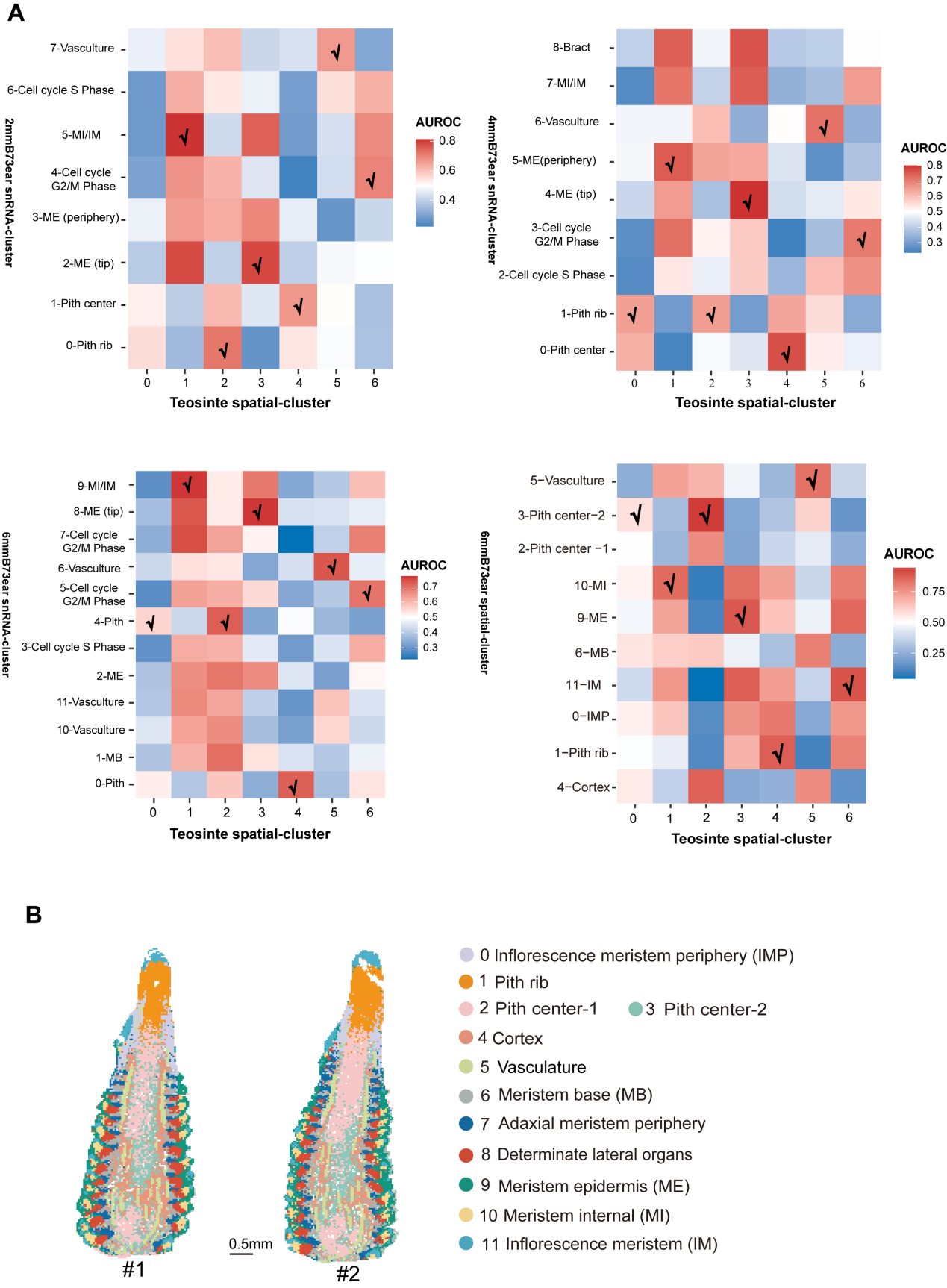

**Fig. S5 MetaNeighbor identifies cell types across teosinte snRNA-seq and maize snRNA-seq/spatial transcriptome data.** **(A)** Labels show the cell types with the highest identity (AUROCs > 0.6). **(B)** Spatial distribution of cell types that were identified in spatial transcriptomics of the 6 mm B73 ear (*1*). Cell types in (A) are shown in (B).

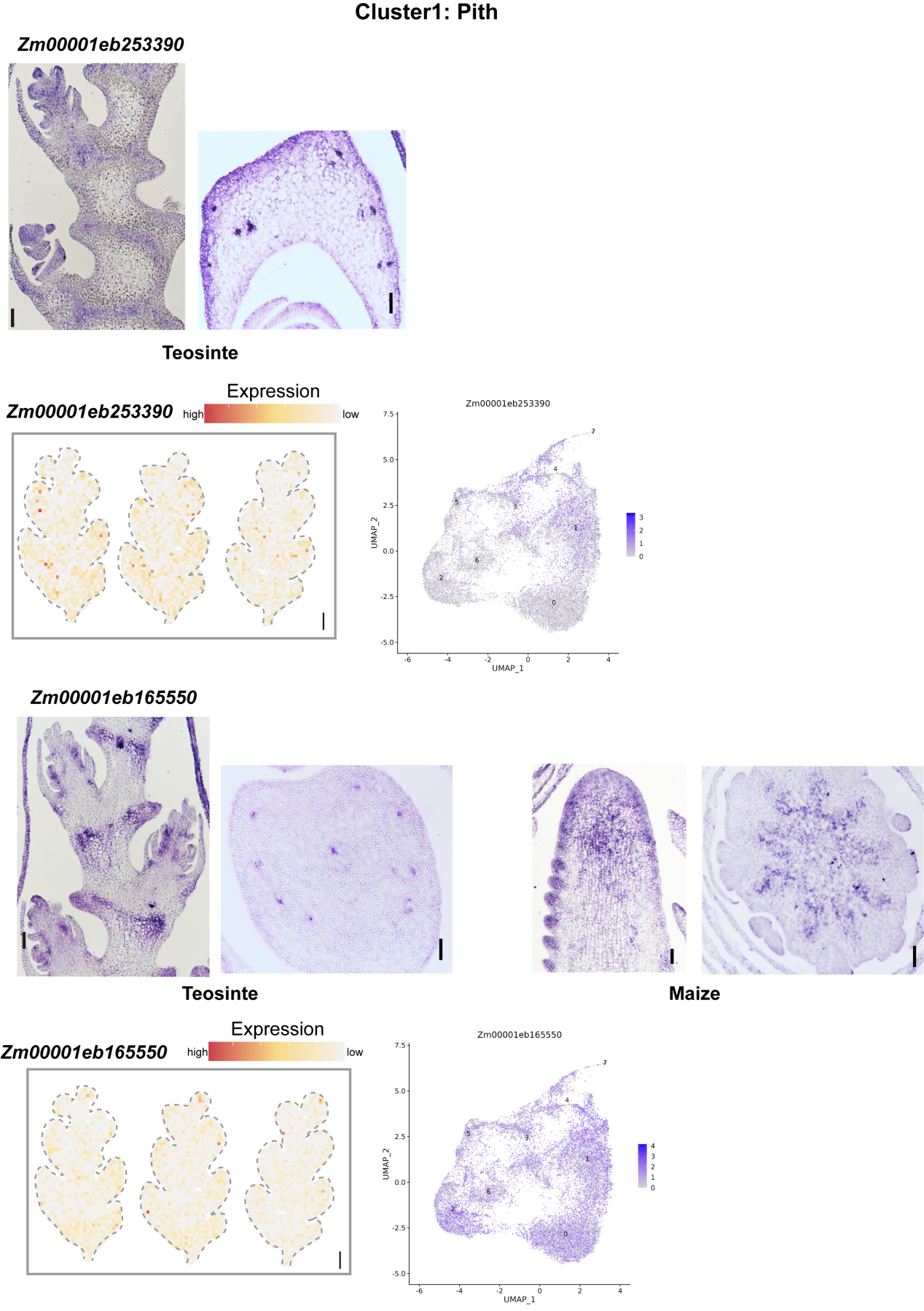

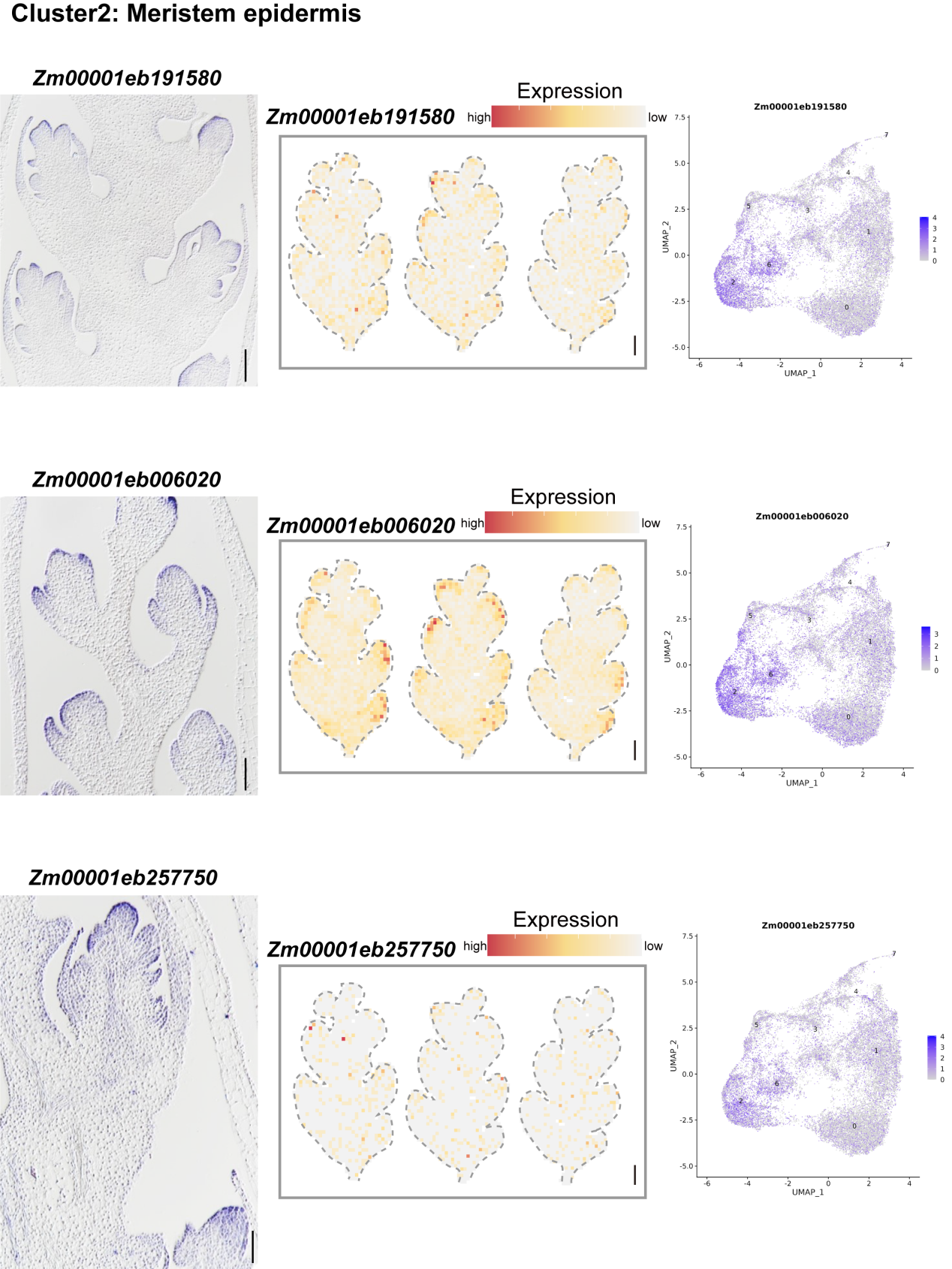

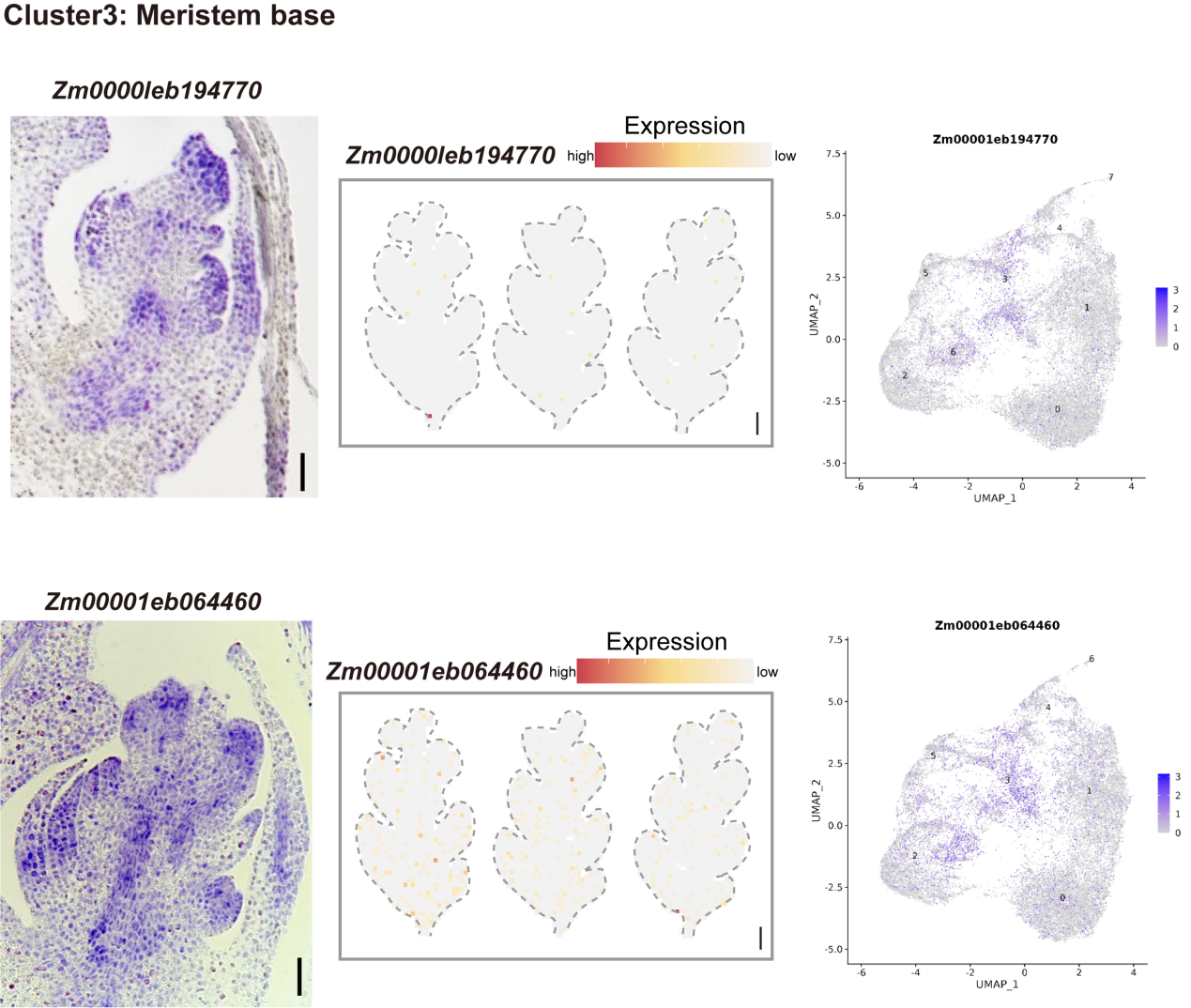

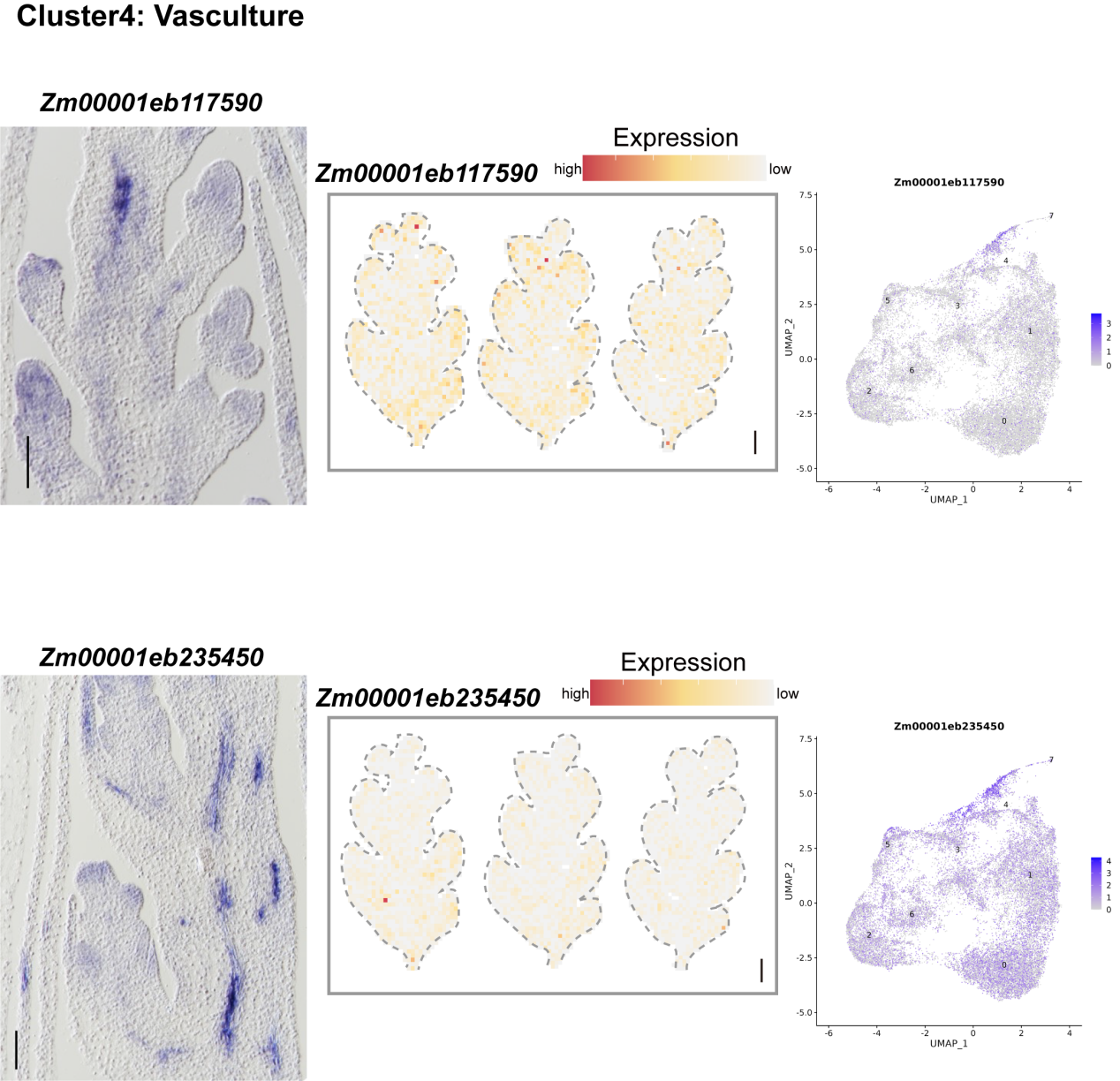

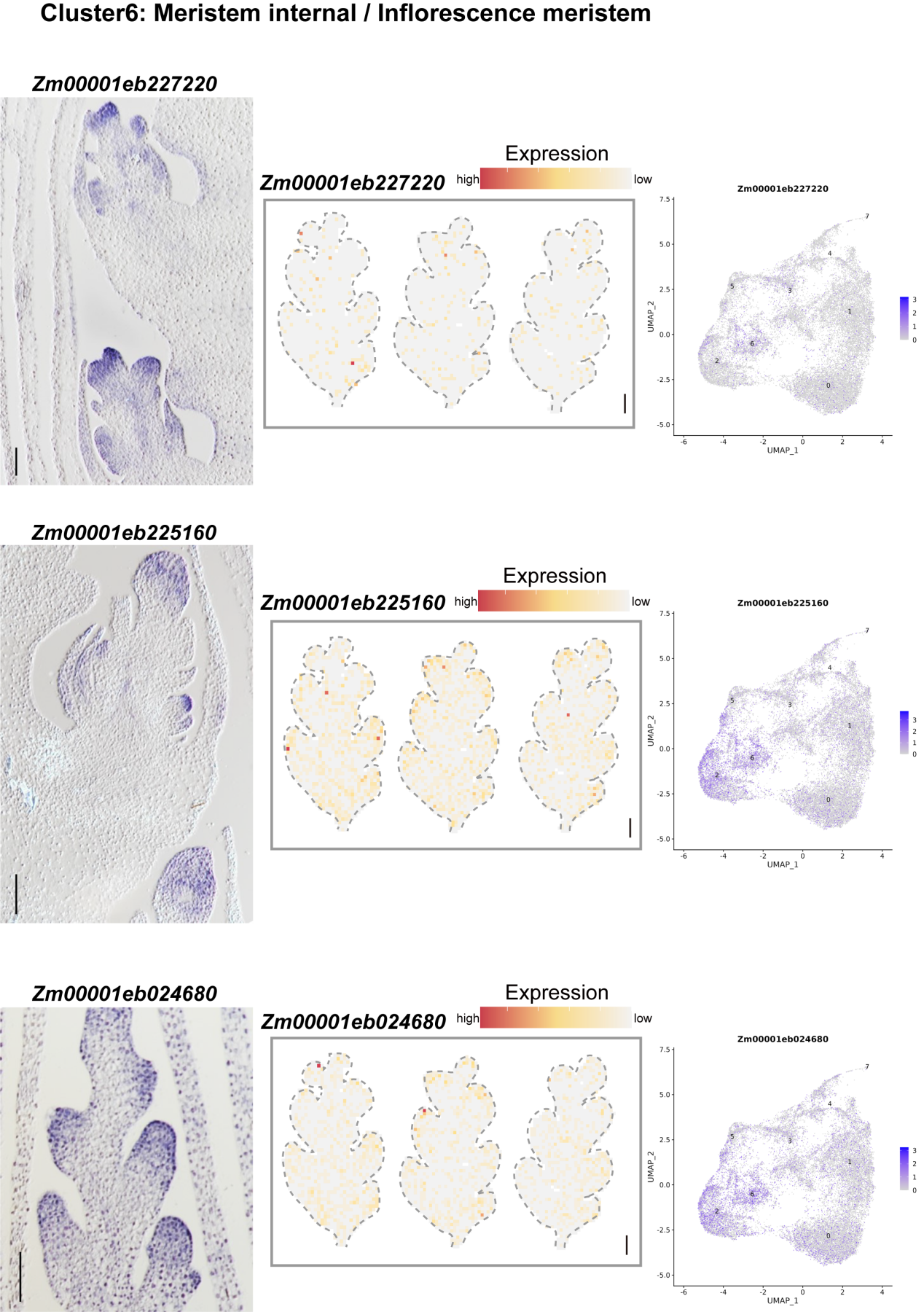

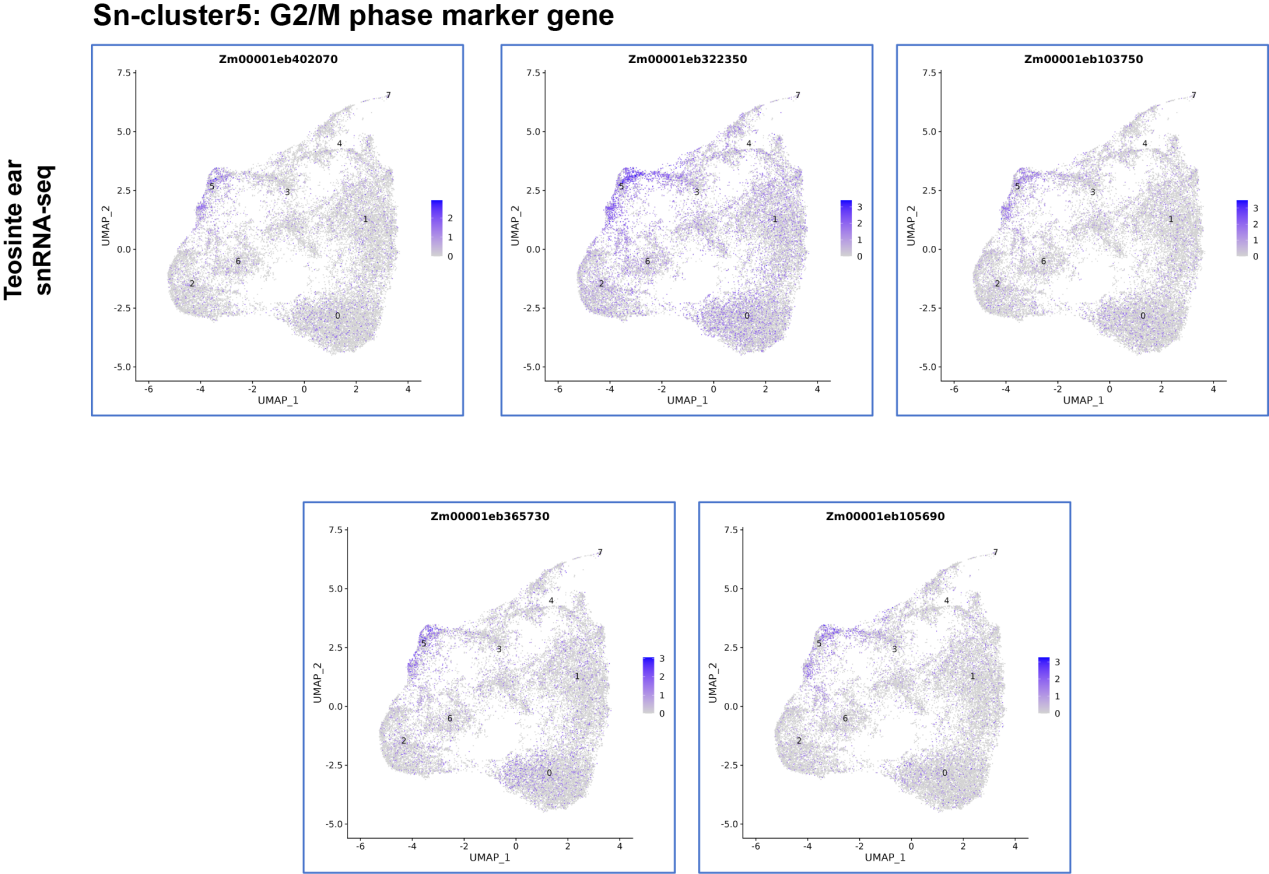

**Fig. S6 mRNA *in situ* hybridization of marker genes identified in teosinte snRNA-seq.** Left, mRNA *in situ* hybridization result. Middle, Stereo-seq expression pattern of teosinte ear, scale bar = 0.1 mm. Right, snRNA-seq expression pattern. The color scale indicates normalized expression level in snRNA-seq. Sn-cluster5: Cell cycle marker genes from (*2*) used to annotate G2/M phase cell types.

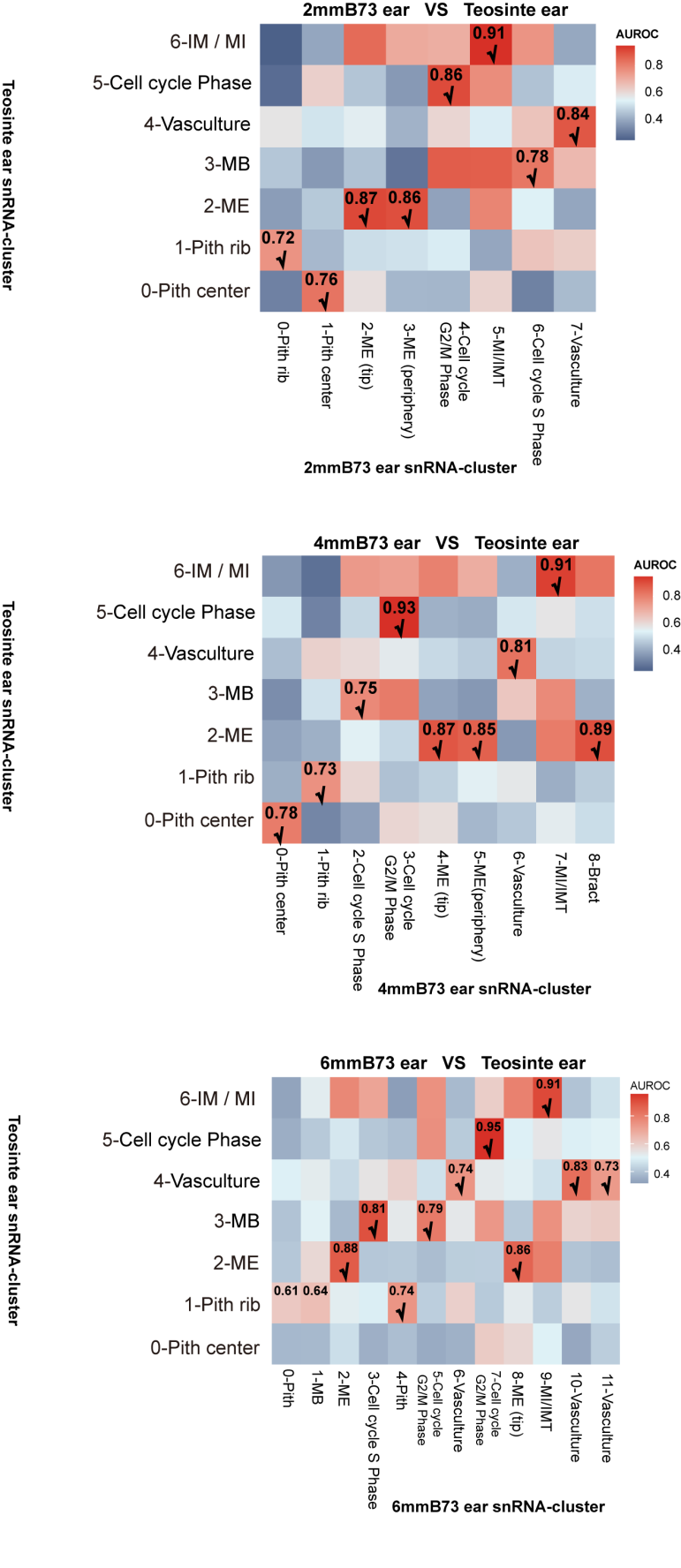

**Fig. S7 MetaNeighbor identifies cell types across teosinte snRNA-seq and maize snRNA-seq datasets.**

Values indicate AUROC scores reflecting cross-species cell type similarity. Cob-related cell types exhibit lower similarity between maize and teosinte compared to floret-related cell types.
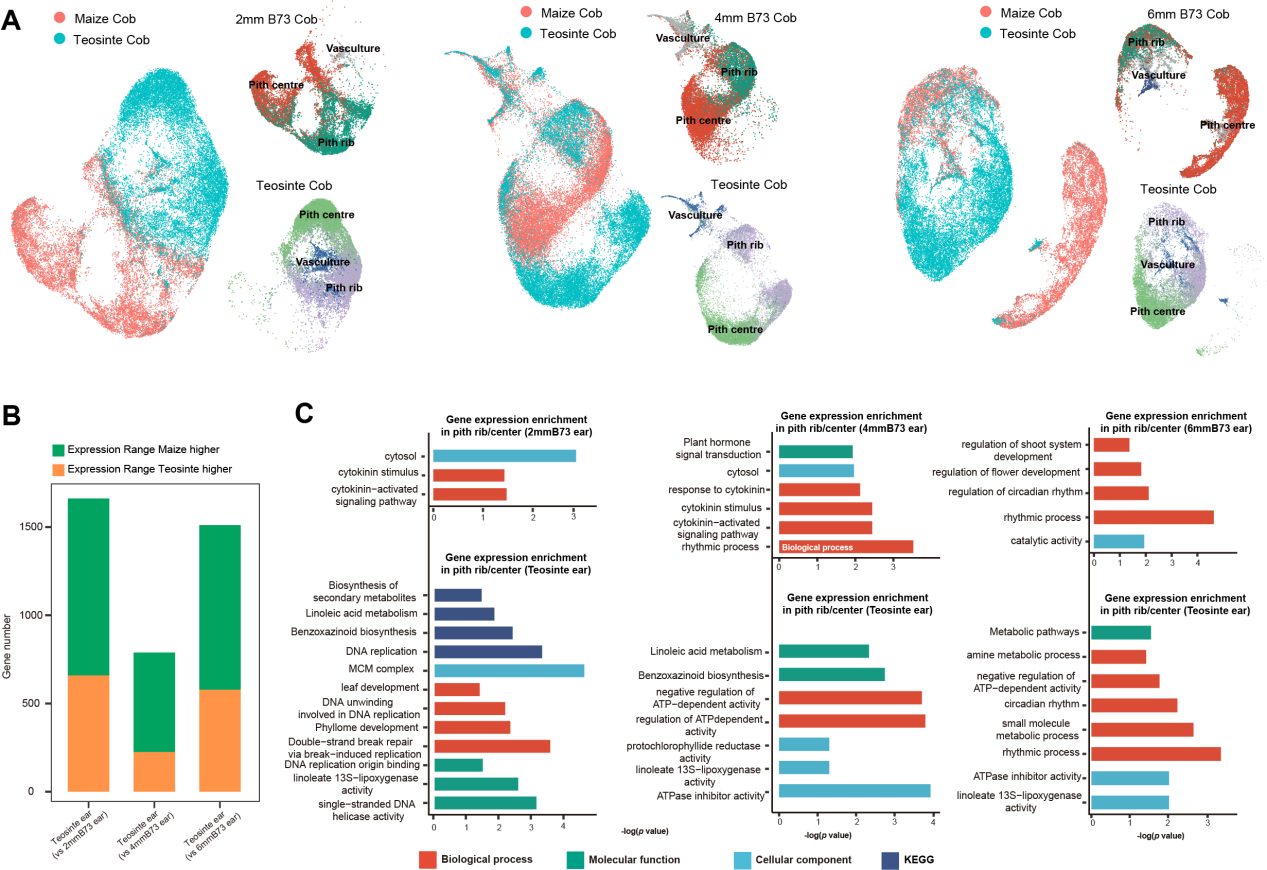

#### Fig. S8 Transcriptomic divergence for cob cells. (A) Integrated transcriptomic comparison of cob cell populations between maize (three developmental stages: 2mm, 4mm, and 6 mm) and teosinte, including pith rib zone, pith center, and vasculature. (B) Number of differentially expressed genes (DEGs) identified between teosinte and maize across three developmental stages (2mm, 4mm, and 6 mm) within specific pith sub-populations. (C) Bar graph showing Gene Ontology (GO) enrichment of maize and teosinte enriched genes in pith cells, with maize sampled at the 2mm, 4 mm, and 6mm stages.

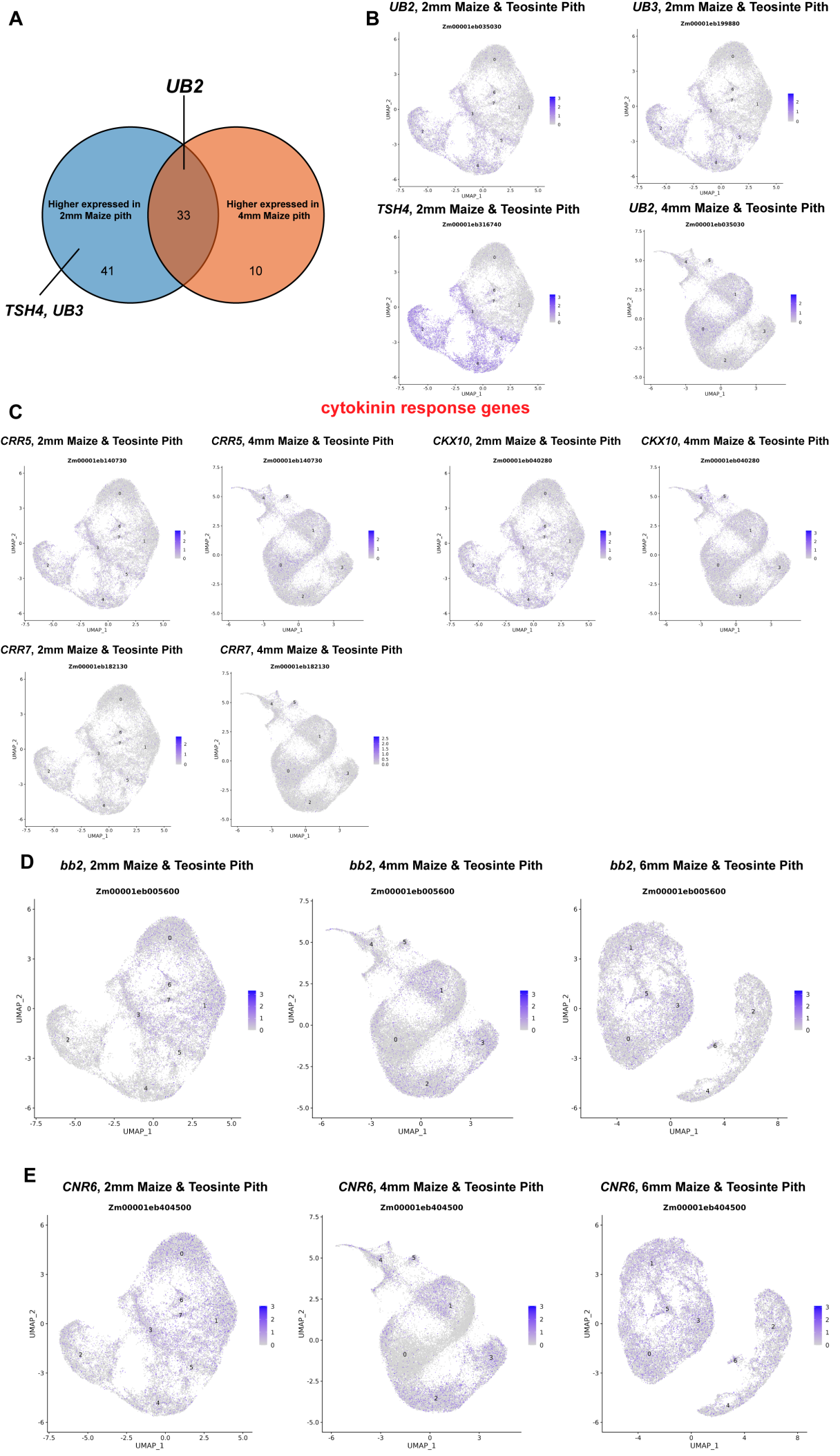

**Fig. S9 Enrichment and expression patterns of selected domestication and cytokinin-related genes in maize pith cells. (A)** Number of genes that are highly expressed in maize 2 mm and 4 mm pith cells and exhibit signatures of selection. **(B-C)** Integrated UMAP visualization of known domestication genes identified in (A), with color scales representing normalized expression levels across 2mm, 4mm, and teosinte snRNA-seq datasets. **(D)** Integrated UMAP visualization of cytokinin-responsive genes, with color scales indicating normalized expression levels. **(E)** Integrated UMAP visualization of *bb2*, with color scales indicating normalized expression levels. **(F)** Integrated UMAP visualization of *CRN6*, with color scales indicating normalized expression levels.

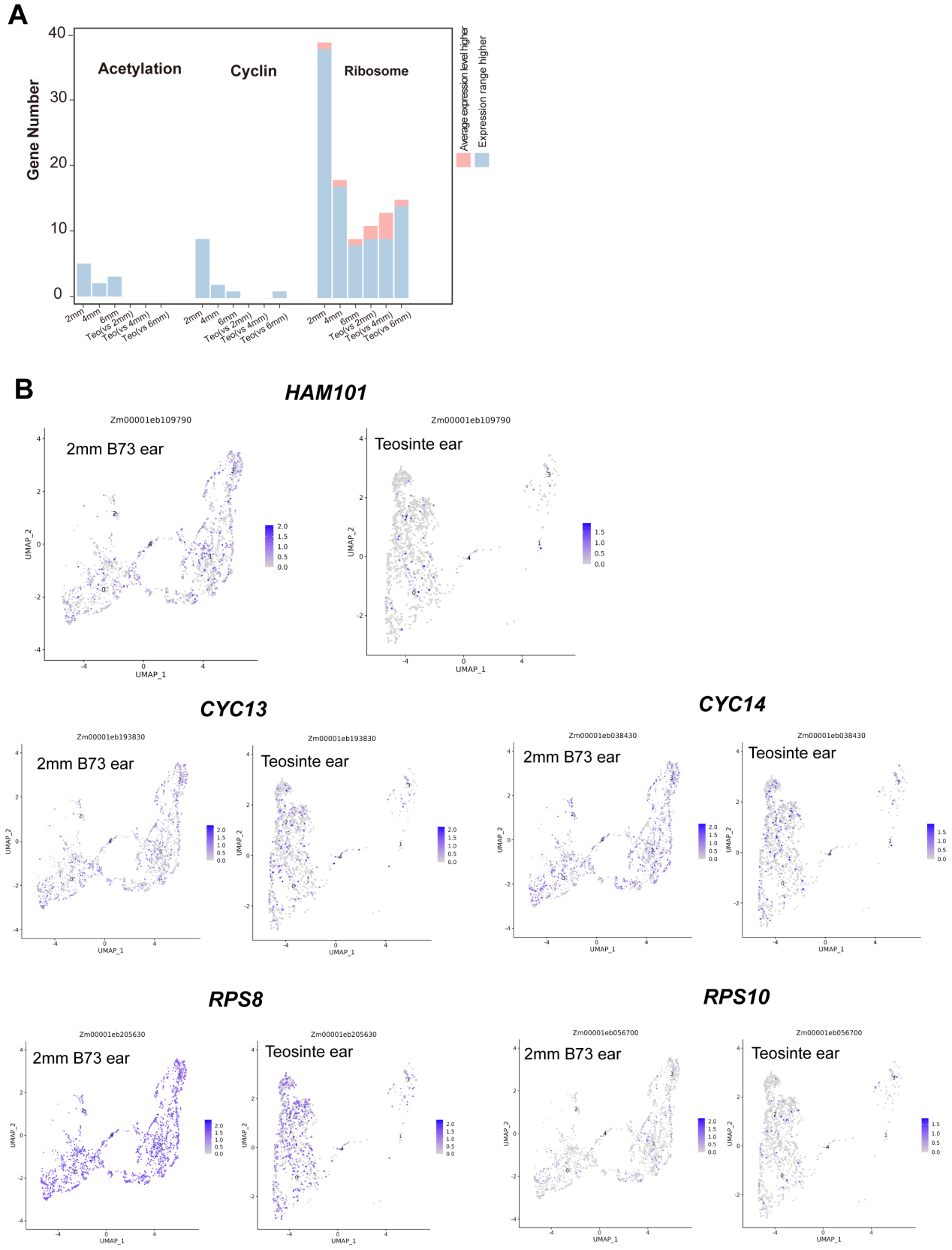

**Fig. S10 Genes within the maize-specific module (M5) were enriched for three major cell cycle related processes.** (A) Number of differentially expressed genes (DEGs) associated with cell cycle related processes, identified between teosinte and maize across three developmental stages (2mm, 4mm, and 6 mm) within IM subpopulations. (B) Left, UMAP plots of cell cycle related genes, with color scale indicating normalized expression level in maize and teosinte snRNA-seq.

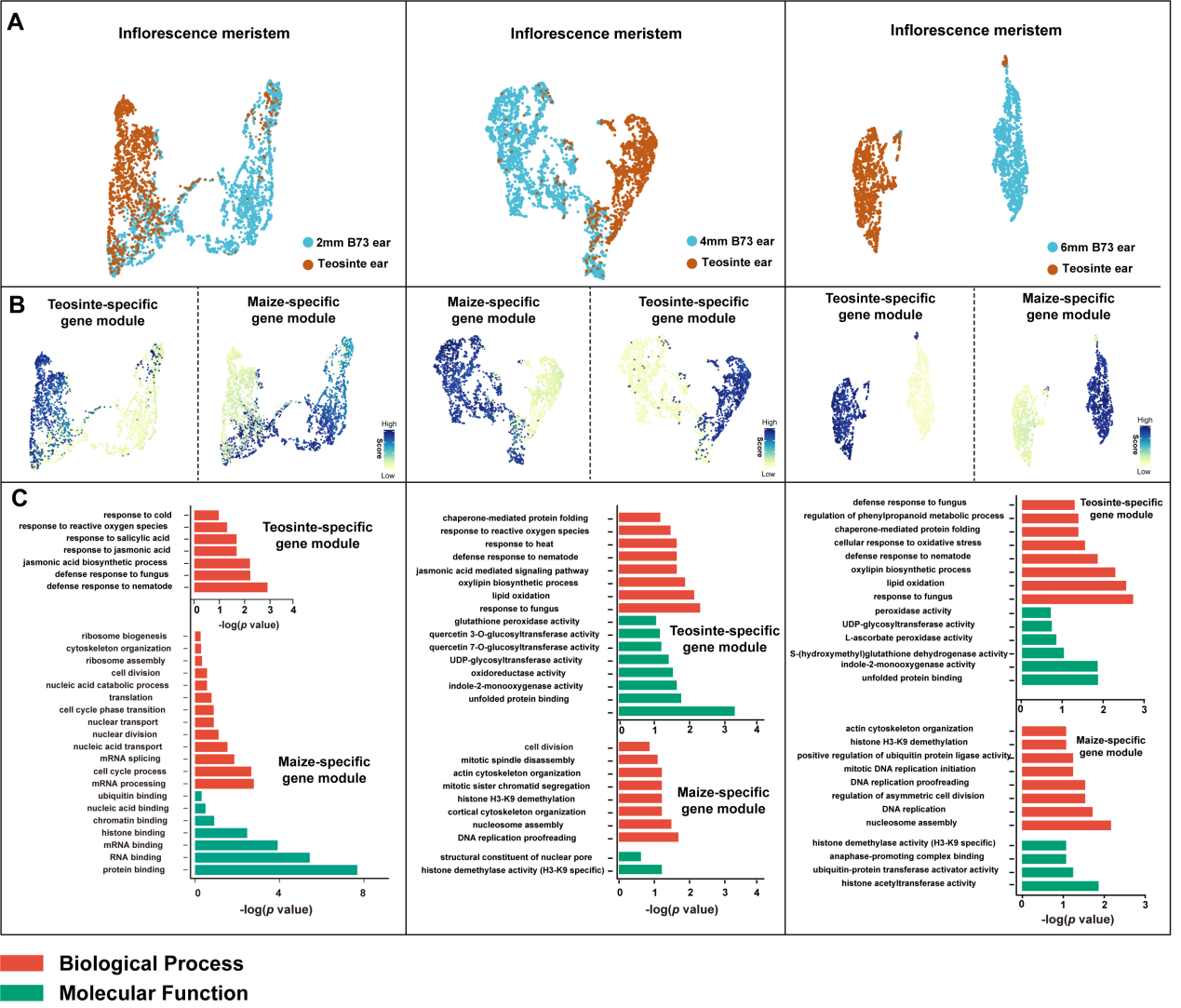

**Fig. S11 Comparison of IM in maize and teosinte ears. (A)** UMAP visualization and clustering of IM cells in maize and teosinte. **(B)** Hotspot analysis and identification of gene modules within isolated IM cells. Gene modules with higher hotspot scores indicate stronger co-expression patterns. **(C)** GO enrichment analysis of the maize-specific and teosinte-specific IM gene modules.

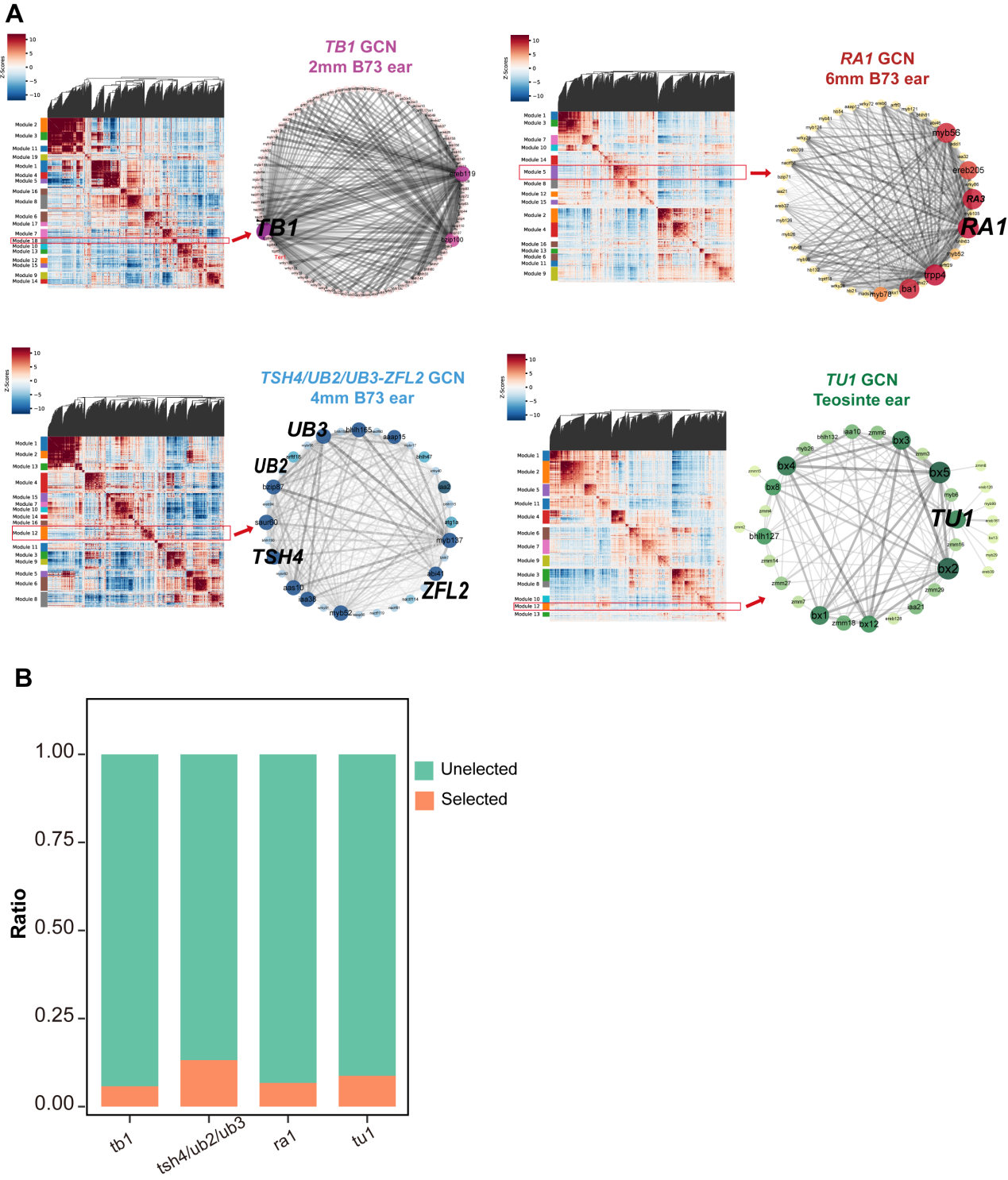

**Fig. S12 Constructing spatial co-expression networks.** (A) Left, Hotspot gene module analysis and identification of gene modules within isolated meristem cells. Right, gene co-expression network (GCN) constructed from a gene module containing TB*1*, RA*1*, TSH*4/UB2/UB3-ZFL2*, and TU*1*. The network is visualized using Cytoscape. Node size represents the degree of connectivity. The shade of the line color represents the weight of the edge between two nodes. (B) The ratio of selected genes in the gene modules described in (A).

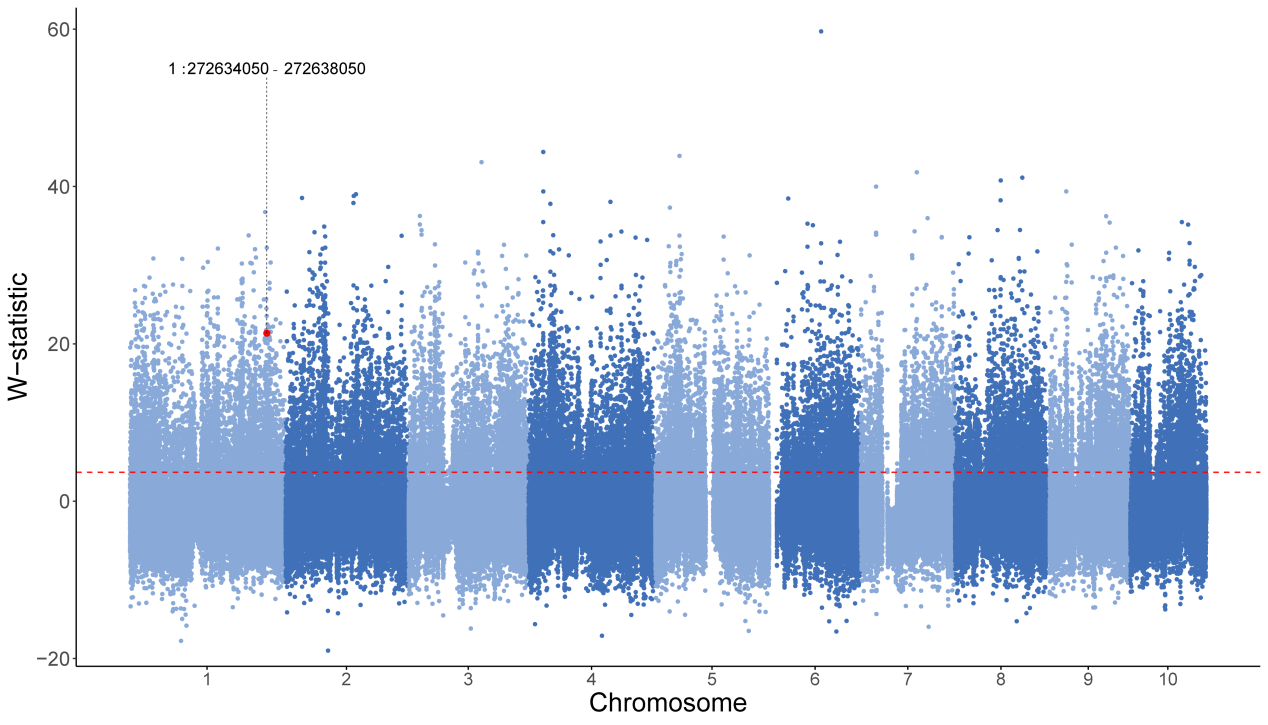

**Fig. S13 Genome-wide distribution of selection signatures (*W*-statistic) between *Zea mays* ssp. *parviglumis* and maize.** The *W*-statistic, derived from the smoothed Cross Population Extended Haplotype Homozygosity (XP-EHH), was calculated across the entire genome to identify regions potentially under selection during maize domestication. Each dot represents a genomic window, with chromosomal positions shown along the x‑axis and *W*‑statistic values on the y‑axis. The horizontal red dashed line indicates the genome‑wide significance threshold corresponding to the top 5% of empirical *W*‑statistic values (threshold = 3.68). Genomic windows exceeding this cutoff are considered as outliers with strong selection signals. The red dot highlights a specific genomic region on chromosome 1 (chr1:272,634,050–272,638,050).

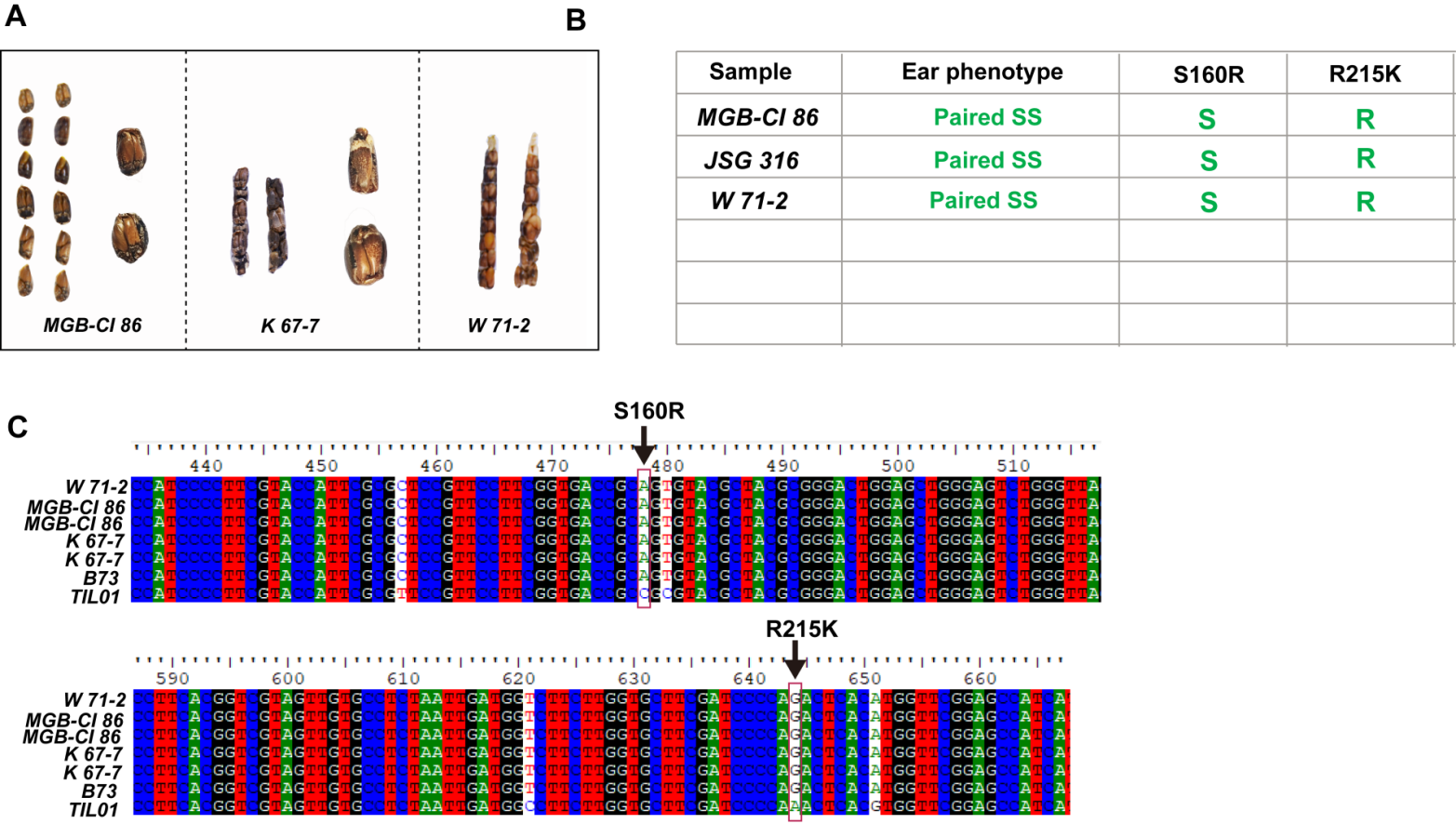

**Fig. S14 Teosinte accessions from wild populations that exhibited paired sessile spikelets. (A) Teosinte ear with double rows of SS, and detail of the paired SS. (B-C) Haplotype of *ZmSPD1* in teosinte with paired SS and sequence from TIL01 published genome from Maizegdb** (*4*)**. Upon translation of the full DNA sequence, the A/C mutation results in the substitution of Serine with Arginine. The G/A mutation results in the substitution of Arginine with Lysine.**

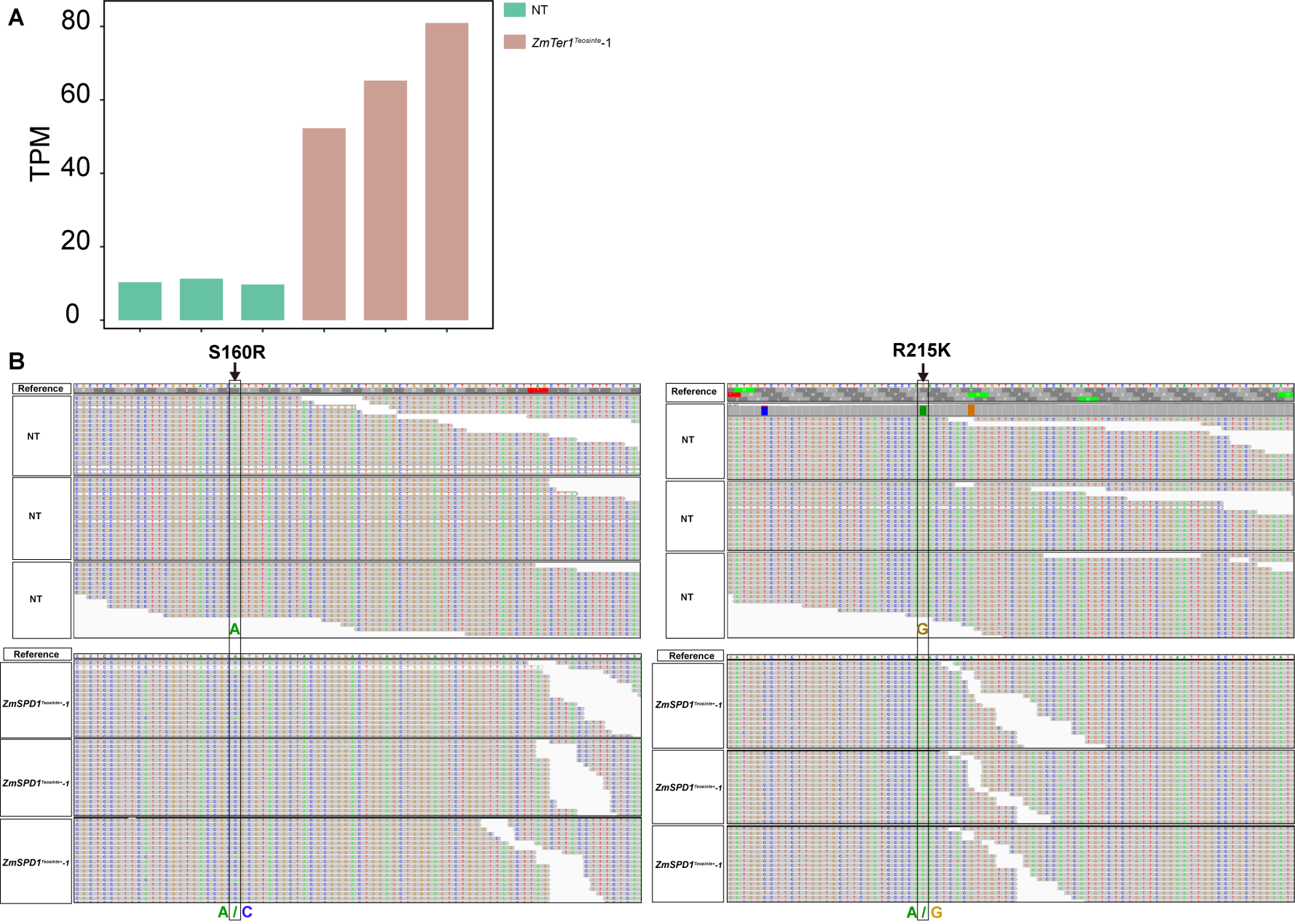

**Fig. S15 Transcriptomic validation of NT (non-transgenic line) and *ZmSPD1^Teo+^* lines. (A)** RNA-seq–based quantification of transcript abundance (TPM) in NT and ***ZmSPD1^Teosinte+^*** lines, shown as bar plots for the *Zm00001eb054470*. **(B)** IGV visualization of RNA-seq read coverage at the *Zm00001eb054470*. In the NT line, RNA-seq reads showed only the endogenous nucleotide at the diagnostic polymorphic site (A and G), whereas the ***ZmSPD1^Teosinte+^*** line displayed reads corresponding to both endogenous and introduced nucleotides (A/C and A/G), confirming successful expression of the introduced ***ZmSPD1^Teosinte+^*** transcript.

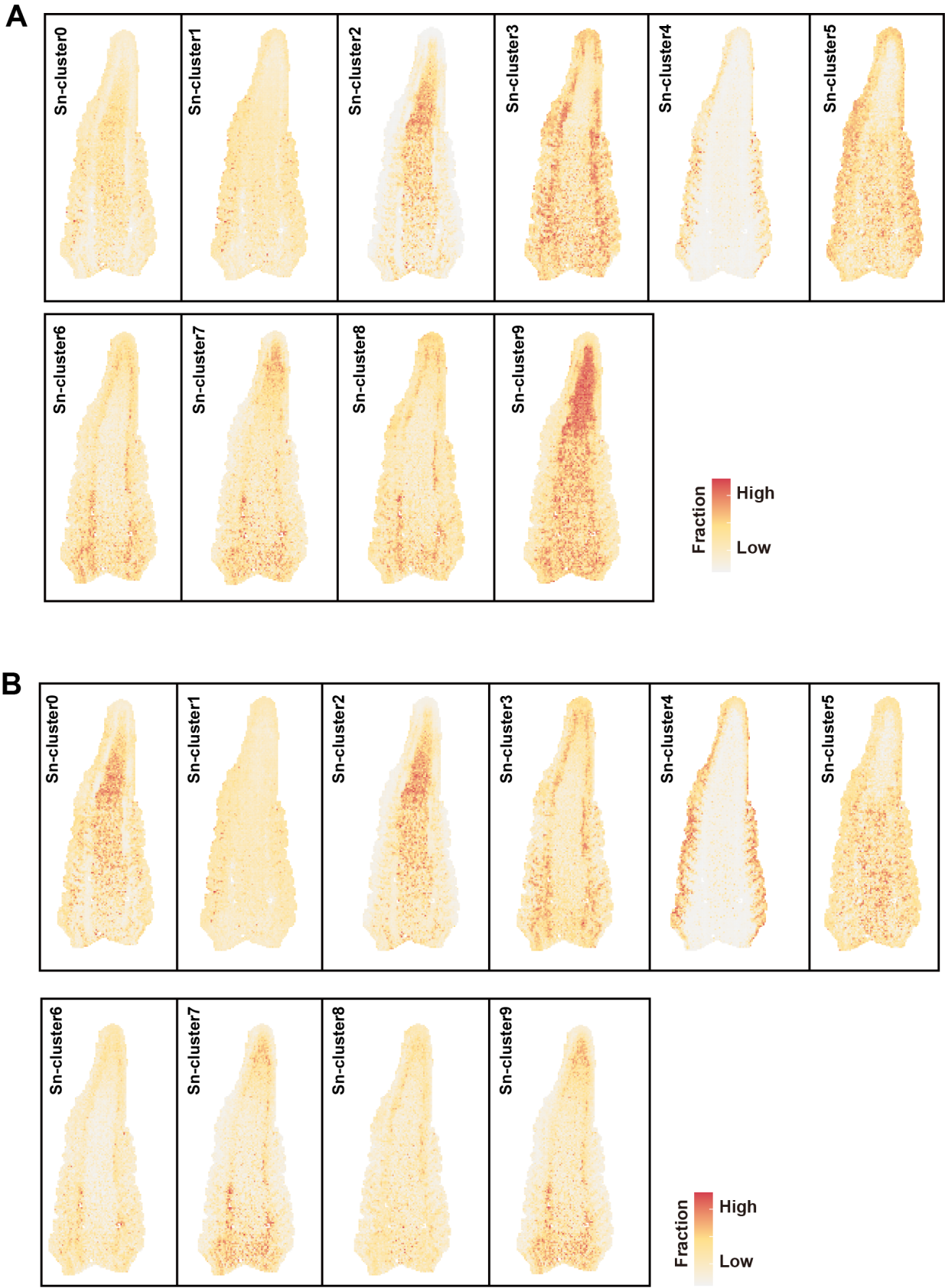

**Fig S16. STRIDE was applied to project the *Ter1*-NIL*^Teosinte^* and *Ter1*-NIL*^Maize^* Sn-Clusters onto the maize spatial transcriptome map (A-B).** Color scale indicates the proportion of cells from different Sn-clusters in the spots. Sn-clusters from *Ter1*-NIL*^Teosinte^* ear (A) and *Ter1*-NIL*^Maize^* ear (B).

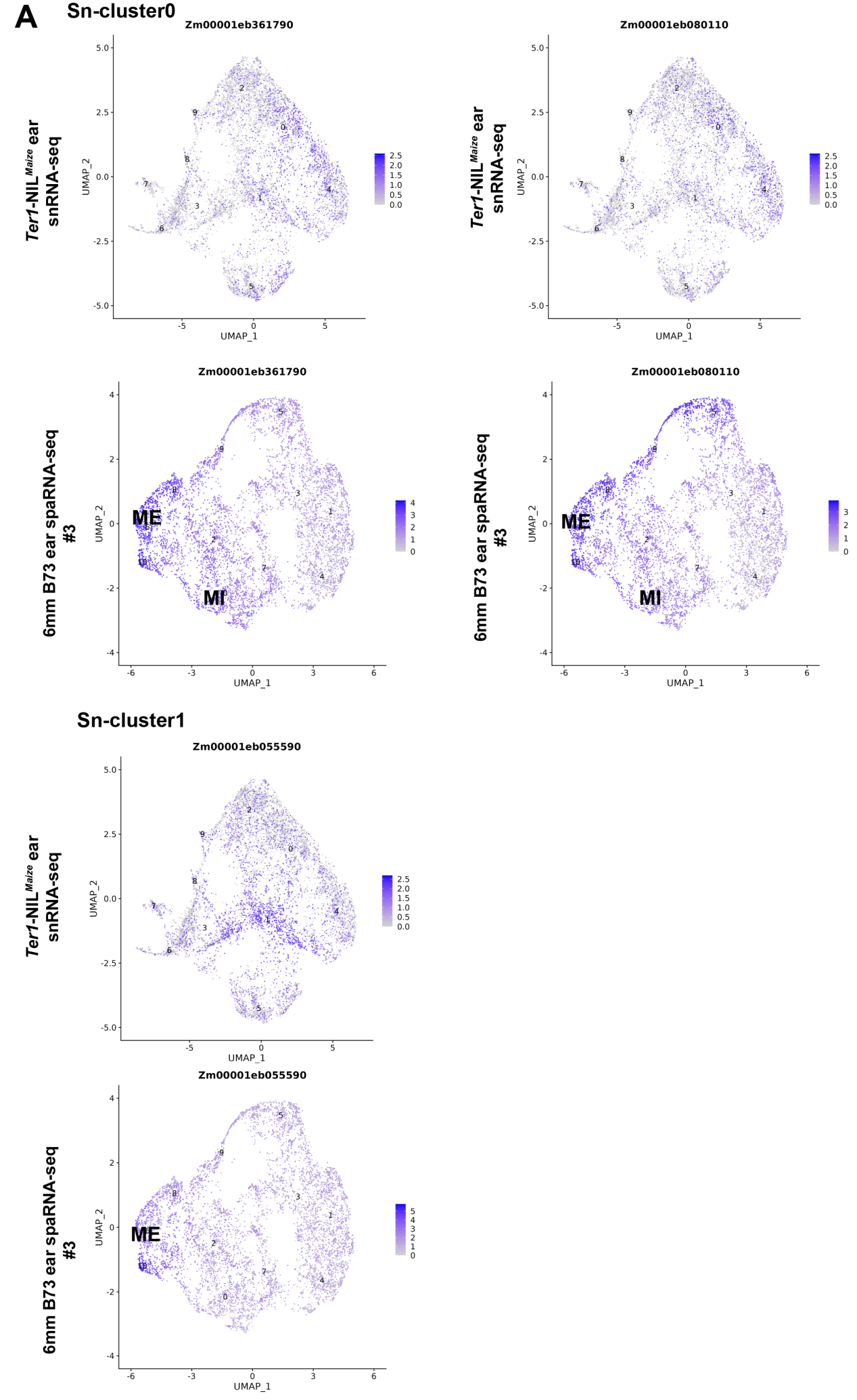

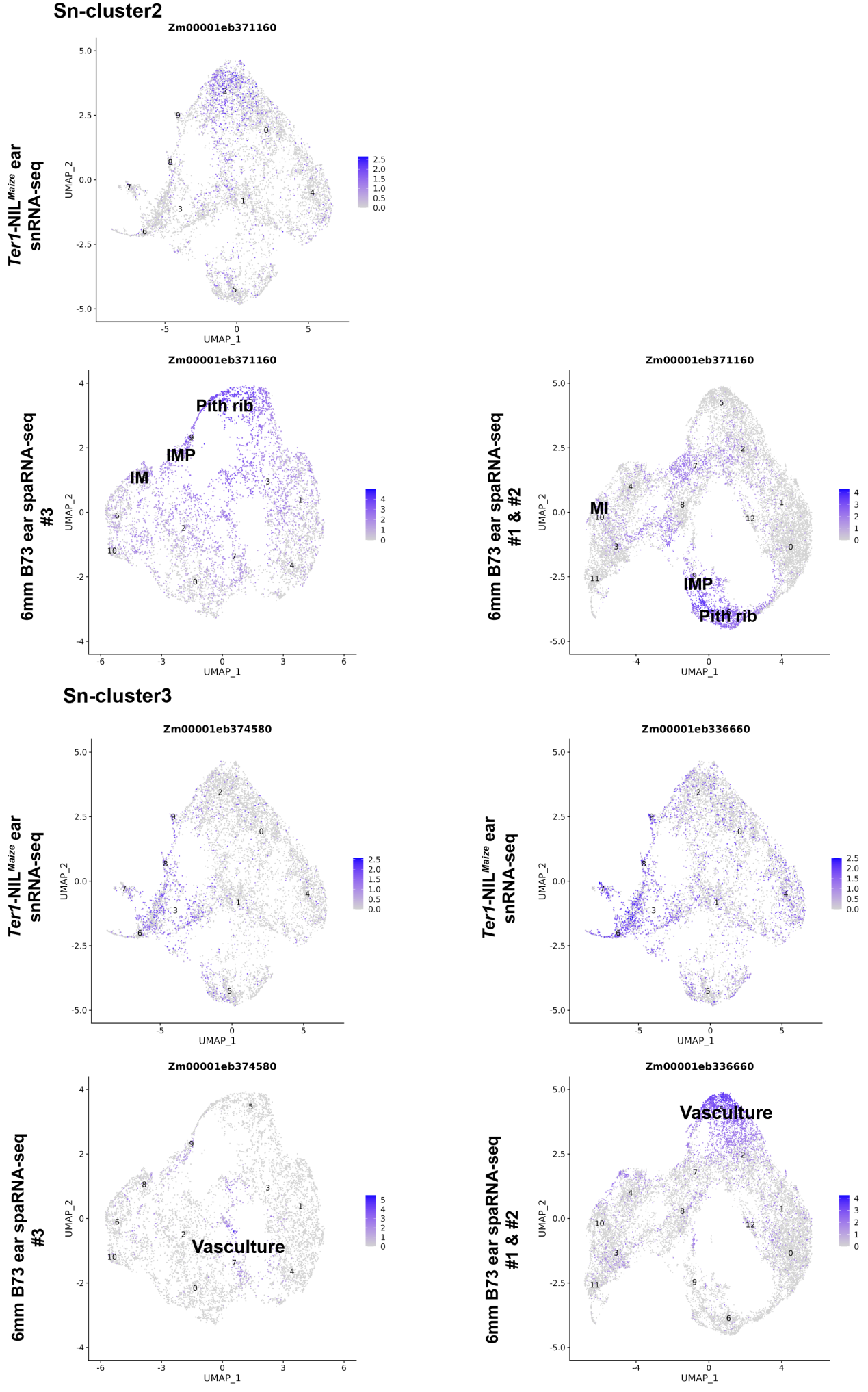

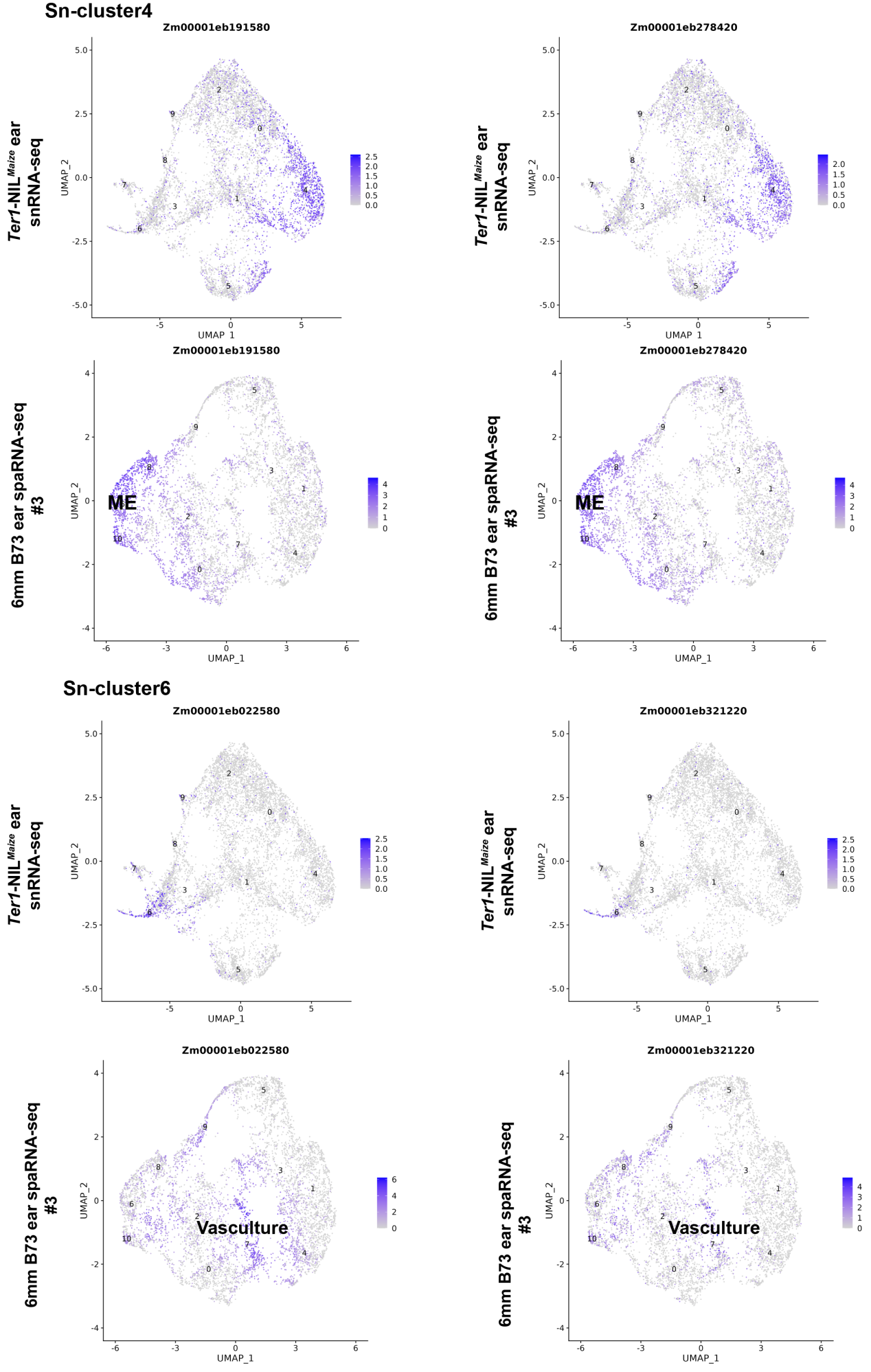

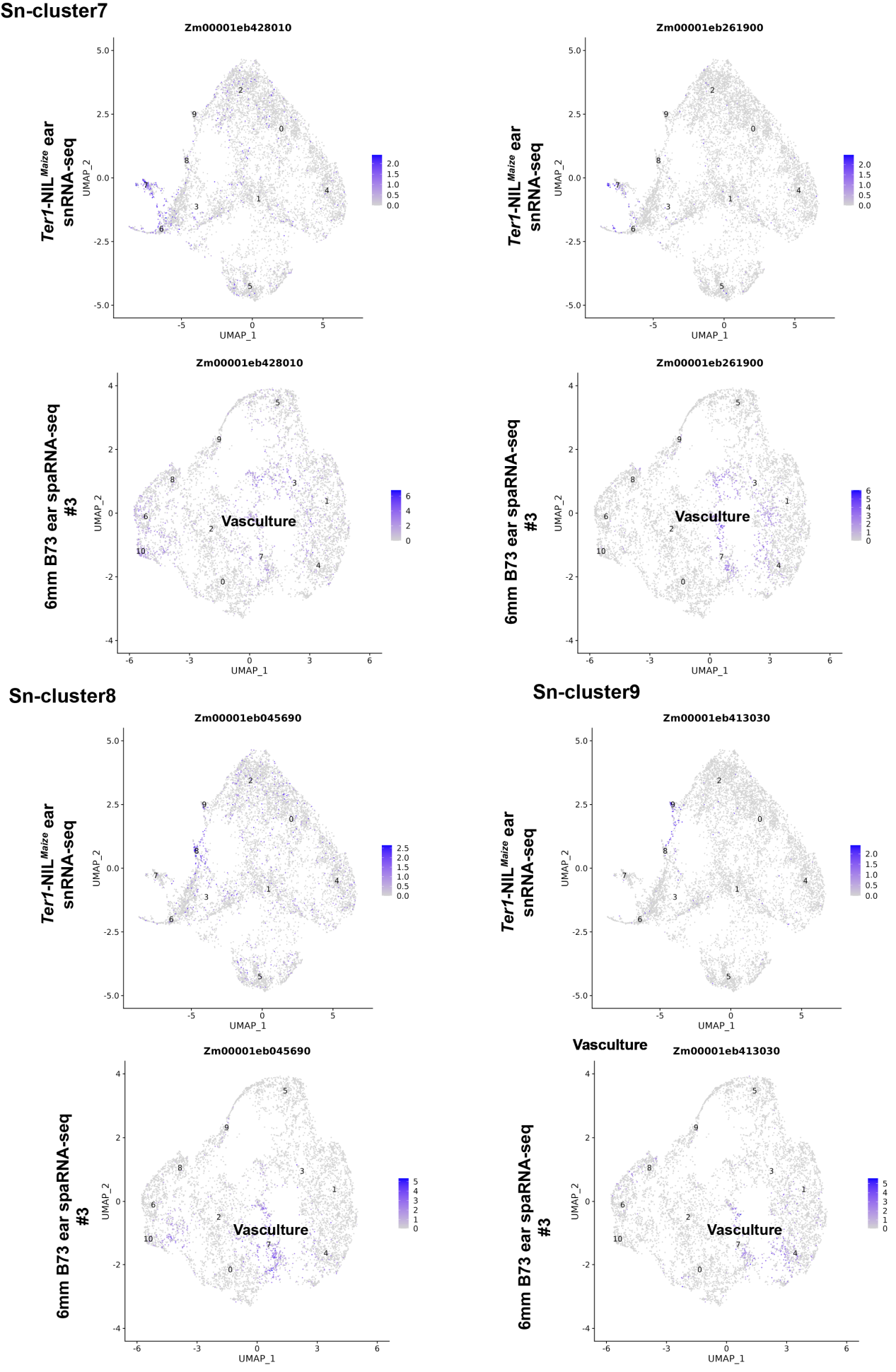

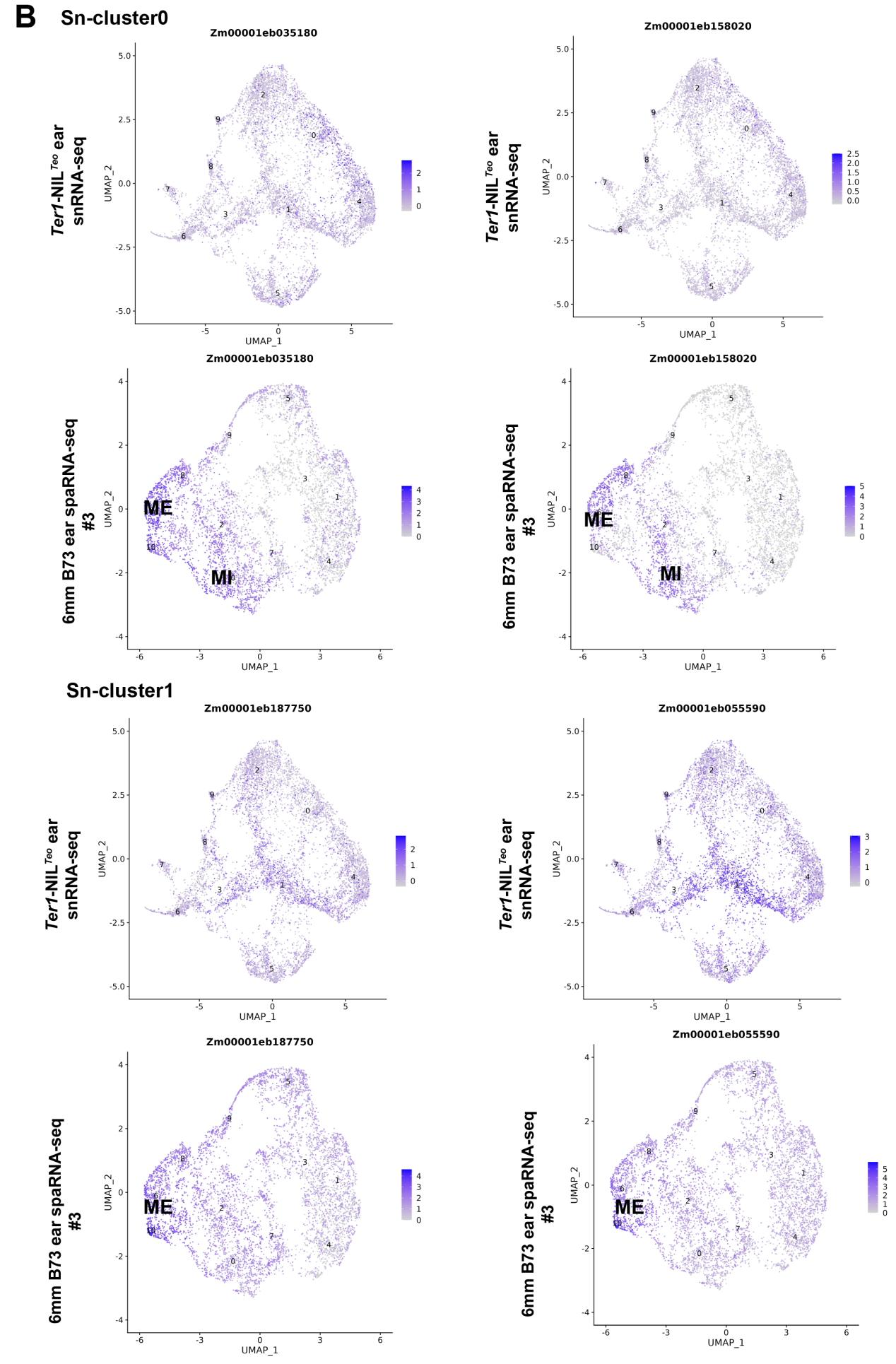

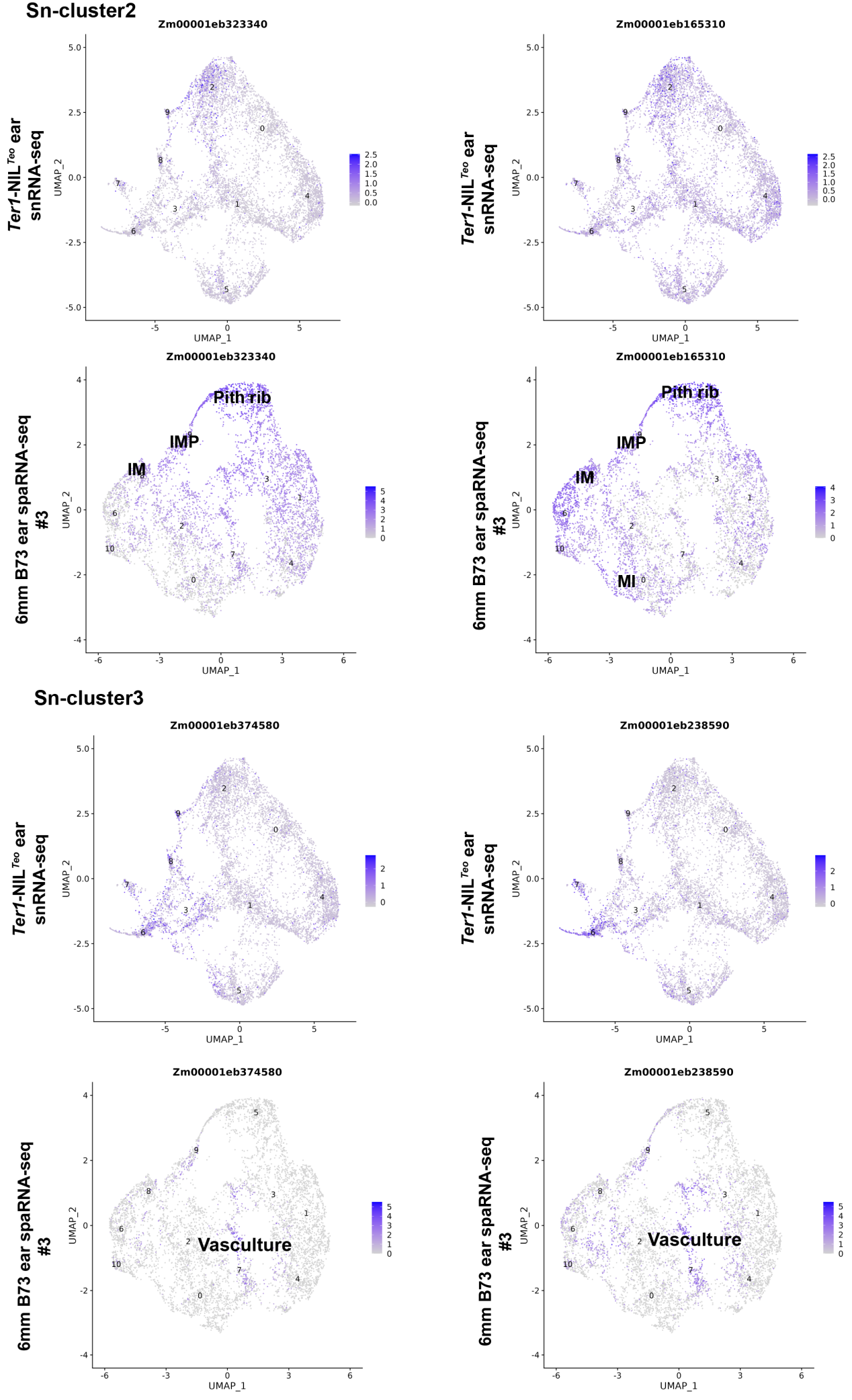

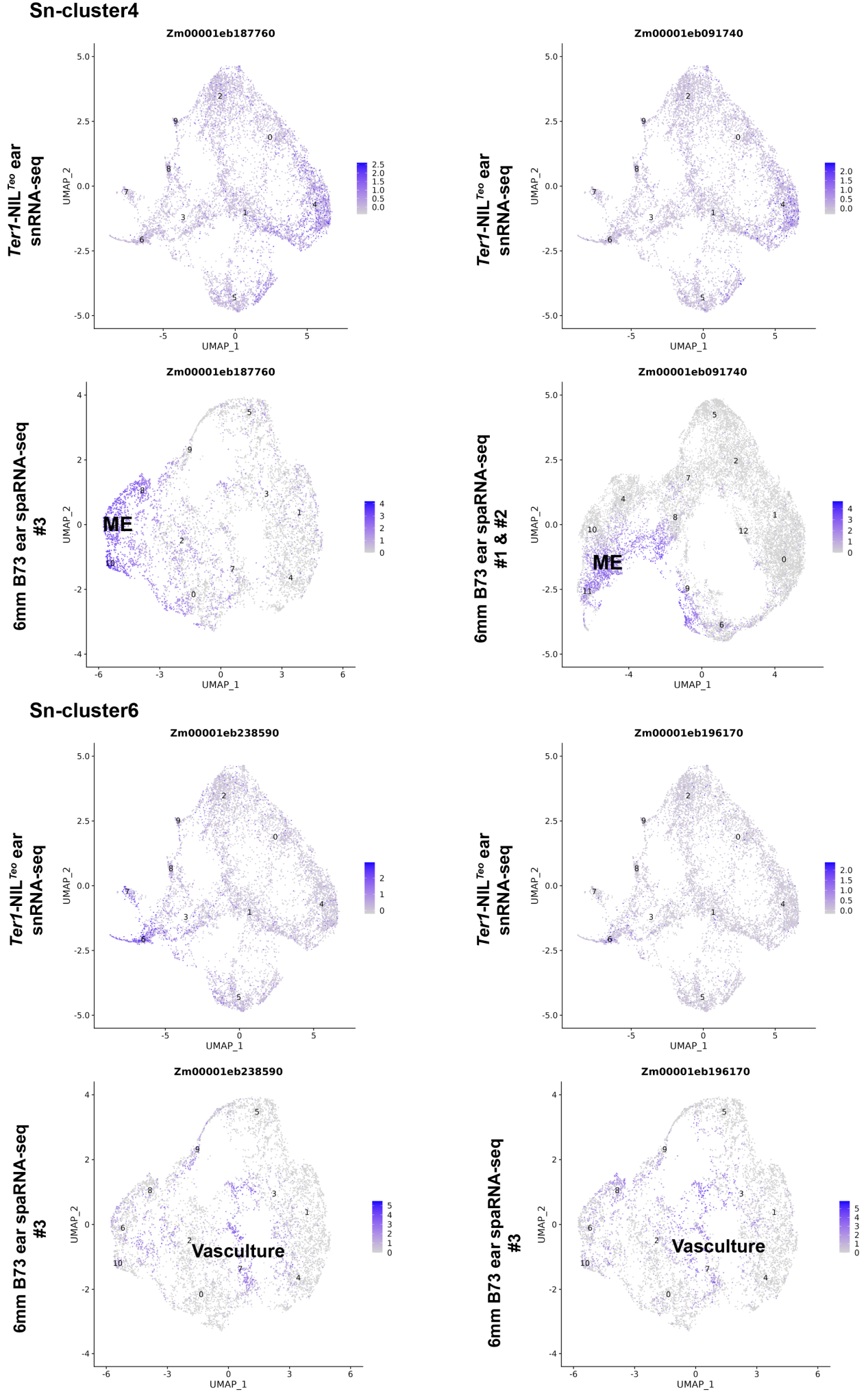

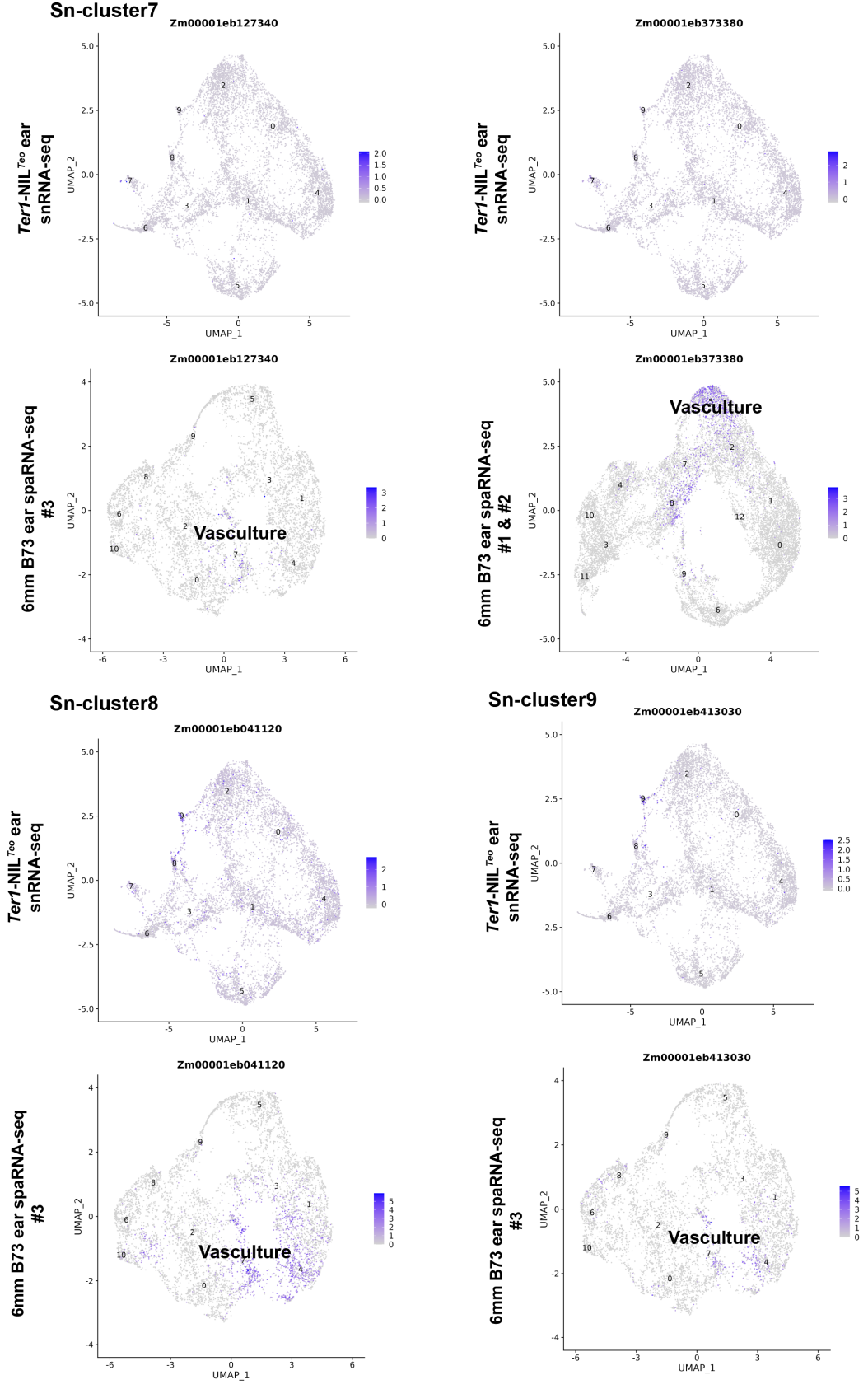

**Fig. S17 UMAP plots of marker genes predicting the identities of *Ter1*-NIL*^Teo^* and *Ter1*-NIL*^Maize^* Sn-Clusters. (A-B)** *Ter1*-NIL*^Maize^* (A) and *Ter1*-NIL*^Teosinte^* (B) snRNA-seq represent UMAP plots of their marker genes, the numbers in the figures indicate that the gene is the marker gene of the corresponding Sn-Cluster, 6mm B73 ear spaRNA-seq #1& #2 represent UMAP plots of the same genes in the published maize spatial transcriptome map shown in fig. S2A-B, 6mm B73 ear spaRNA-seq #3 represent UMAP plots of the same genes in the published maize spatial transcriptome map in Fig. S2C-D, with a color scale indicating normalized expression level.

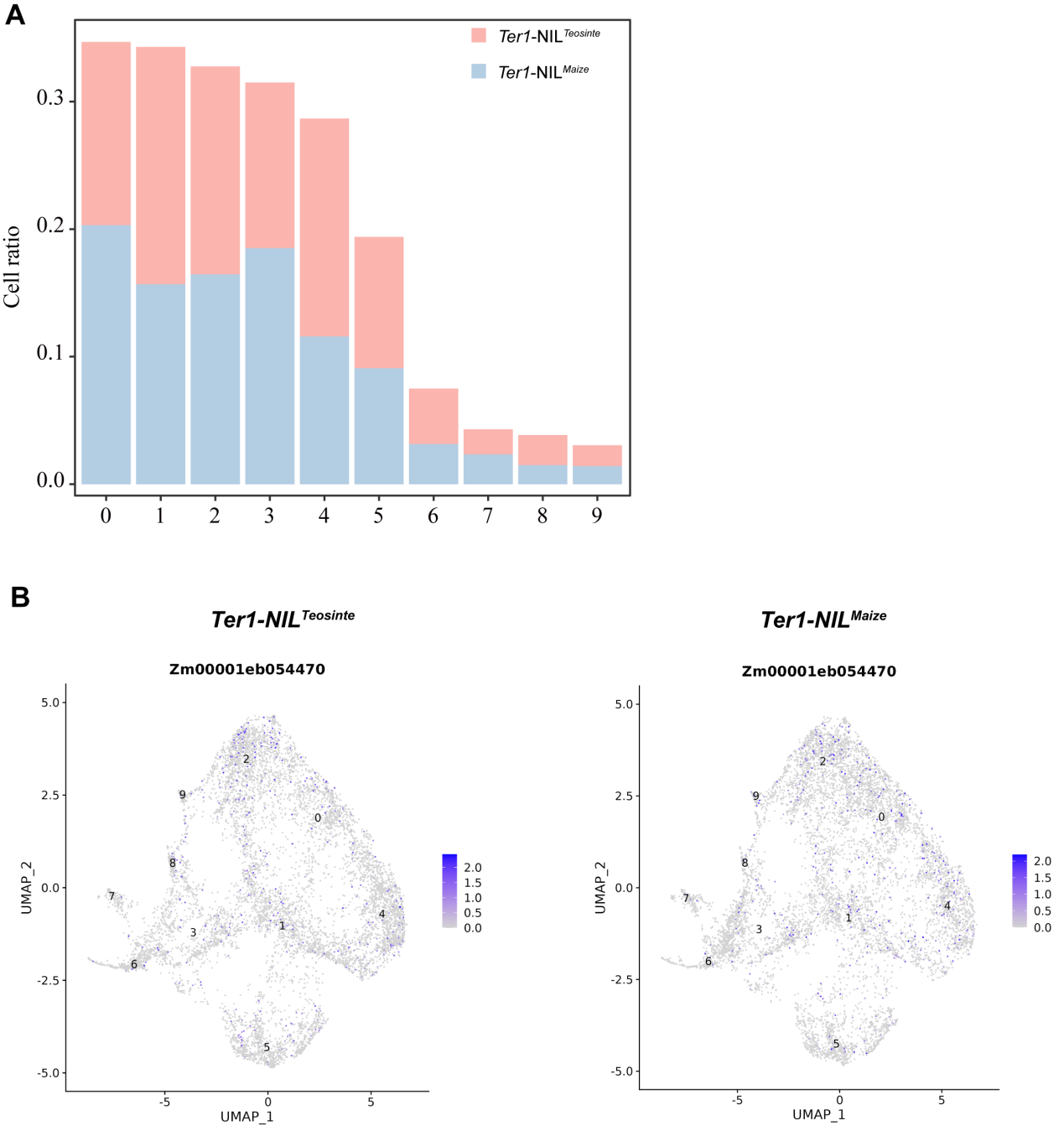

**Fig. S18 Integration of *Ter1*-NIL*^Teosinte^* and *Ter1*-NIL*^Maize^* snRNA-seq.** (A) The proportions of cells from *Ter1*-NIL*^Teosinte^* and *Ter1*-NIL*^Maize^* in each cell cluster, respectively. (B) Expression patterns of *Ter1* candidate gene *Zm00001eb054470* in *Ter1*-NIL*^Teosinte^* and *Ter1*-NIL*^Maize^* snRNA-seq data.

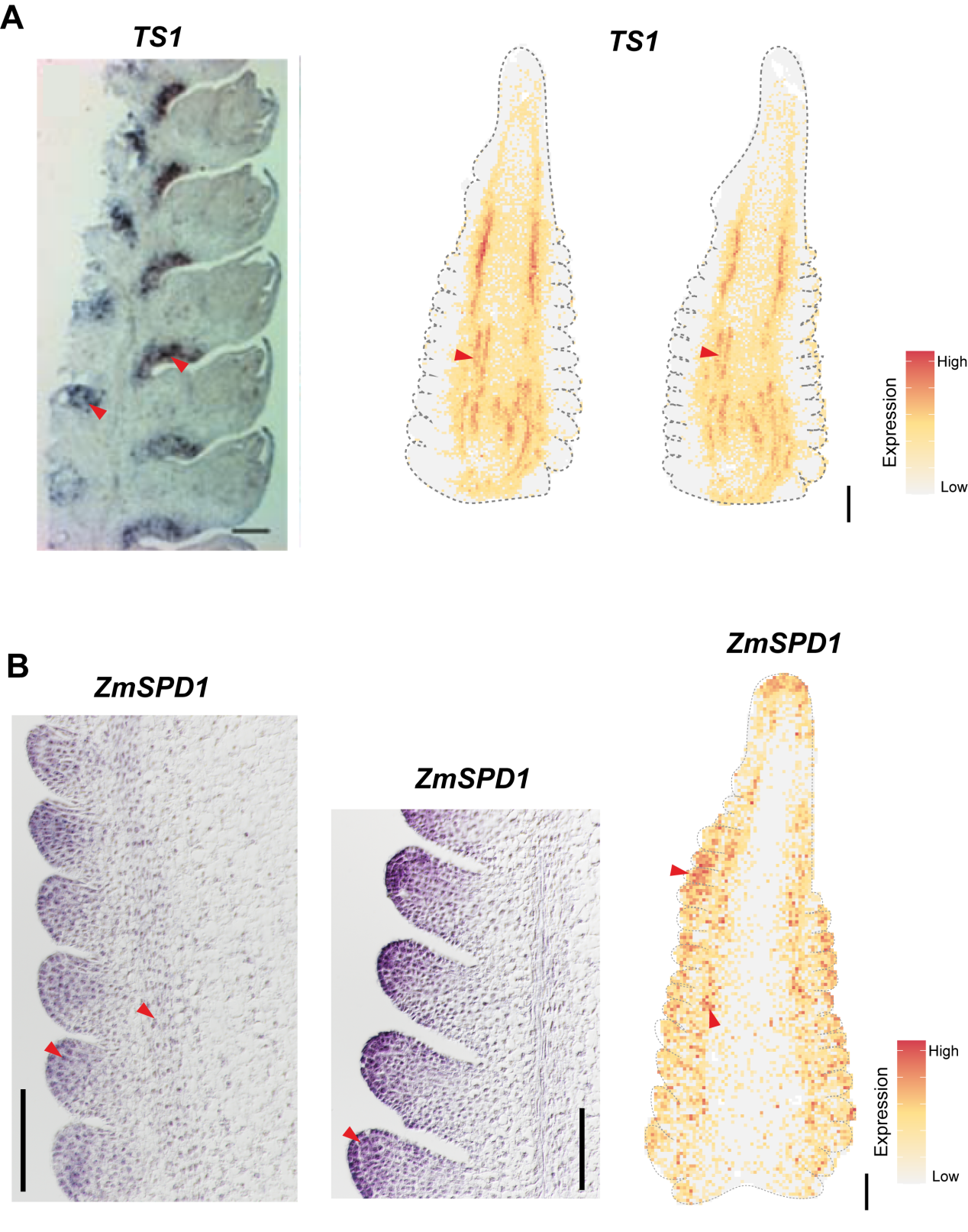
**Fig. S19 mRNA *in situ* hybridization of *TS1* and *ZmSPD1* in maize.** (A) Left, *TS1* mRNA *in situ* hybridization result from (*5*). Right, Stereo-seq expression pattern of maize 6mm ear, scale bar = 0.5 mm. (B) Left, *ZmSPD1* mRNA *in situ* hybridization result in maize, scale bar = 0.1 mm. Right, Stereo-seq expression pattern of maize 6mm ear, scale bar = 0.5 mm.

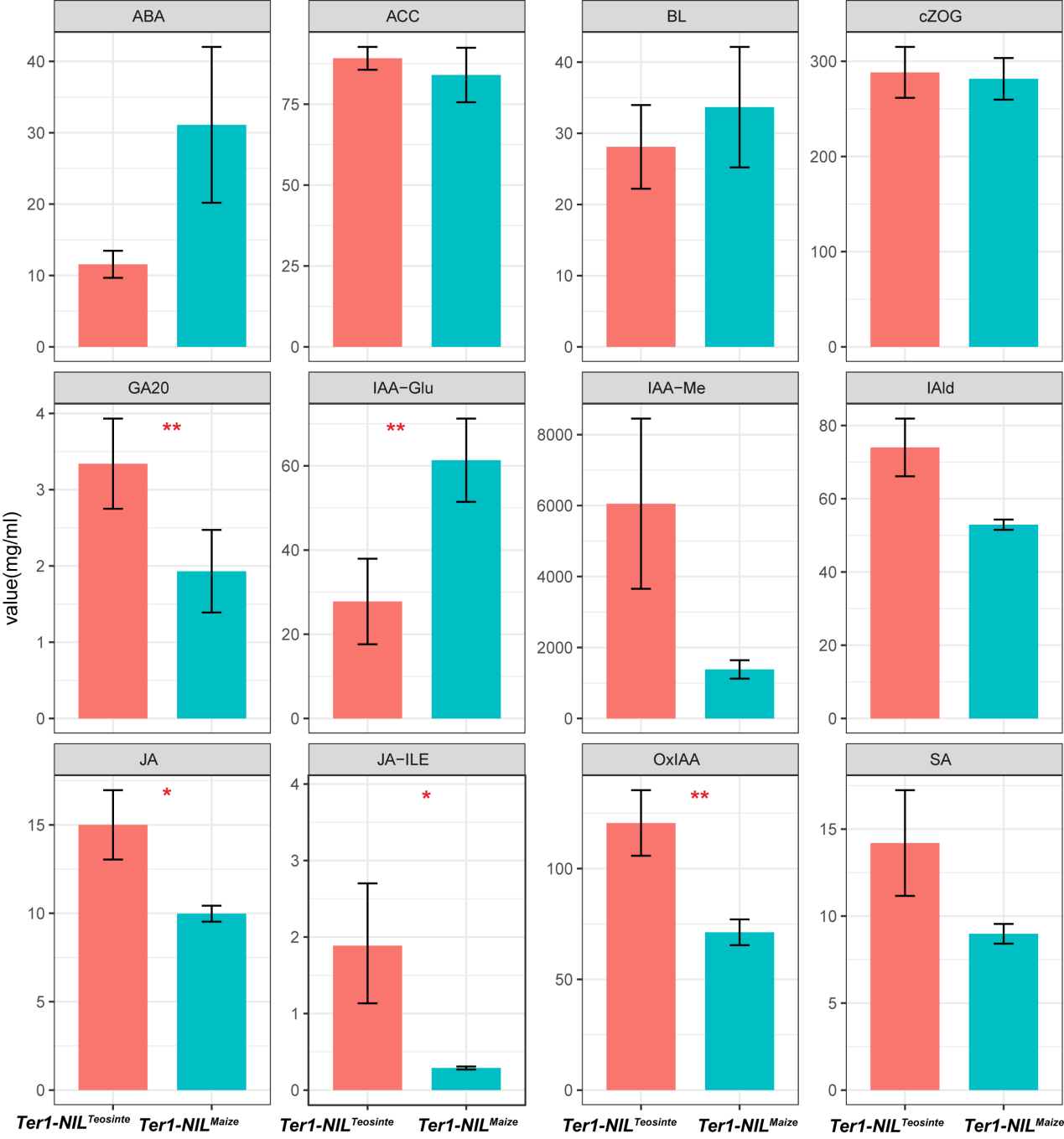

**Fig. S20 Quantification of endogenous hormone levels in *Ter1*-NIL*^Teosinte^* and *Ter1*-NIL*^Maize^* ears.** Abscisic acid (ABA); 1-Aminocyclopropanecarboxylic acid (*4*); Brassinolide (BL);cis-Zeatin-O-glucoside (cZOG); Gibberellin A20 (GA20); Indole-3-acetyl-glutamate (IAA-Glu); Indole-3-acetic acid methyl ester (IAA-Me); Indole-3-carboxaldehyde (IAld); Jasmonic acid (JA); Jasmonic acid-isoleucine (JA-ILE); 2-Oxoindole-3-acetic acid (OxIAA); Salicylic acid (SA). Error bars indicate SD (n = 3). Asterisks indicate significant differences (Student's t-test: *P < 0.05, **P < 0.01).

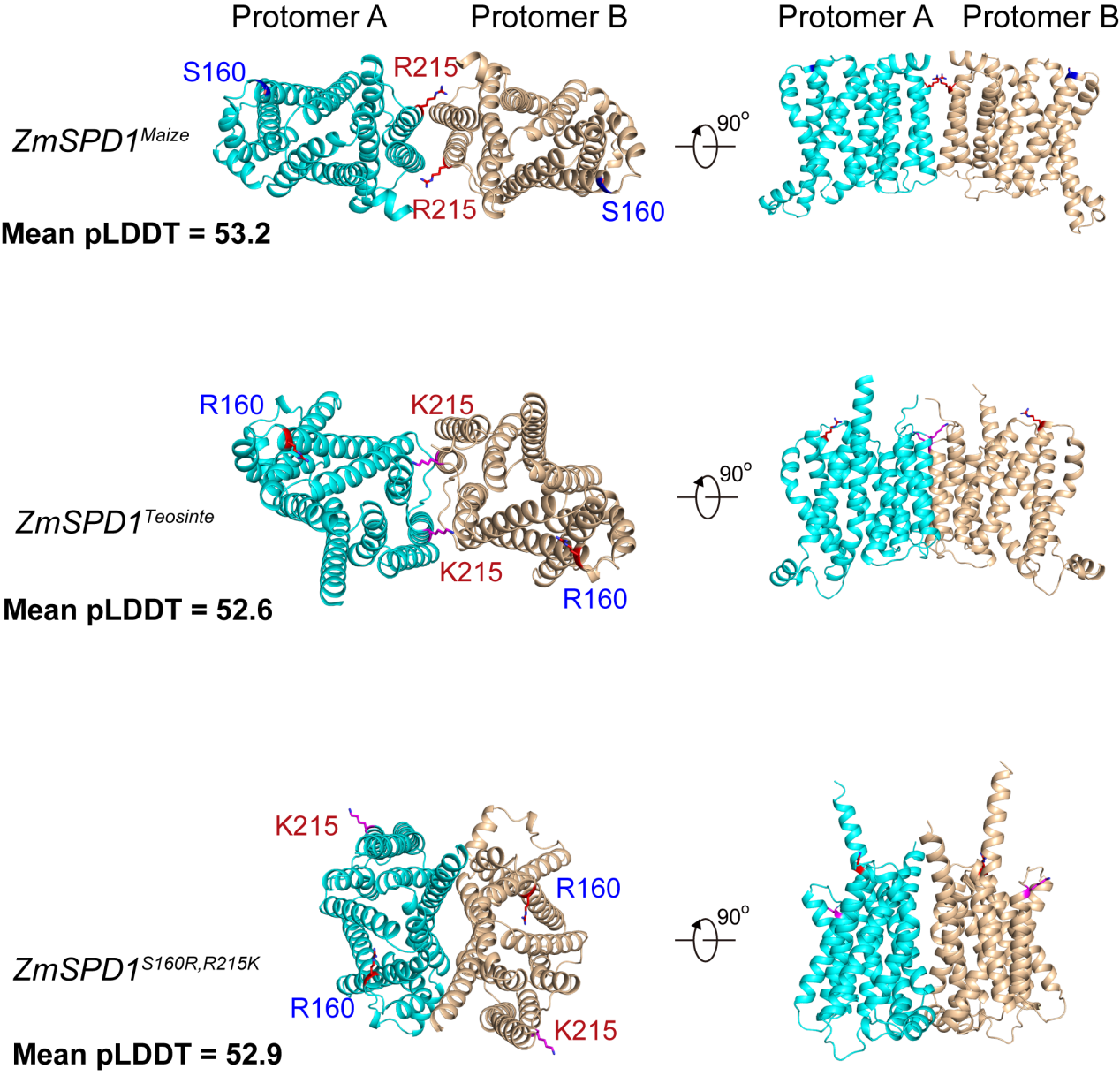

**Fig. S21 Protein structural comparison of different alleles of *ZmSPD1*.** The *ZmSPD1^Maize^* allele is based on the DNA and corresponding protein sequences derived from the B73 reference genome. The *ZmSPD1^Teosinte^* allele is based on sequences from *Zea mays ssp. parviglumis* (TIL01). The *ZmSPD1^S160R,R215K^* variant represents a modified version of the B73 *ZmSPD1* sequence, in which two nonsynonymous mutations were replaced with the teosinte-type alleles, and the corresponding DNA and protein sequences were used for analysis. pLDDT (predicted local distance difference test) scores indicate per-residue confidence in the predicted structure, ranging from 0 to 100, with higher scores representing higher confidence

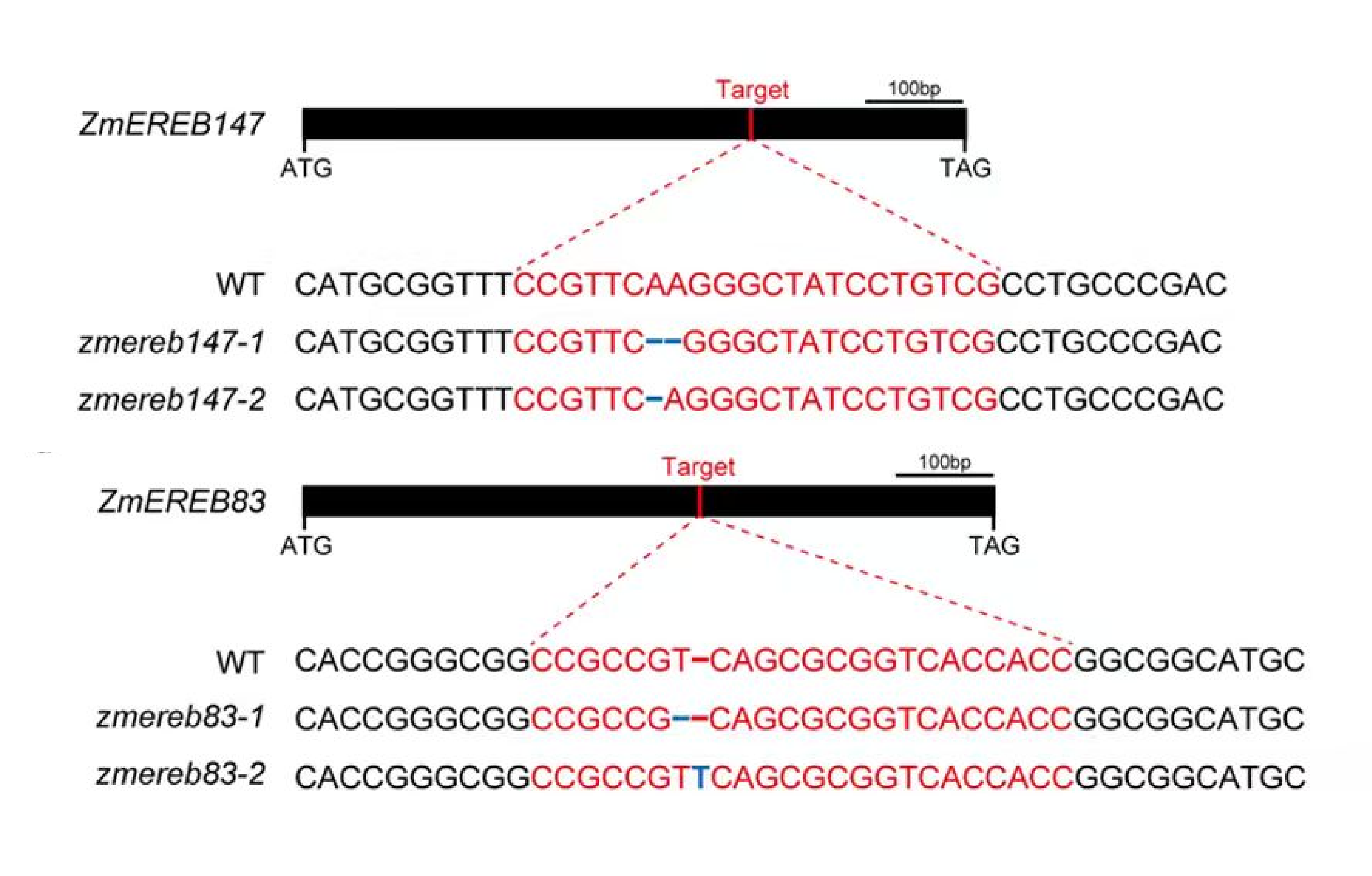

**Fig. S22 Genotype of *ZmEREB83* and *ZmEREB147* CRISPR-Cas9 edited plants.** The red lines denote sgRNA, and the black boxes indicate the coding sequence.

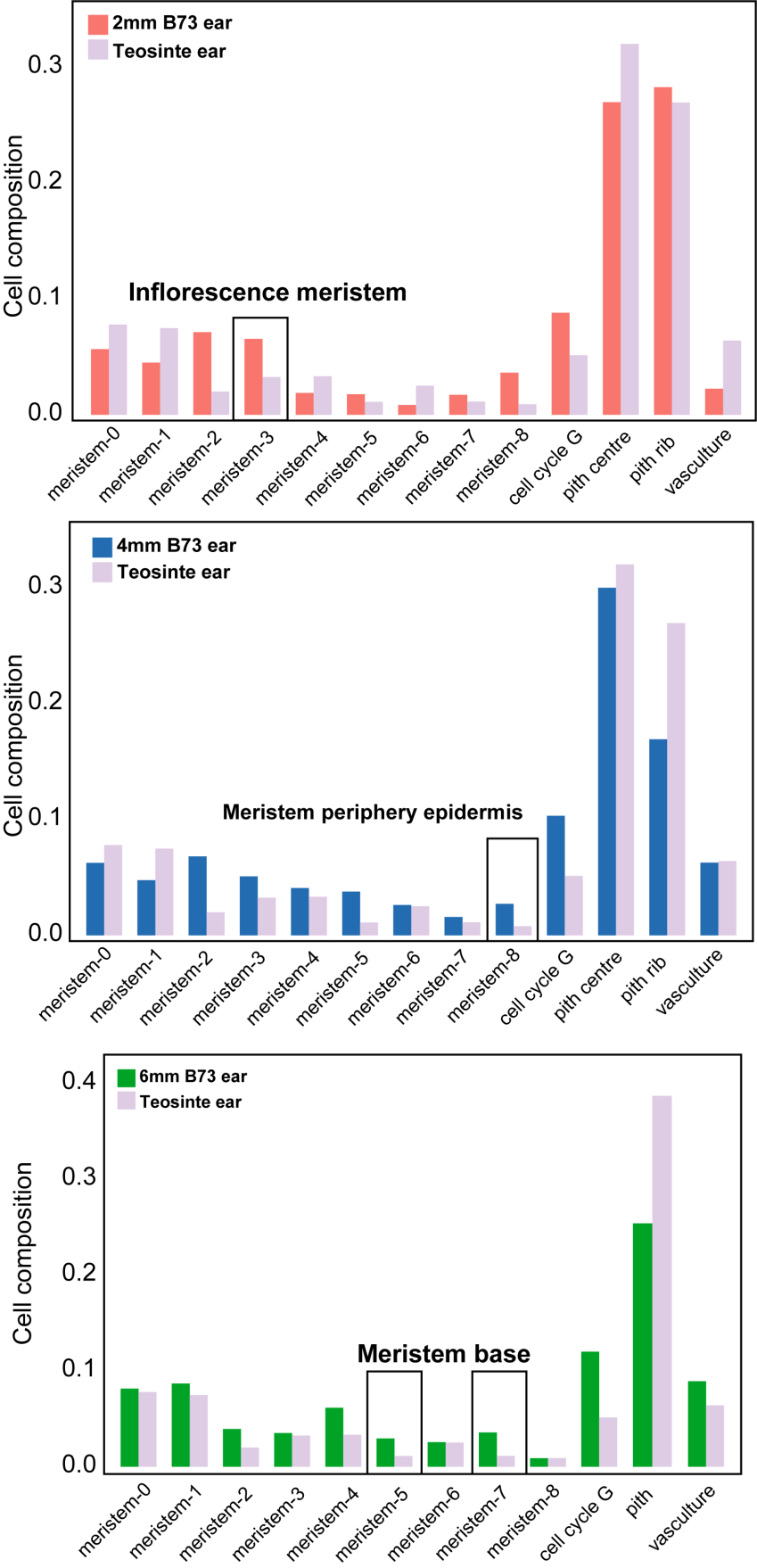

**Fig. S23 Comparison of cellular composition between maize and teosinte.** Meristem-0 to meristem-8 present cell types from spikelet meristem cells, and are shown in Fig. 2F. Other cell types from the maize ear were shown in Fig. 1B, and cell types of the teosinte ear were shown in Fig. 1E. The black boxes indicate cell types with a more than two-fold difference in proportion.

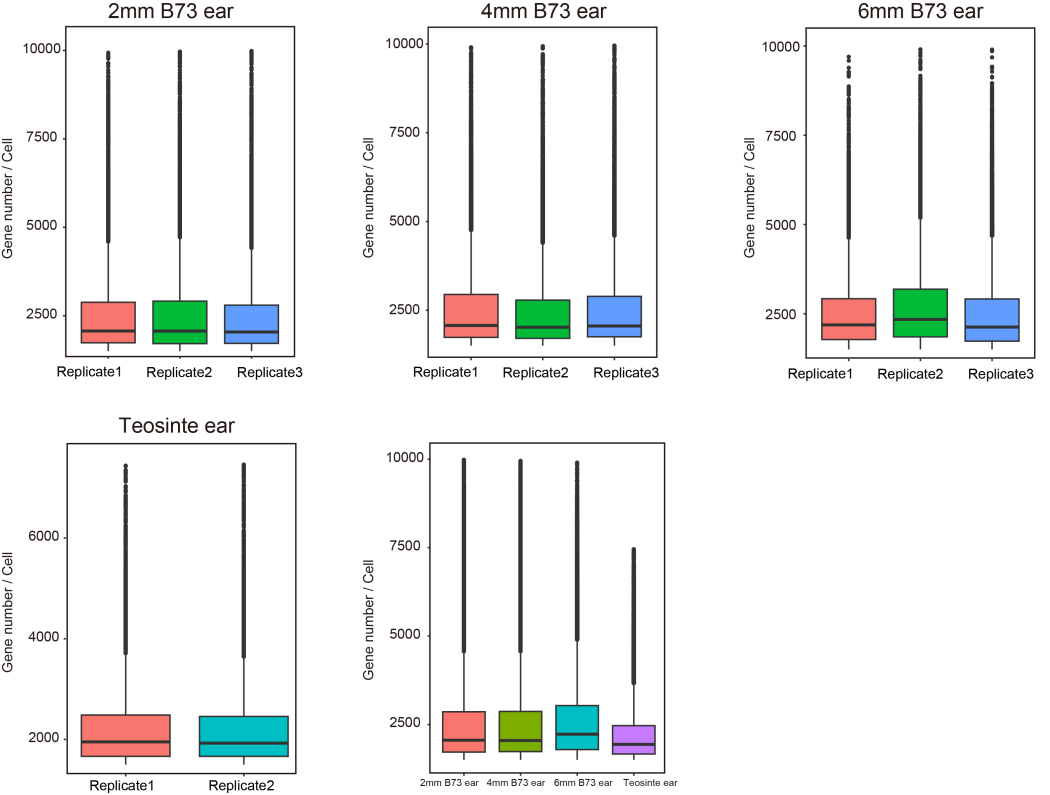

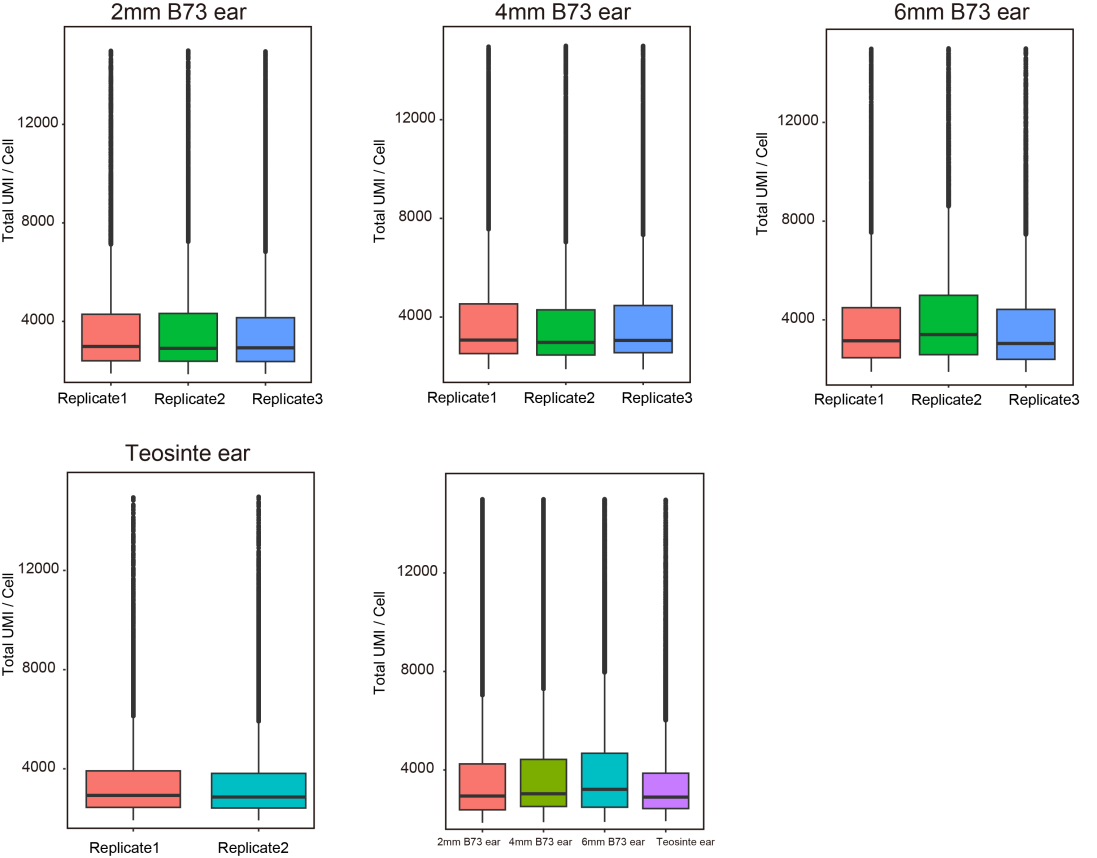

**Fig. S24 Box plots of the number of total UMI/cell and gene number/cell of maize and teosinte snRNA-seq.**

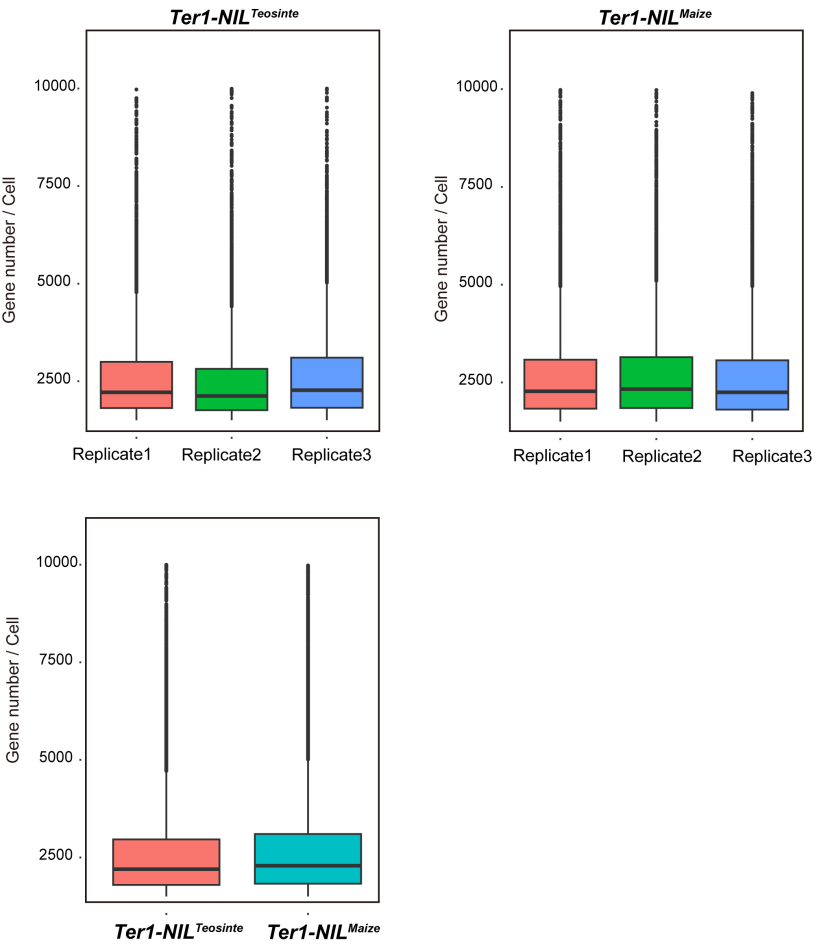

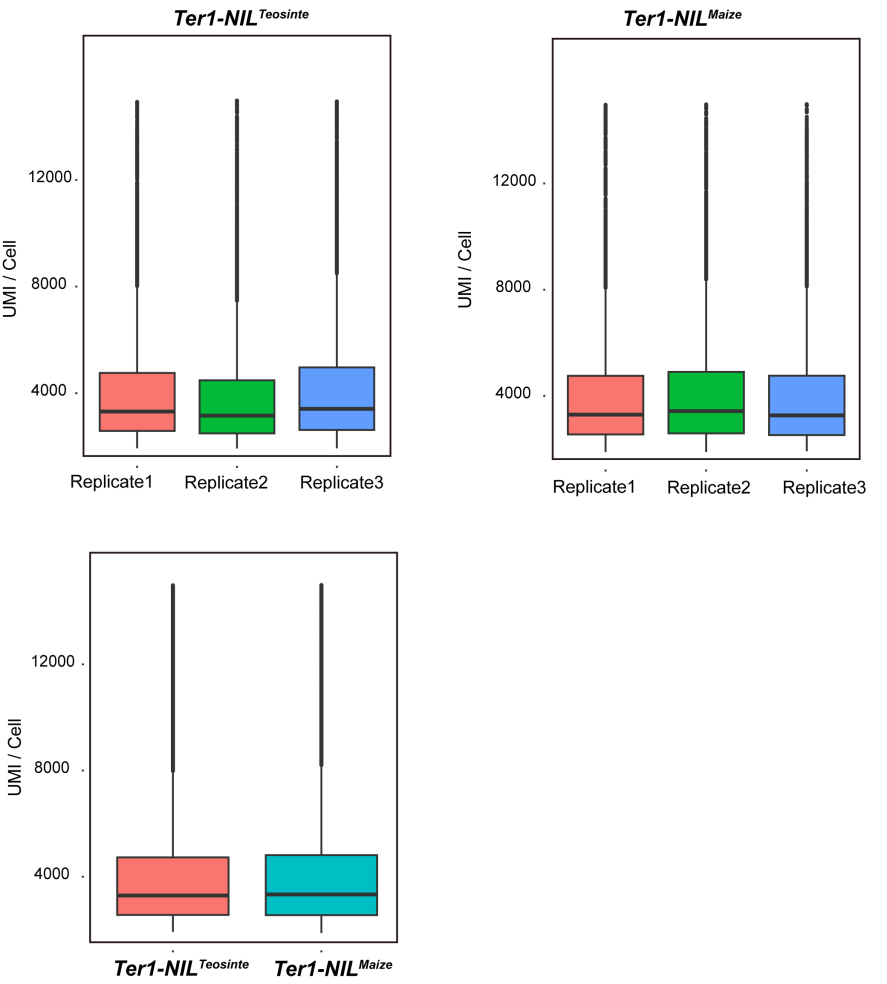

**Fig. S25 Box plots of the number of total UMI/cell and gene number/cell of *Ter1*-NIL*^Teosinte^* and *Ter1*-NIL*^Maize^* snRNA-seq.**

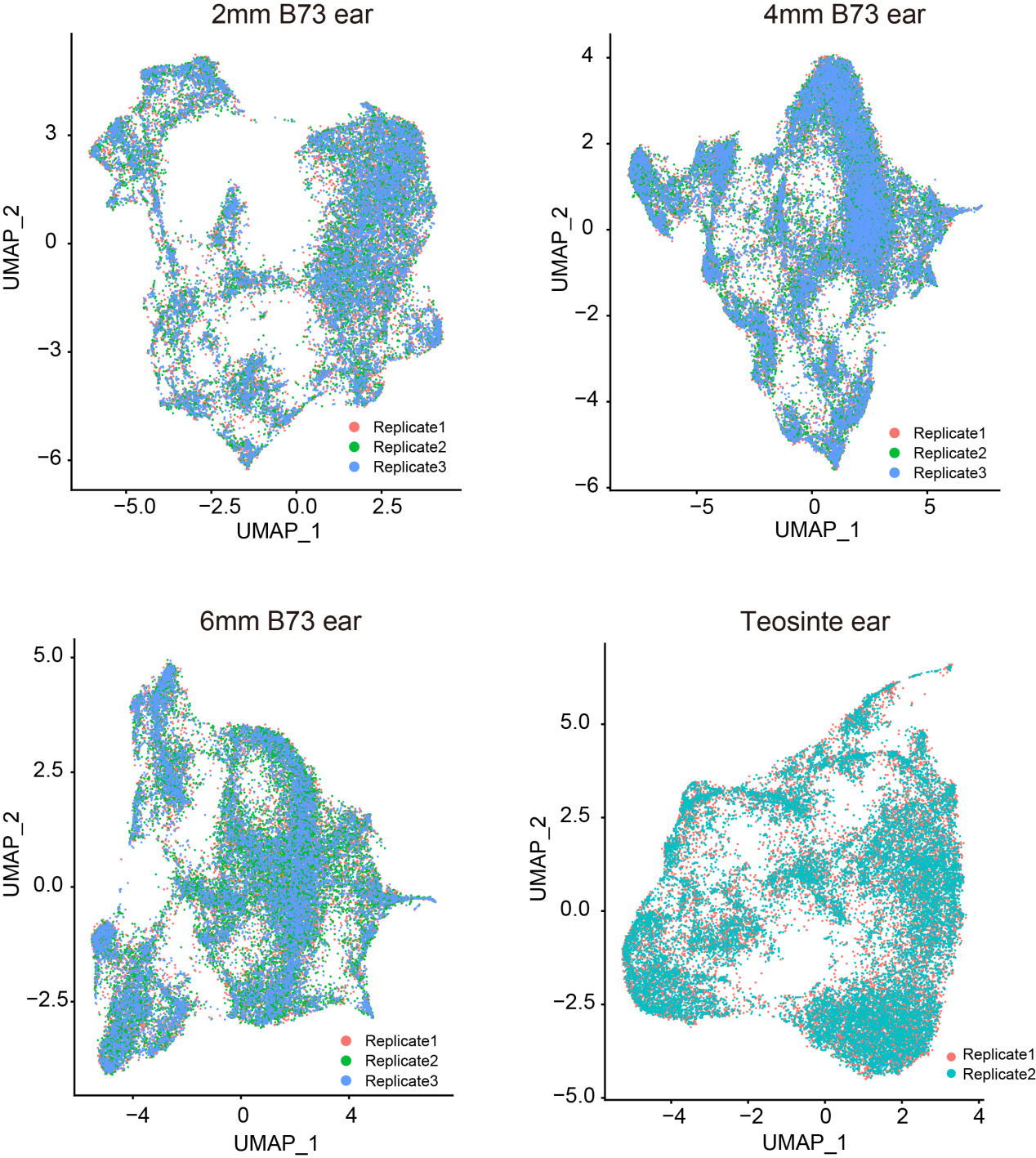

**Fig. S26 UMAP plot showing good reproducibility of maize and teosinte snRNA-seq**

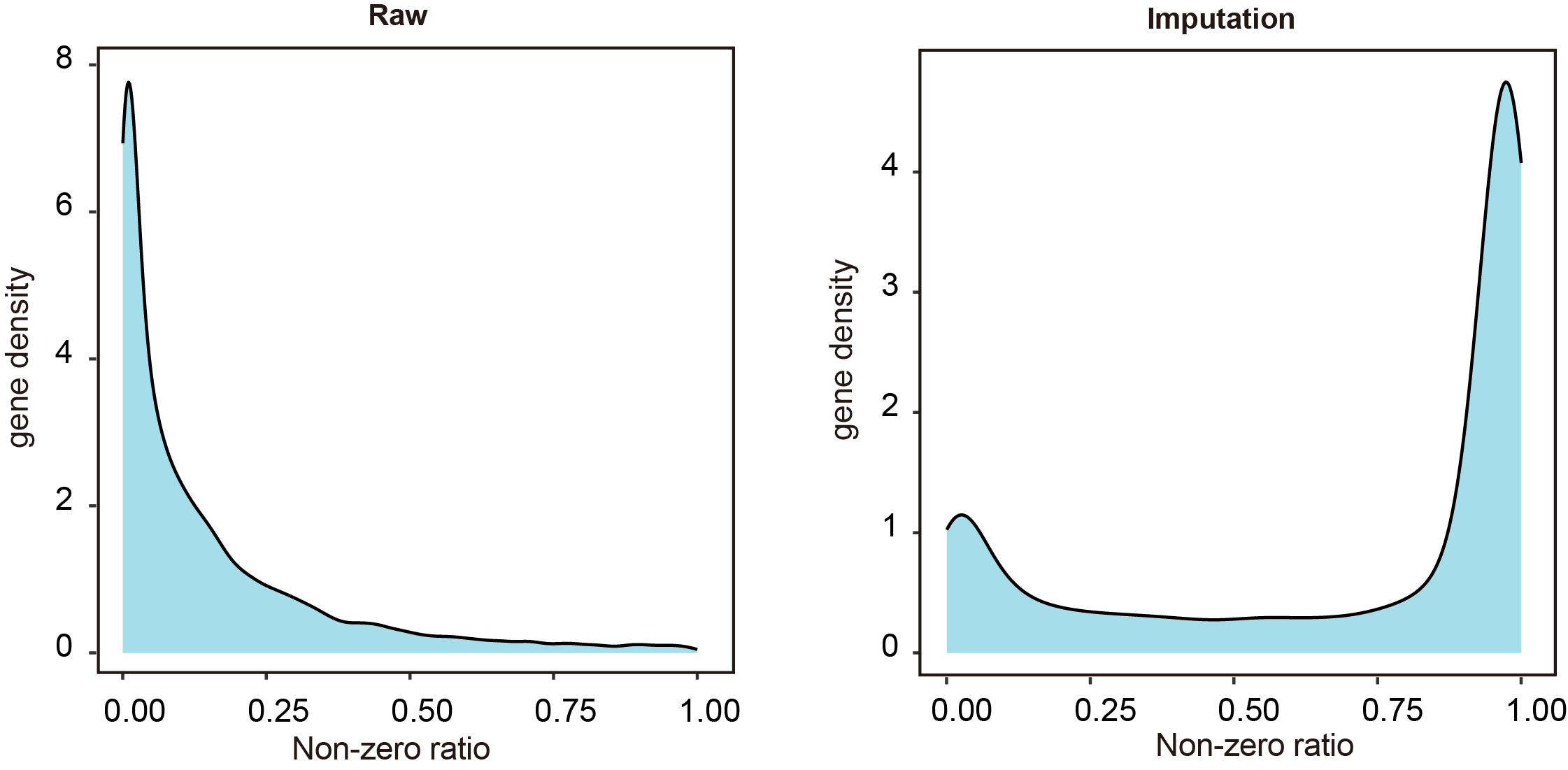

**Fig. S27 Gene density of non-zero ratio across three teosinte ear sections before and after imputation.** Gene density of non-zero ratio across three teosinte sections before and after imputation

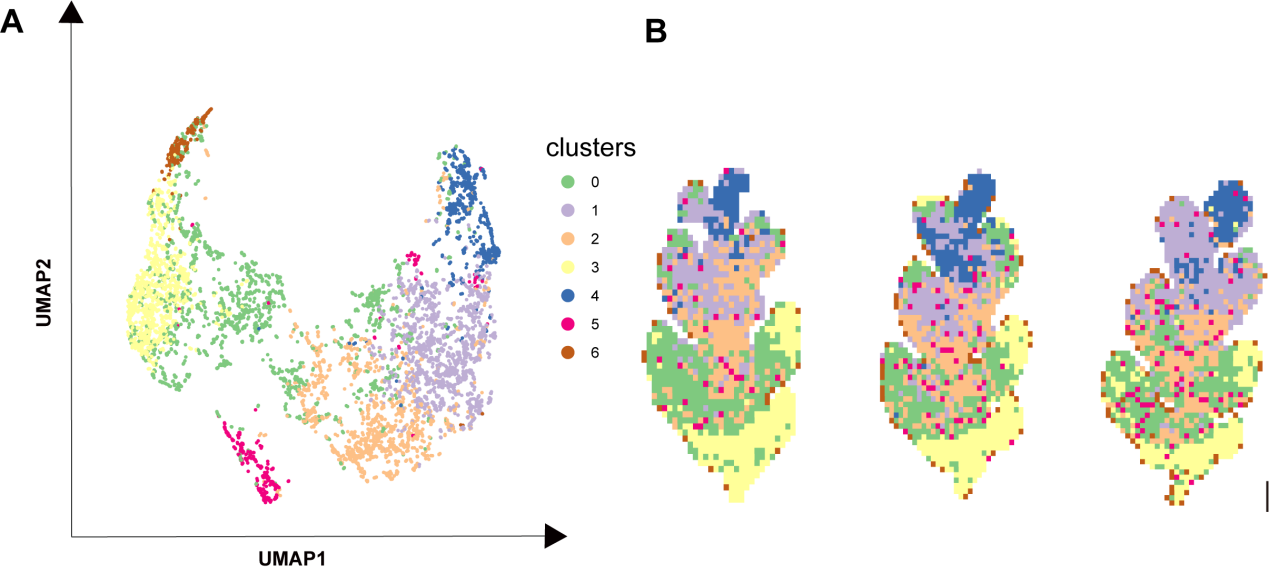

**Fig. S28 Unsupervised clustering of three teosinte sections based on raw Stereo-seq data.** (A) **7** meta-clusters displayed by an integrated uniform manifold approximation and projection (UMAP) plot in two dimensions, with each dot representing a bin50. (B) Physical distribution of bin50s corresponding to different clusters. Scale bar = 0.1 mm.

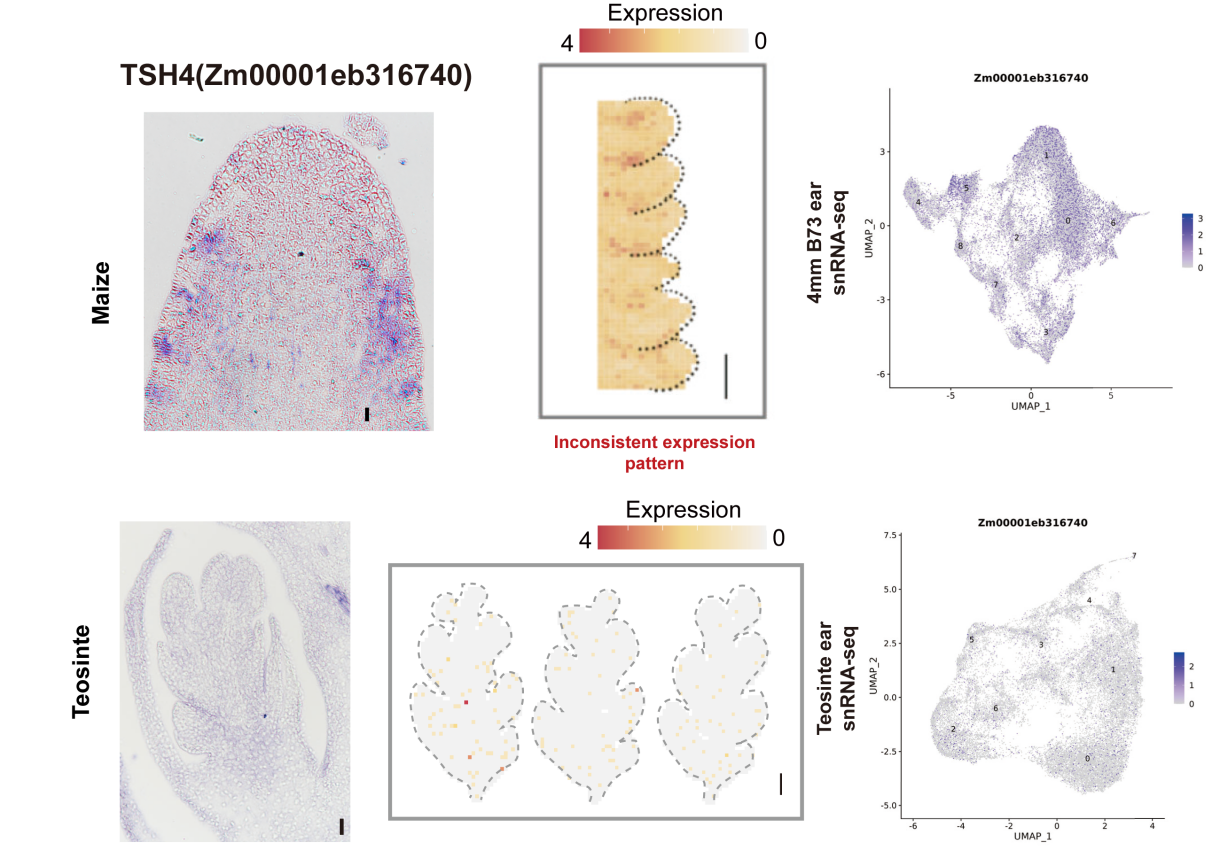

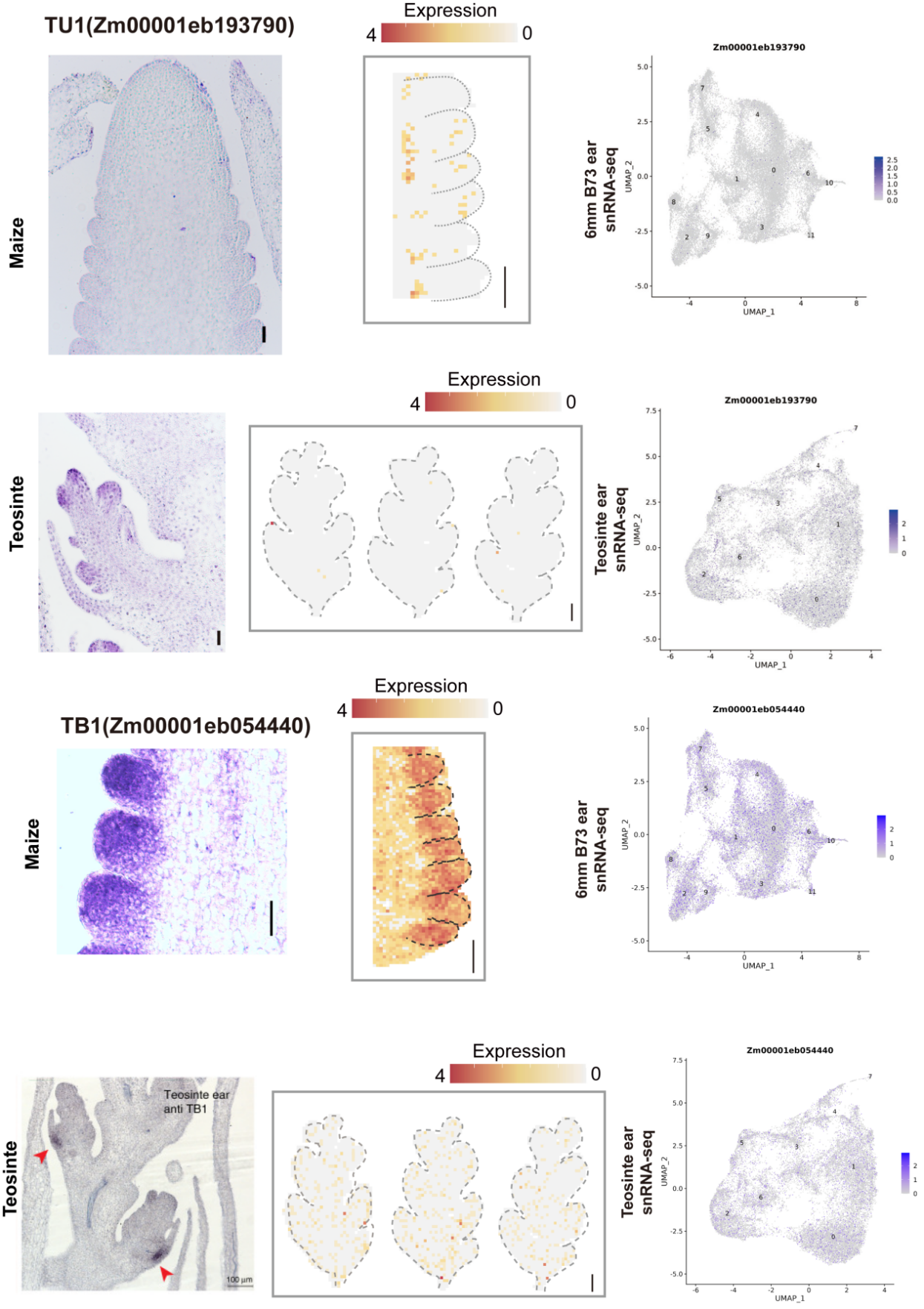

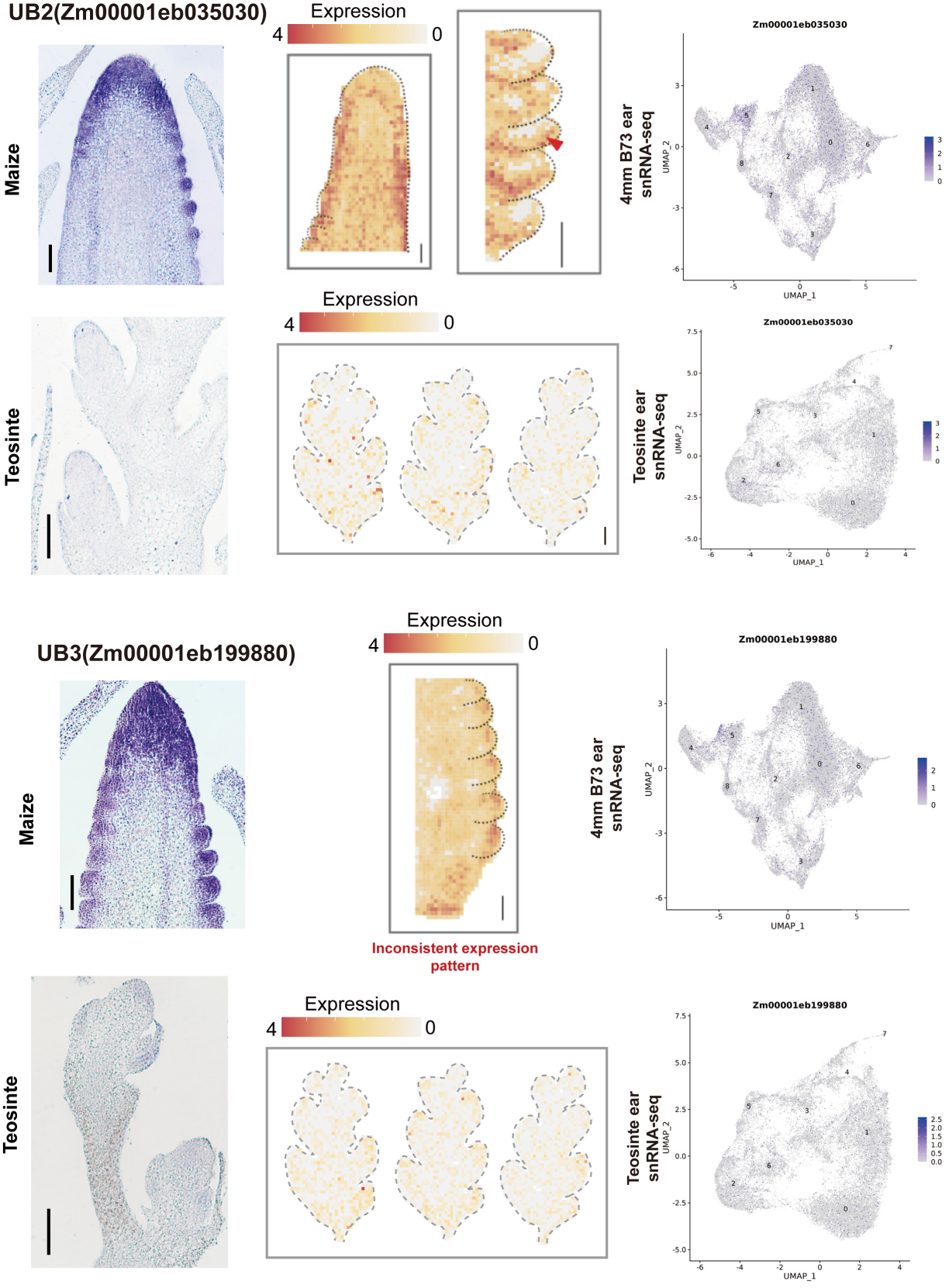

**Fig. S29** mRNA *in situ* hybridization of known genes exhibits different expression patterns in maize and teosinte. Left, maize mRNA *in situ* hybridization result. Middle, Stereo-seq expression pattern of teosinte ear, scale bar = 0.1 mm. Right, Teosinte snRNA-seq expression pattern. Color scales indicate normalized expression levels. Known gene expression pattern were validated from *TSH4* (*6*), *UB2* (*7*), *UB3* (*6*), *TB1*(*8, 9*).
